# The integration of Tgfβ and Egfr signaling programs confers the ability to lead heterogeneous collective invasion

**DOI:** 10.1101/2020.11.14.383232

**Authors:** Apsra Nasir, Sharon Camacho, Alec T. McIntosh, Garrett T. Graham, Raneen Rahhal, Molly E. Huysman, Fahda Alsharief, Anna T. Riegel, Gray W. Pearson

**Author notes:** To whom correspondence should be addressed., Gray W. Pearson, Lombardi Comprehensive Cancer Center, Georgetown University, 3970 Reservoir Road NW, Washington, DC 20057. Equal contribution (Apsra Nasir, Sharon Camacho).

## Abstract

Phenotypic heterogeneity promotes tumor evolution and confounds treatment. Minority subpopulations of trailblazer cells enhance the heterogeneity of invading populations by creating paths in extracellular matrix (ECM) that permit the invasion of phenotypically diverse siblings. The regulatory programs that induce a trailblazer state are poorly understood. Here, we define a new Tgfβ induced trailblazer population that is more aggressive than previously characterized Keratin 14 expressing trailblazer cells. Rather than triggering a binary switch to a single trailblazer state, Tgfβ induced multiple unique states that were distinguished by their expression of regulatory transcription factors, genes involved in ECM reorganization and capacity to initiate collective invasion. The integration of a parallel Egfr signaling program was necessary to induce pro-motility genes and could be targeted with clinically approved drugs to prevent trailblazer invasion. Surprisingly, Egfr pathway activity also had the collateral consequence of antagonizing the expression of a cohort of Tgfβ induced genes, including a subset involved in ECM remodeling. Together, our results reveal a new compromise mode of signal integration that promotes a trailblazer state and can be therapeutically targeted to prevent collective invasion.

## INTRODUCTION

Tumor cell phenotypic heterogeneity is the product of genetic and epigenetic variability, localized differences in the microenvironment and stochastic fluctuations in intracellular signaling pathways (1). This diversity in cell phenotypes drives tumor evolution (2). Cells with traits that provide a competitive advantage over siblings become predominant, improving the fitness of the population (3). High variance in cell phenotypes also increases the odds that a subpopulation has the potential to survive when the local environment changes, such as during treatment or when disseminating cells seed other organs (4, 5). Interestingly, analysis of intrinsic and genetically engineered heterogeneity has revealed that phenotypic diversity also promotes functional interactions between tumor cells (6, 7). Rare enabler tumor subpopulations can promote the growth, immune evasion, invasion and metastasis of abundant siblings harboring different phenotypic traits (8-13). However, the regulatory programs that promote functional relationships between phenotypically distinct subpopulations remain poorly understood. This lack of mechanistic insight limits our ability to define the presence of functional tumor cell interactions, evaluate their impact on patient outcome, and develop new treatment options designed to target rare enabler cells (14).

To better understand how subpopulation relationships are induced during tumor progression, we sought to understand how tumor cells are conferred with the ability to promote the collective invasion of intrinsically less invasive siblings. Collective invasion is the predominant mode of invasion in carcinomas and promotes the dispersion of tumor clusters that seed metastases (15-19). Subpopulations of trailblazer cells detected in breast, lung and thyroid tumor populations have an enhanced ability to reorganize the extracellular matrix (ECM) into micro-tracks that facilitate the initiation of collective invasion (20-23). Importantly, sibling opportunist populations invade through the micro-tracks created by minority trailblazer populations in three-dimensional culture models and xenograft tumors (21, 24, 25). Moreover, opportunist cells can have greater proliferative capacity than trailblazers in vitro and in xenografts, indicating that trailblazer-opportunist relationships have the potential to increase the proliferative fitness of invading cell populations (24, 25). Thus, trailblazer cells promote phenotypic diversity in invading cell cohorts, which may increase the risk of metastasis and elevate the risk of resistance to treatment (26, 27).

Trailblazer cells are distinguished from the bulk tumor population by the increased expression of trailblazer genes, including regulators of the actin cytoskeleton, integrins, receptor tyrosine kinases, keratins, and cancer testes antigens (20, 21, 23, 24, 28). Trailblazer genes contribute to the formation of actin-rich cellular protrusions that provide traction and exert tensile forces necessary for ECM remodeling and other undefined functions (21, 23, 25). Interestingly, trailblazer genes are not required for other forms of cell movement, including migration along two-dimensional surfaces, movement within spheroids and opportunistic collective invasion (21). While genes that specifically contribute to unique invasive characteristics of trailblazer cells have begun to be defined, the regulatory programs underpinning the conversion to a trailblazer state remain largely unknown. Trailblazer cells in breast and lung cancer cell line models are epigenetically distinct from opportunist siblings, although the processes that initiate the conversion to the trailblazer state are not known (21, 23). Breast cancer trailblazer cells have basal characteristics, with Keratin 14 (Krt14) expression proposed as a universal feature for identifying trailblazer cells in mammary tumors (18, 20, 29, 30). Yet, the program controlling the induction of a basal trailblazer state is undefined. Breast cancer trailblazer cells also have features of epithelial-mesenchymal transition (EMT) re-programming (21). However not all EMT programs yield trailblazer activity and whether EMT programs regulate the expression of trailblazer genes is not known (25, 31). Thus, if and how the EMT process imbues a trailblazer state is unclear. Our poor understanding of how trailblazer populations are induced limits options for potential interventions targeting trailblazer cells and advances in employing subpopulation identification to refine patient outcome predictions.

To define factors regulating trailblazer program activity, we determined how cells were induced to collectively invade in the C3(1)/SV40 T-antigen (C3-TAg) genetically engineered mouse model (GEMM) of breast cancer. Based on histological analysis, breast cancer is defined as Estrogen Receptor-positive (ER-positive), Human Epidermal Growth Factor Receptor-positive (HER2-pos) or triple-negative (TNBC), which have low to undetectable levels of ER, HER2 and the Progesterone Receptor (32). Breast cancer is further stratified into luminal A, luminal B, HER2-enriched and basal-like based on mRNA expression. Luminal A are Estrogen Receptor-positive (ER-pos) while luminal B are more proliferative and are ER-pos or HER2-pos (33). The HER2-enriched subtype is generally HER2-pos and 70-80% of the basal-like breast cancers are classified as TNBC (33). The C3-TAg model is most similar to human basal-like TNBC (34, 35).

Through the analysis of tumor organoid invasion dynamics, a data driven functional dissection of cellular re-programming and quantitative imaging of fixed tumor sections, we discovered a Tgfβ regulated program that was necessary to induce and sustain a trailblazer state in C3-TAg tumors and validated this discovery in breast cancer patient tumor samples. Surprisingly, the Tgfβ regulated trailblazer cells were not specified by Krt14 expression and were more aggressive than a previously defined Krt14 expressing trailblazer population detected in a luminal B GEMM, indicating that there are multiple different programs for conferring trailblazer ability in breast tumors. In addition, trailblazer cells induced by Tgfβ signaling were themselves heterogeneous, having different degrees of re-programming and capacities to initiate invasion. Tgfβ signaling collaborated with a parallel Egfr-Mek1/2-Erk1/2-Fra1 pathway to coordinate the expression of a subset of pro-motility genes, including Axl. Interestingly, the integration was also equivalently antagonistic in scope, with Egfr activity restricting the Tgfβ induced expression of a cohort of pro-invasive genes. Notably, clinically approved inhibitors of Egfr, Mek1/2 and Axl suppressed collective invasion. This indicated that the collateral inhibition of subset of pro-invasive genes by the Egfr pathway was a feature of the trailblazer state. Thus, our results reveal how a new type of trailblazer cell is induced by a therapeutically addressable compromise between Tgfβ and Egfr signaling programs.

## RESULTS

### The trailblazer cells in C3-TAg basal-like tumors are distinct from trailblazer cells in PyMT luminal B tumors

To define factors that initiate the conversion to a trailblazer state, we investigated the invasive properties of tumor cells isolated from the C3-TAg GEMM of basal-like TNBC (34, 35) and **(Supplementary file 1)**. We first determined if there was heterogeneity with respect to the ability to invade in the tumor populations, which would indicate the potential for trailblazer-opportunist relationships. Indeed, there was extensive heterogeneity in the invasive ability of tumor organoids and single dissociated tumor cells **(Figure 1A, Figure 1—figure supplement 1A-B, Figure 1—Videos 1-2)**. Clonal invasive (D6 and C7) and clonal non-invasive cells (B6 and G2) were also derived from a single C3-TAg tumor **(Figure 1—figure supplement C-E, Supplementary file 1)**. To test whether the invasive subpopulations in C3-TAg tumors could lead the invasion of siblings, we generated heterogeneous clusters containing C3-TAg tumor cells and intrinsically non-invasive B6 cells **(Figure 1B, Figure 1—Videos 3-5)**. The C3-TAg tumor cells and the D6 cells led the collective invasion of B6 cells **(Figure 1B, Figure 1—figure supplement 1E, Figure 1—Videos 3 and 6)**. These combined results indicated that there were trailblazer subpopulations in C3-TAg tumors with an enhanced ability to lead the collective invasion of intrinsically less invasive siblings.

**Figure 1 with 1 supplement.**
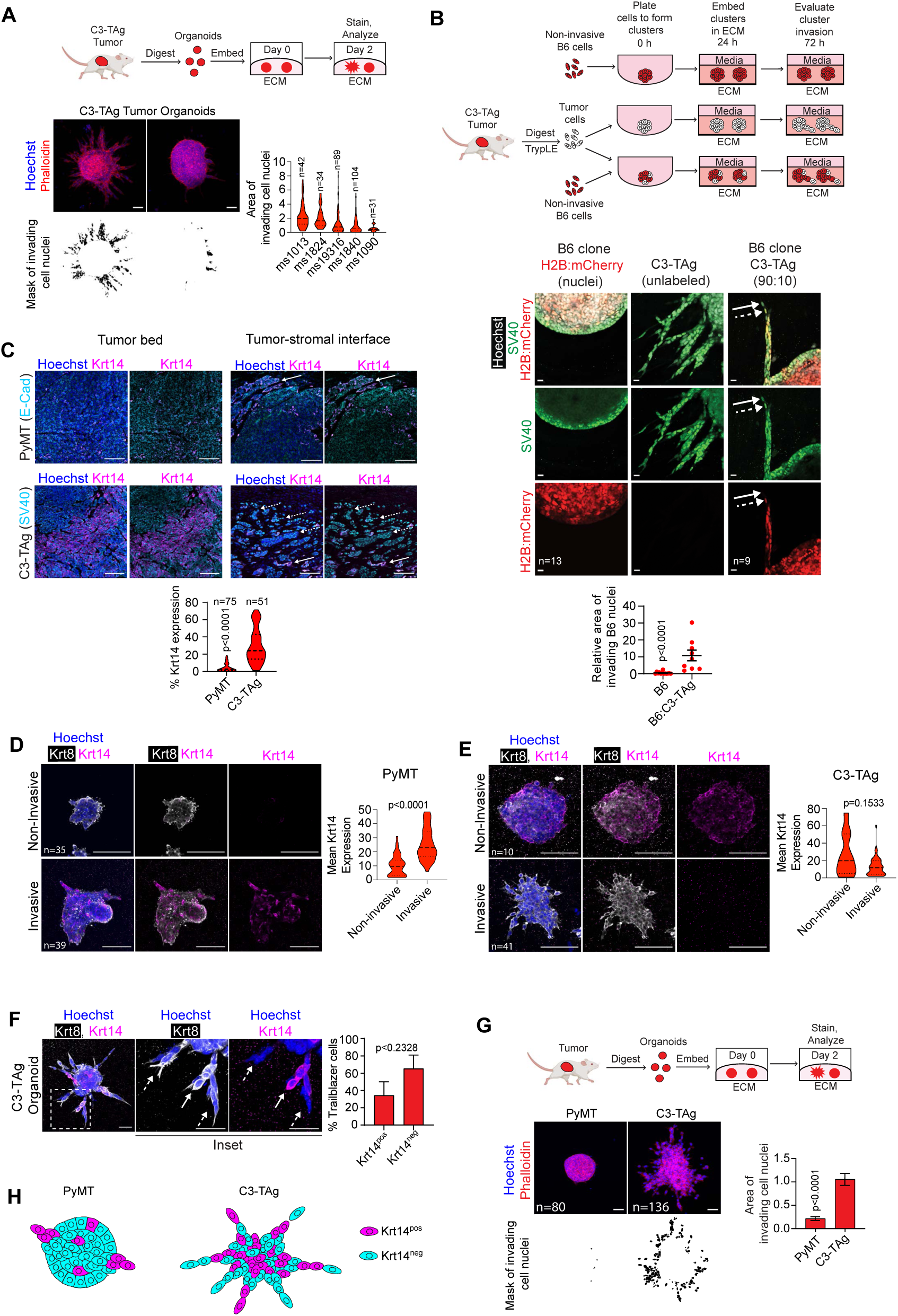
Trailblazer cells in C3-TAg basal-like tumors are distinct from trailblazer cells in PyMT luminal B tumors. **A.** There is intra- and inter-tumor invasive heterogeneity in C3-TAg organoids. Graphic shows the process for the derivation and analysis of tumor organoids. Representative images show the invasive heterogeneity of C3-TAg organoids derived from a single tumor. The area of invasion is determined using an image mask of the nuclei of invading cells. Violin plot shows the quantification of organoid invasion within 48 h of tumor isolation. (Mann Whitney test, n= organoids). Scale bars, 50 µm. **B.** C3-TAg tumors contain trailblazer cells that can lead the collective invasion of intrinsically less invasive cells. Graphic shows the process for generating homogeneous and heterogeneous clusters. Images show clusters immunostained with anti-SV40 antibody (detects TAg in all tumor cells), counterstained with Hoechst (nuclei) and H2B:mCherry fluorescence (B6 cells only). C3-TAg tumor cells (unlabeled) promoted the invasion of B6 cells (H2B:mCherry). Solid arrow indicates a C3-TAg tumor cell leading invasion. Dashed arrow indicates a B6 cell invading behind the C3-TAg cell. Graph shows the relative area of B6 cell invasion from 3 independent experiments (mean±SEM, Mann Whitney test, n=clusters). Scale bars, 20 µm. **C.** Krt14 expression does not correlate with C3-TAg tumor invasion. Krt14, SV40 (C3-TAg) and E-Cad (PyMT) expression in the tumor core and tumor-stromal interface of C3-TAg and PyMT tumors. Solid and dashed arrows indicate Krt14^pos^ and Krt14^neg^ cells respectively leading collective invasion. Violin plot shows the percentage of tumor cell area expressing Krt14. (Mann Whitney Test, n=ROIs from 5 tumors). Scale bars, 100 µm. **D.** Krt14 expression positively correlates with PyMT organoid invasion. Violin plot shows mean Krt14 expression in invasive and non-invasive PyMT organoids. (Mann Whitney Test, n=organoids from 3 tumors). Scale bars, 100 µm. **E.** Krt14 expression does not correlate with C3-TAg organoid invasion. Violin plot shows mean Krt14 expression in invasive and non-invasive C3-TAg organoids. (Mann Whitney Test, n=organoids from 3 tumors). Scale bars, 100 µm. **F.** Krt14 expression does not correlate with trailblazer phenotype in C3-TAg organoids. Solid and dashed arrows indicate Krt14^neg^ and Krt14^pos^ trailblazer cells respectively. Bar graph shows Krt14 expression status in trailblazer cells. (mean±SEM, unpaired Student’s t test, n=organoids from 3 tumors) Scale bars, 50 µm. **G.** C3-TAg organoids are more invasive than PyMT organoids. Bar graph shows the area of invasion (mean±SEM, Mann Whitney test, n=clusters, r=2) Scale bars, 50 µm. **H**. Model depicting the relationship between Krt14 expression and trailblazer phenotype in PyMT and C3-TAg tumors.

To understand how the trailblazer state was induced in C3-TAg tumor cells, we next evaluated the relationship between Krt14 expression and the ability of cells to lead collective invasion. Krt14 identifies cells that have converted from a luminal non-invasive state to a basal trailblazer state in the MMTV-PyMT (PyMT) GEMM model of luminal B breast cancer, which is commonly used to study breast cancer invasion and metastasis (20, 36). Based on these results, it has been suggested that a basal gene expression program exemplified by Krt14 expression is a universal feature of breast cancer trailblazer cells (20, 29). Consistent with prior results, Krt14 was expressed in PyMT trailblazer cells and Krt14 expression correlated with PyMT tumor organoid invasion **(Figure 1C-D)**. Krt14 was expressed in a greater proportion of cells in C3-TAg tumors compared to PyMT tumors **(Figure 1C)**. The high Krt14 expression was consistent with the early acquisition of basal traits detected in C3-TAg and TNBC tumors during the initial stages of growth within mammary ducts (37, 38). However, Krt14 expression did not correlate with C3-TAg tumor collective invasion or a trailblazer state in our analysis of primary tumors, tumor organoids and trailblazer and opportunist cell lines **(Figure 1C, E-F, Figure 1—figure supplement 1F)**. In fact, extensive collective invasion was detected in C3-TAg tumor organoids with few, if any, Krt14 expressing cells **(Figure 1E)**. Krt14^neg^ C3-TAg tumor cells also led collective invasion as frequently as Krt14^pos^ cells **(Figure 1F)**. C3-TAg organoids were relatively more invasive than PyMT organoids, demonstrating that the C3-TAg tumor invasion was more aggressive than the Krt14 *s*pecified invasion of PyMT tumors **(Figure 1G, 1H, Figure 1—figure supplement 1G, Figure 1—Videos 7-8)**. Together, these results suggested that essential features of the C3-TAg trailblazer program were different than those of the PyMT trailblazer program, and that and more than a single Krt14 specified trailblazer program functioned in breast cancer patient tumors **(Figure 1H)**.

### Trailblazer cells in C3-TAg GEMM tumors are distinguished by a subset of EMT traits

Given the lack of concordance with the established Krt14 specified trailblazer program, we next took an unbiased approach to identify the key trailblazer regulatory program features in C3-TAg tumors. To do this, we used next generation RNA sequencing to define differentially expressed genes (DEGs) in tumor organoids from 4 C3-TAg tumors compared with non-invasive spheroids formed by 3 clonal C3-TAg tumor derived cell lines **(Figure 2A, Figure 2—figure supplement 1A)**. Among epithelial keratins, Krt14 expression was reduced in C3-TAg tumor organoids compared to the non-invasive clonal C3-TAg cells, consistent with a lack of Krt14 involvement in C3-TAg collective invasion (**Figure 2B, Supplementary file 2)**. Expression of the transcription factor p63 was also reduced in C3-TAg tumor organoids and trailblazer clones, further indicating that canonical traits of normal basal mammary epithelial cells and Krt14 specified trailblazer cells (20) were not essential features of C3-TAg trailblazer cells (**Figure 2B, Figure 2—figure supplement 1B, Supplementary file 2)**. In addition, Krt14 expression was only detected in 1 out of 4 trailblazer populations in human breast cancer cell lines, which were all classified as TNBC **(Figure 2—figure supplement 1C)**. The expression of p63 was also low in all 4 trailblazer populations **(Figure 2—figure supplement 1C)**, consistent with our previous results showing that p63 expression is sufficient to suppress the induction of a trailblazer state (25).

**Figure 2 with 1 supplement.**
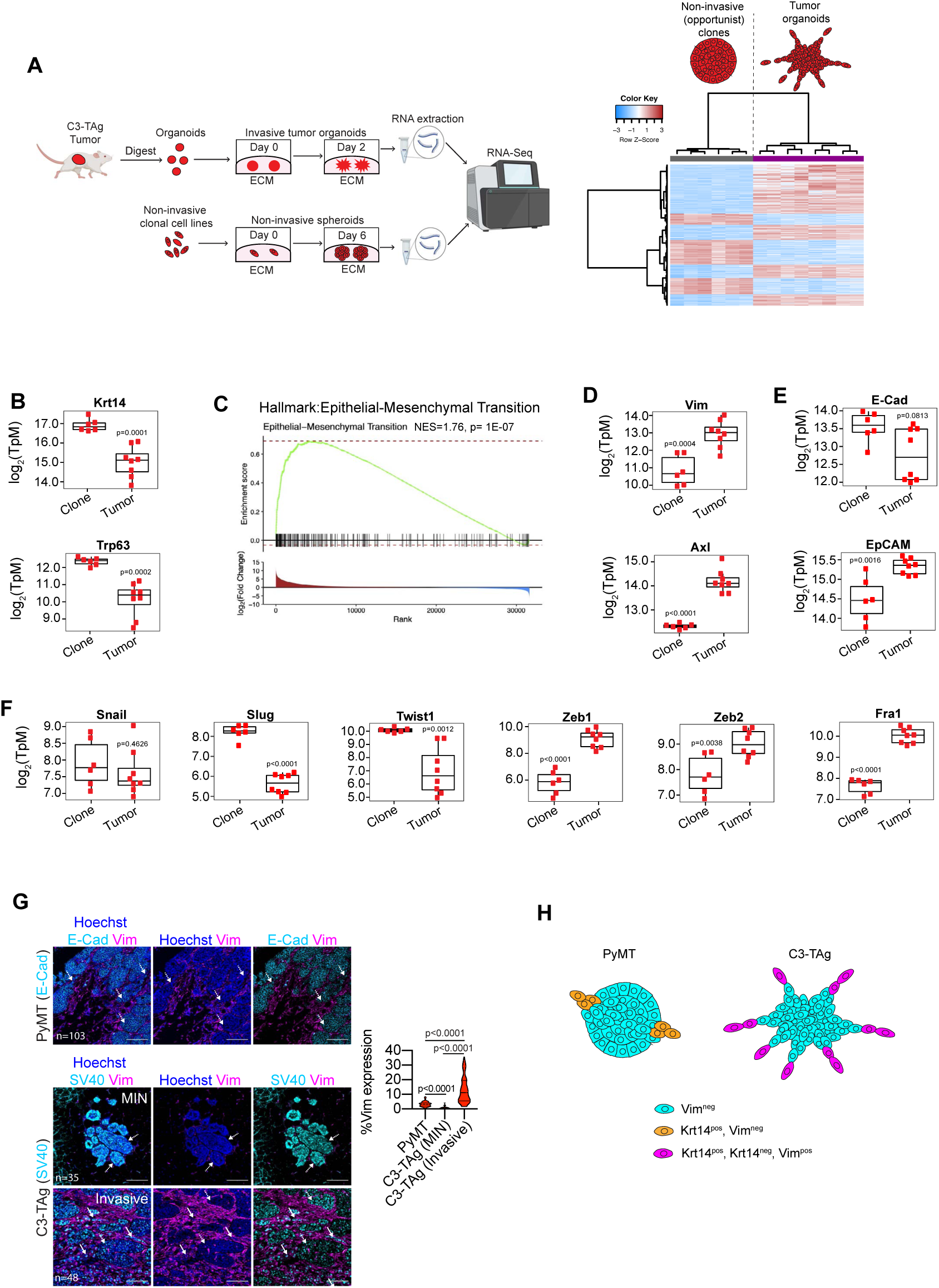
Trailblazer cells are distinguished by a subset of EMT traits in C3-TAg basal-like mammary tumors. **A.** Graphic shows preparation of C3-TAg organoids and non-invasive clonal cell line spheroids for RNA-seq analysis. Heatmap shows unsupervised clustering and differentially expressed genes in non-invasive C3-TAg clones (B6, C6, E8) and invasive organoids from 4 C3-TAg tumors. **B.** Boxplots showing the expression of the basal genes Krt14 and Trp63. **C.** There was increased expression of Hallmark EMT genes in C3-TAg tumor organoids compared to non-invasive C3-TAg clones. Plot shows the details of the Hallmark EMT generated by GSEA. **D-F.** Boxplots showing the expression of canonical mesenchymal (Vimentin, Axl), epithelial (E-cadherin, EpCAM) and EMT-TF (Snai1, Slug, Twist1, Zeb1, Zeb2, Fra1) genes in C3-TAg tumor organoids compared to non-invasive clonal spheroids from RNA-seq data (Mann Whitney test for E-cad, Student’s t-test for all others, n=6 for clones, (3 biological replicates); n=8 for tumor (4 biological replicates). **G.** Vimentin is more highly expressed in invasive C3-TAg primary tumors compared to noninvasive C3-TAg tumors and PyMT tumors. Solid arrows indicate Vim^pos^ tumor cells leading collective invasion. Dashed arrows indicate Vim^neg^ tumor cells. Violin plot shows percentage of tumor cell area expressing Vimentin (Mann Whitney Test, n=ROIs from 5 tumors). Scale bars, 100 µm. **H.** Model showing the distinct features of trailblazer cells in PyMT and C3-TAg tumors.

Genes highly expressed in C3-TAg tumor organoids were associated with biological processes that contribute to invasion (ECM Receptor Interaction, Focal Adhesions and Regulation of the Actin Cytoskeleton) and EMT **(Figure 2C-D, Figure 2—figure supplement 1D-E, Supplementary file 2)**. The C3-TAg tumor organoids retained canonical epithelial gene expression as well, indicating that the majority of cells were in a hybrid EMT state (**Figure 2E, Supplementary file 2)**. The EMT inducing transcription factors (EMT-TFs) Zeb1, Zeb2 and Fra1 were more highly expressed in C3-TAg organoids, consistent with their high expression in trailblazer cells in human breast cancer cell lines **(Figure 2F, Figure 2—figure supplement 1C)**. However, the EMT-TFs Snail, Slug and Twist1 were expressed at similar or higher levels in the noninvasive clones, which was consistent with a lack of concordance between the induction of a trailblazer state and Snail, Slug and Twist1 expression in human breast cancer cell lines **(Figure 2F, Figure 2—figure supplement 1C)**. These results indicated that the specific features of a trailblazer EMT program determined the invasive phenotype of C3-TAg tumor cells, rather than simply the presence or absence of EMT traits, consistent with results obtained analyzing human breast cancer cell lines (25, 31). Our combined results demonstrated that an EMT program exemplified by Vimentin expression conferred a trailblazer state in C3-TAg tumor subpopulations **(Figure 2G)**.

To understand the relationship between the EMT program features of the C3-TAg organoids and collective invasion, we evaluated the expression of the microfilament protein Vimentin, a common EMT marker that was more highly expressed in C3-TAg organoids **(Figure 2D)**. Vimentin expression was more frequently and extensively detected in invasive C3-TAg tumor cells compared to non-invasive C3-TAg mammary intraepithelial neoplasia (MIN) cells, consistent with Vimentin expression specifying a trailblazer EMT state in C3-TAg tumors. Notably, Vimentin expression was rarely detected in invasive PyMT tumor cells, further indicating that the regulatory programs inducing a trailblazer state in C3-TAg and PyMT tumors were distinct **(Figure 2G, Figure 2—figure supplement 1F)**. The heterogeneity of Vimentin expression in C3-TAg tumors was consistent with the heterogeneity in the extent of C3-TAg organoid invasion **(Figure 1A)**. Vimentin^pos^ and Vimentin^neg^ cells also collectively invaded together in C3-TAg primary tumors and organoids, suggesting that there was heterogeneity in collectively invading cell populations with respect to EMT state **(Figure 2G, Figure 2—figure supplement 1F)**.

### Trailblazer cells are specified by Vimentin expression in basal-like breast cancer patient tumors

We next investigated the characteristics of collectively invading cells in breast cancer patient tumors. KRT14 was expressed at a high level in basal like patient tumors compared to luminal B type breast cancer patient tumors, consistent with our results comparing the C3-TAg basal like and PyMT luminal B breast cancer GEMMs **(Figure 3A)**. However, KRT14 expression did not correspond with the induction of collective invasion in TNBC patients, identical to the lack of association between Krt14 expression and invasion in C3-TAg primary tumor and organoids **(Figure 3B, Figure 3—figure supplement 1A)**. In contrast, Vimentin expression was higher in the basal-like breast cancer compared to other breast cancer subtypes. Importantly, Vimentin expression was heterogeneous in TNBC tumor sections, with a trend towards more extensive Vimentin expression in regions of collective invasion **(Figure 3C-D, Figure 3—figure supplement 1B)**. These combined results indicated that the difference in trailblazer program features in C3-TAg (basal-like) and PyMT (luminal B) tumors reflected a feature of inter-tumor heterogeneity found in human breast cancer.

**Figure 3 with 1 supplement.**
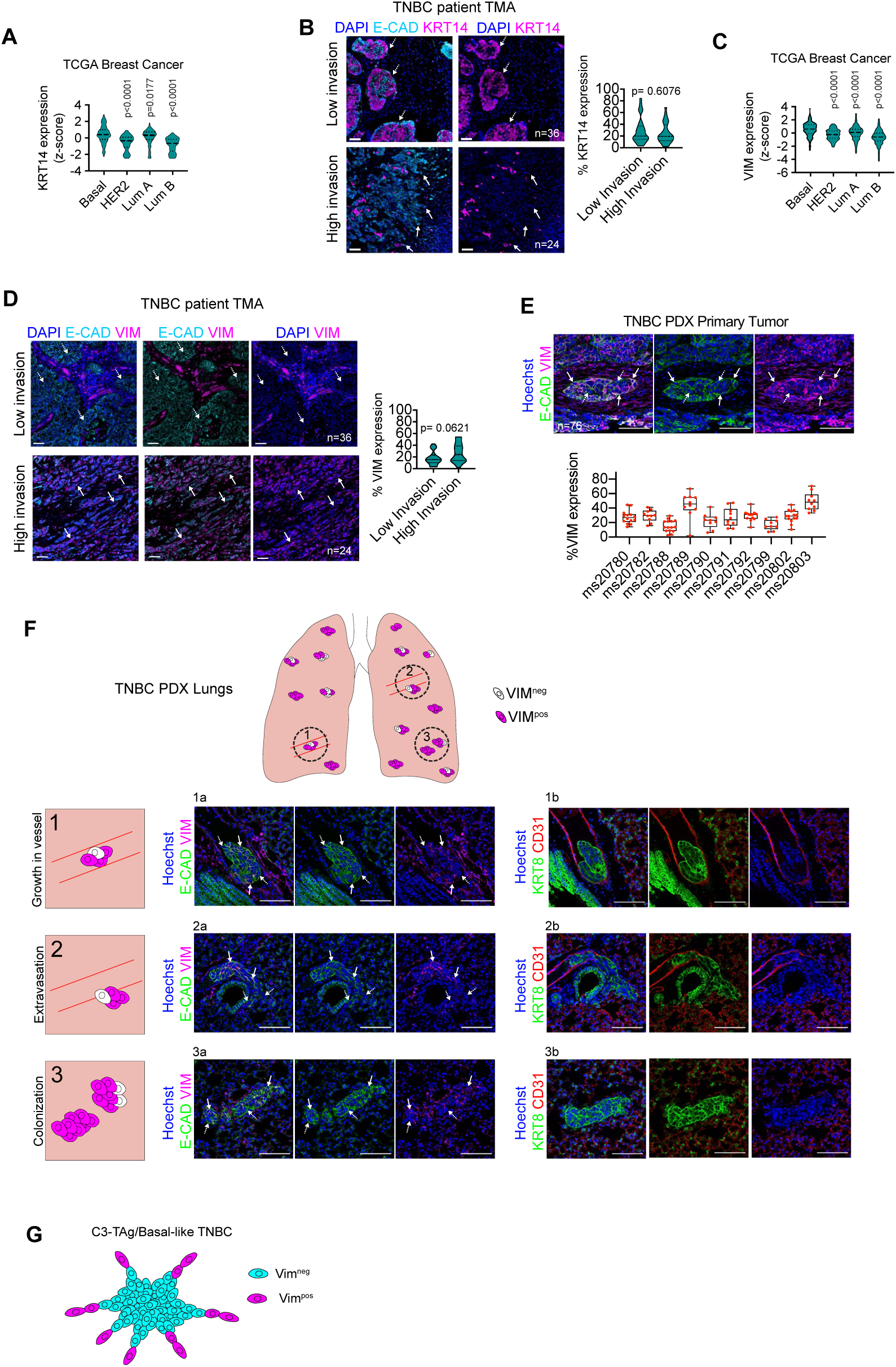
Trailblazer cells are distinguished by a subset of EMT traits in basal-like subtype of breast cancer. **A.** KRT14 mRNA is more highly expressed in the basal-like breast cancer patient tumors compared to other breast cancer subtypes. Z-scores relative to all samples are shown (Kruskal Wallis test with Dunn’s multiple comparisons test) From the TCGA. P-values are in comparison to the basal-like patient tumors. **B.** Krt14 expression in TNBC patient tumors does not correlate with invasion. Violin plot shows percentage of Krt14 expressing cells in regions of low (dashed arrows) and high invasion (solid arrows) (Mann Whitney test, n=tumor sections). Scale bars, 50 µm. **C.** VIM mRNA is more highly expressed in the basal-like breast cancer patient tumors compared to other breast cancer subtypes. Z-scores relative to all samples are shown (Kruskal Wallis test with Dunn’s multiple comparisons test) From the TCGA. P-values are in comparison to basal-like tumors. **D.** Invasive regions in TNBC patient tumors trend towards higher Vimentin expression (solid arrows) compared to regions of low invasion (dashed arrows). Violin plot shows percentage of Vimentin expressing tumor cells (Mann Whitney test, n=tumor sections). Scale bars, 50 µm. **E.** Vimentin is expressed in trailblazer cells in HCI-001 basal-like TNBC patient derived xenografts. Solid arrows indicate high Vimentin expression. Dashed arrows indicate low Vimentin expression. Violin plot shows percentage of tumor cell area expressing Vimentin (Mann Whitney Test, n=ROIs from 10 HCI-001 tumors). Scale bars, 100 µm. **F.** metastasis data. **G.** Model showing that Vimentin expression specifies trailblazer cells in basal-like C3-TAg tumors and TNBC patient tumors.

Metastasis is rarely detected in female mice bearing C3-TAg mammary tumors, likely because the rapid rate growth in primary tumors requires mice to be euthanized before metastases have time to develop (34). Therefore, to understand the relationship between the induction of Vimentin high cells and metastasis, we evaluated Vimentin expression in metastatic HCI-001 patient derived xenograft (PDX) TNBC tumors and their corresponding lung metastases (39). Intra- and inter-tumor heterogeneity in Vimentin expression was detected in the HCI-001 primary tumors **(Figure 3E)**. The heterogeneity and range in Vimentin expression was similar to what was observed in C3-TAg tumors and TNBC patient tumors **(Figure 2G, Figure 3D)**. An evaluation of the lungs revealed disseminated HCI-001 tumor cell clusters in the pulmonary vasculature **(Figure 3F, top row)**, partially localized within the vasculature and lung tissue **(Figure 3F, middle row)** and fully localized within lung tissue **(Figure 3F, bottom row)**. Notably, heterogeneous Vimentin expression was detected in each category of disseminated cell clusters, consistent with the induction of Vimentin-positive trailblazer cells metastasizing to the lungs **(Figure 3F)**. Together, our analysis indicated that Vimentin expression specified a trailblazer state in basal-like TNBC tumors, consistent with the features of the C3-TAg trailblazer populations.

### Tgfβ induces a trailblazer state in C3-TAg tumors

We next sought to define regulatory factors that activated the Vimentin specified trailblazer program. Gene set enrichment analysis (GSEA) indicated that Tgfβ signaling was more active in C3-TAg tumor organoids than non-invasive spheroids **(Figure 4A, Supplementary file 2)**. Thus, we evaluated the contribution of Tgfβ towards regulation of a trailblazer state. A pharmacological inhibitor of Tgfβ receptor 1 (Tgfbr1), A83-01, suppressed the invasion of freshly derived C3-TAg tumor organoids **(Figure 4B)**. 1863 cells derived in the presence of A83-01 were also less invasive than 1863T cells established from the same tumor in the absence of A83-01 **(Figure 4—figure supplement 1A, Supplementary file 1)**. Depletion of the Tgfβ regulated transcription factor Smad3 reduced the invasion of TGFβ1 stimulated 1863 cells and 1863T cells, indicating that TGFβ regulated gene expression contributed to the observed phenotypes **(Figure 4—figure supplement 1B-C)**. These results indicated that Tgfβ signaling was necessary to sustain a trailblazer state in C3-TAg cells.

**Figure 4 with 3 supplements.**
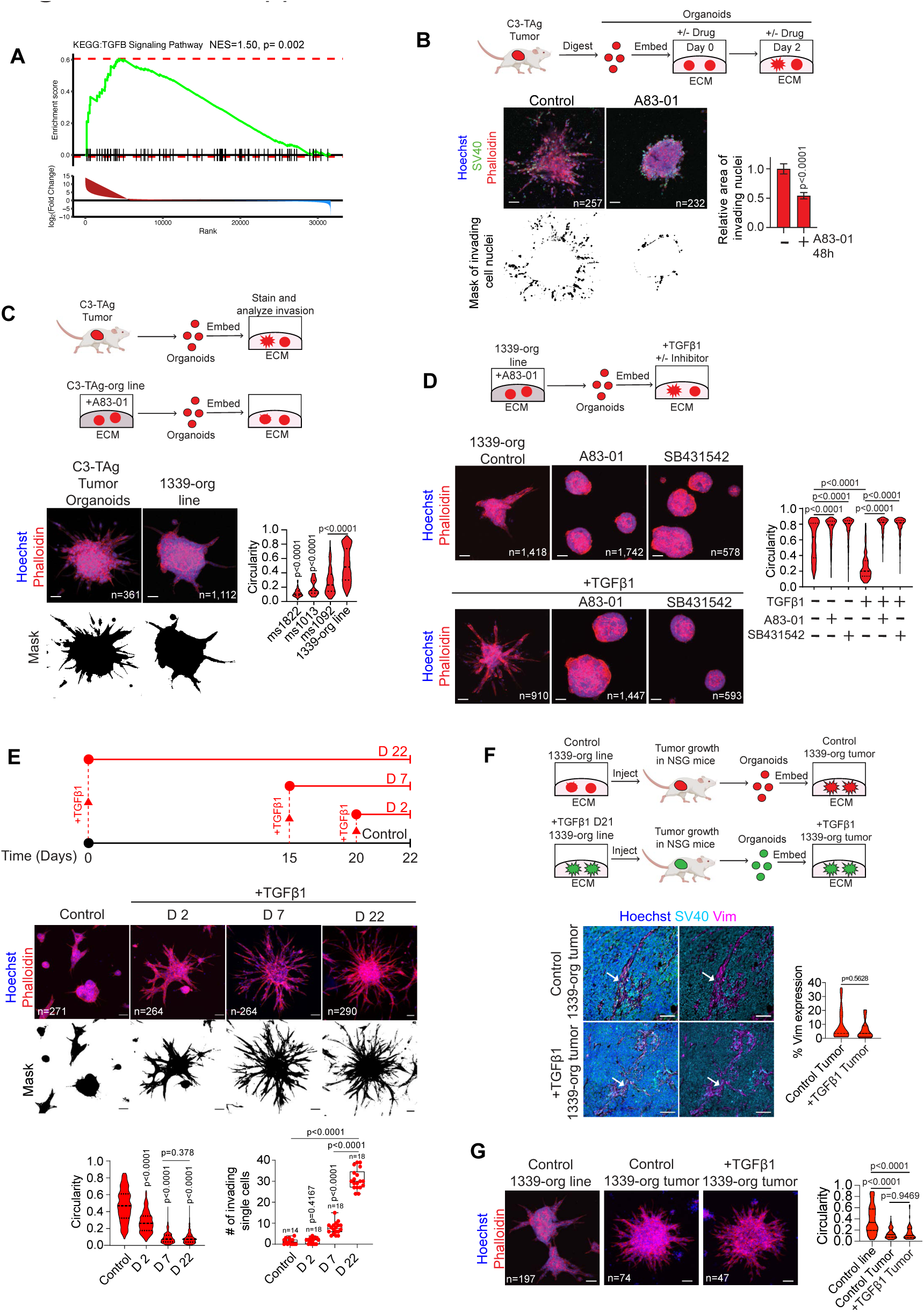
TGFβ induces a trailblazer state in C3-TAg tumors. **A.** GSEA was performed on RNA-seq data from C3-TAg organoids and non-invasive C3-TAg clonal cell line spheroids to identify pathway enrichment in C3-TAg organoids. Plot shows the GSEA for the KEGG: TGFβ signaling pathway. **B.** The Tgfbr1 inhibitor A83-01 suppresses C3-TAg organoid trailblazer phenotype within 48 h of tumor isolation. Graph shows the quantification of invasion (mean±SEM, Mann Whitney test, n=organoids from 4 tumors). Scale bars, 50µm. **C.** 1339-orgs grown in A83-01 were less invasive than organoids tested immediately after isolation from primary tumors. Graph shows organoid circularity. Greater circularity indicates a reduced trailblazer phenotype (mean±SEM, Mann Whitney test, n=organoids). Scale bars, 50 µm. **D**. Tgfbr1 inhibitors suppress the autocrine and exogenous TGFβ1 induced trailblazer phenotypes in 1339-orgs. Organoids were treated with the Tgfbr1 inhibitors A83-01 (500 nM) or SB431542 (1 µM) and exogenous TGFβ1 (2 ng/ml) as indicated. Violin plots show the quantification of organoid circularity (Mann Whitney test, n=organoids, r=3). Scale bars, 50 µm. **E.** Persistent exposure to exogenous TGFβ1 progressively enhanced the invasive phenotype in C3-TAg organoid lines. Graphs show the quantification of organoid circularity and the number of invading single cells (Mann Whitney test, n=organoids, r=3) Scale bars, 50 µm. **F.** Graphic shows the process for generating new primary tumors from control 1339-org line cells (Control 1339-org tumors) and 1339-org cells treated with TGFβ1 for 21 days (+TGFβ1 1339-org tumors). The expression of Vimentin was similar in Control 1339-org tumors and +TGFβ1 1339-org tumors. Violin plot shows quantification of Vimentin expression (Mann Whitney Test, 4 tumors from each group). Scale bars, 100 µm. **G.** The trailblazer phenotype of control 1339-org line cells was enhanced after tumor growth and similar to +TGFβ1 1339-org tumor cells. Violin plot shows quantification of organoid circularity (Mann Whitney Test, n=organoids representative of 4 tumors from each group).

To determine if Tgfβ could initiate the conversion from an opportunist to a trailblazer state, we first established organoid lines from 3 different C3-TAg tumors in the presence of A83-01 for at least 4 weeks **(Figure 4—figure supplement 2A, Supplementary file 1)**. These conditions yielded 1339-org, 1863-org and 1788-org tumor organoid lines that had a diminished Tgfβ activity and a reduced ability to invade relative to tumor organoids tested immediately after isolation from C3-TAg tumors **(Figure 4C, Figure 4—figure supplement 2A-C, Supplementary file 1)**. Removal of A83-01 or SB431542, a second inhibitor of Tgfbr1, promoted tumor organoid line invasion **(Figure 4D, Figure 4—figure supplement 2B-C)**. The invasion of the tumor organoid lines was further enhanced by exogenous TGFβ1 within 48 h **(Figure 4D, Figure 4—figure supplement 2B-C, Figure 4—Videos 1-3)**. Persistent exposure of 1339-org cells to Tgfβ signaling for up to 22 days progressively increased the rate of collective invasion **(Figure 4E, Figure 4—Videos 1-4)**. TGFβ signaling for at least 7 days also increased the frequency that trailblazer cells detached from collectively invading cohorts and invaded as single cells, suggesting that at least some single cell invasion reflected a loss of cohesion rather than the activation of a separate invasion program **(Figure 4E, Figure 4—Video 4)**.

To understand how the induction of a trailblazer state by Tgfβ influenced the properties of cells during primary tumor growth, we injected control 1339-org cells cultured in A83-01 (Control) or 1339-org cells stimulated with Tgfβ for 21 days (+Tgfβ) into the mammary fat pads of immune-compromised mice. Both Control and +Tgfβ 1339-org cells initiated the growth of primary tumors **(Figure 4F)**. Interestingly, newly isolated organoids from the Control and +Tgfβ tumors were equivalently invasive **(Figure 4G)**. This similarity was a consequence of the induction of a trailblazer state in the Control 1339-org primary tumors, which was indicated by the new organoids derived from Control 1339-org tumors being more invasive than Control 1339-org line cells prior to tumor growth **(Figure 4G)**. This induction of a trailblazer state was potentially due to stimulation of primary tumor cells by Tgfβ in the tumor microenvironment produced my nontumor cells, such as macrophages (40). Consistent with the equivalent invasion of the new primary tumor organoids, the tumors that developed from Control and +Tgfβ organoids contained a similar fraction of Vimentin expressing cells **(Figure 4F)**. The equivalent invasive behavior of the tumors also corresponded with a similar number of disseminating tumor cells detected in the blood **(Figure 4—figure supplement 3A)**. Metastatic tumor cells were not detected in the lung, indicating that the extent of dissemination was not yet sufficient to initiate colonization, consistent with metastasis being a highly inefficient process **(Figure 4—figure supplement 3B)**. Thus, Tgfβ signaling promoted the conversion of C3-TAg cells to a trailblazer state.

### Tgfβ induces multiple trailblazer states through distinct phases of re-programming

To understand how Tgfβ induced a trailblazer state, we used RNA-sequencing (RNA-seq) to analyze gene expression changes 1339-org cells after 2, 7 and 22 days after Tgfβ pathway activation. Hierarchical clustering revealed 4 different groups of genes regulated by TGFβ **(Figure 5A, Supplementary file 3)**. If cells were undergoing binary conversion between states, the population averaged mRNA expression obtained by RNA-seq would have shown 2 groups of genes (suppressed and induced) undergoing a gradual change in expression as percentage of cells in the 2 different states shifted (41). The 4 different gene groups, each with a unique timing in their onset of change and maximal expression difference, therefore indicated that the Tgfβ stimulated C3-TAg tumor cells progressed through different stages of re-programming. Consistent with inducing multiple different states, TGFβ promoted a shift in the distribution of EpCAM expression rather than a switch between two unique EpCAM-high and EpCAM-low states, **(Figure 5—figure supplement 1A)**. TGFβ also induced a shift in the distribution of Vimentin expression intensity in organoids at day 7 and day 22, as opposed to a switch between two distinct Vimentin^neg^ and Vimentin^pos^ states **(Figure 5—figure supplement 1B)**.

**Figure 5 with 1 supplement.**
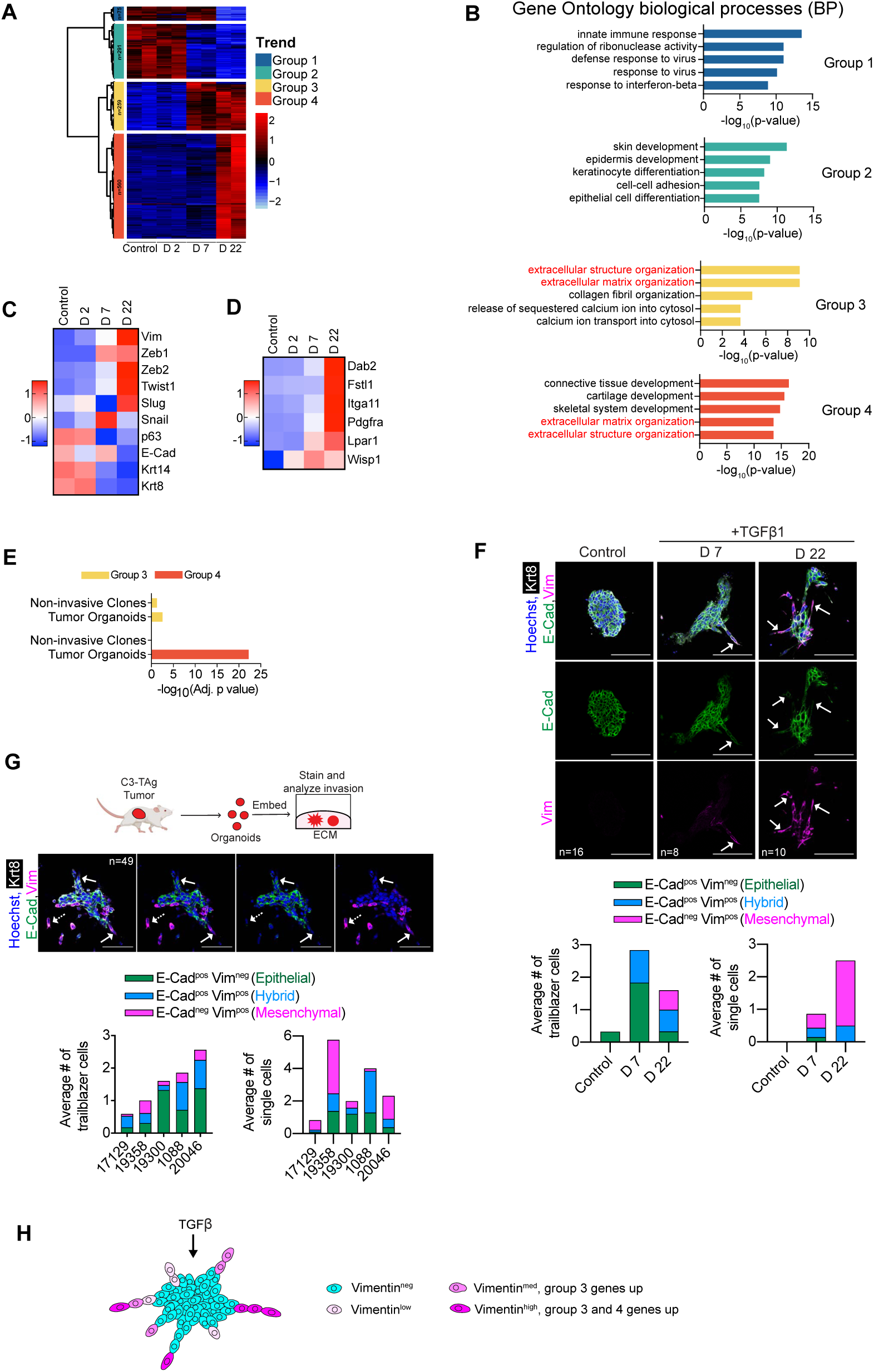
TGFβ1 induces multiple trailblazer states through distinct phases of re-programming. **A.**. RNA-seq was performed on 1339-org cells treated for 2, 7 and 22 days with TGFβ1. Heatmap shows unsupervised clustering of control and TGFβ1 treated cells along with 4 groups of DEGs. **B.** Graphs show the top 5 Gene Ontology terms associated with each group. **C.** Heatmap shows changes in expression of mesenchymal (Vim), EMT-TF (Zeb1 and 2, Twist1, Snail, Slug), epithelial (E-Cad), basal (p63, Krt14) and luminal (Krt8) markers after TGFβ1 exposure for 0, 2, 7 and 22 days. **D.** Heatmap shows that persistent activation of Tgfβ signaling increaseds the expression of previously defined trailblazer genes. **E.** Graph shows the enrichment of Group 3 and Group 4 induced genes in non-invasive clones and invasive C3-TAg tumor organoids. **F-G.** Trailblazer cells in multiple different EMT states lead collective invasion in freshly isolated C3-TAg tumor organoids and TGFβ1 treated 1339-org line organoids. Sections from paraffin-embedded organoids were immunostained with Vimentin and E-cadherin antibodies to define different EMT states. The phenotype of trailblazer cells leading invasion or invading single cells was quantified. Bar graphs indicate the mean number of trailblazer or single invading cells per organoid and their EMT phenotype. **H.** Model showing the heterogeneity in trailblazer states induced by TGFβ1.

The gene groups identified by RNA-seq were associated with distinct combinations of biological processes **(Figure 5B, Supplementary file 3)**. Of note, genes in Groups 3 and 4 were increased in expression with different kinetics in response to Tgfβ and involved with ECM regulation **(Figure 5B, Supplementary file 3)**. This indicated that there were different mechanisms for regulating genes associated with the same biological functions in response to TGFβ. Group 3 and 4 genes were also connected with unique functions, which indicated that the nature of reprogramming changed over time and was controlled by distinct mechanisms. Interestingly, genes commonly associated with EMT were included in Group 3 (Vcam1, Itgb3, Fn1) and Group 4 (Vimentin, N-cadherin, Periostin) **(Supplementary file 3)**. EMT-TFs were also split between Group 3 (Zeb1), Group 4 (Zeb2, Twist), or not included in any group (Snail) **(Figure 5C)**. E-cadherin (Group 1) and the mammary epithelial lineage markers p63, Krt8 and Krt14 (Group 2) were also decreased in expression on different time scales **(Figure 5C)**. Consistent with these results, high Vimentin and Zeb1 expression and reduced E-cadherin expression was observed in D6 and 1863T trailblazer cells, indicating that these cell lines reflected features of trailblazer cells after persistent activation of Tgfβ signaling **(Figure 5—figure supplement 1C-E)**. Previously defined trailblazer genes were in Group 3 (Wisp1) and Group 4 (Dab2, Fstl1, Itga11, Pdgfra, Lpar1) **(Figure 5D)**. Notably, Group 3 and Group 4 genes were enriched in the genes that were highly expressed in C3-TAg organoids, which indicated that Tgfβ re-activated a program that was present in C3-TAg tumors **(Figure 2A, 5E, Supplementary file 3)**.

The distinct expression profile of EMT-TFs, EMT effectors and ECM regulatory genes indicated that persistent Tgfβ pathway activation induced re-programming yielded multiple different trailblazer states in response to regulatory mechanisms that evolve over time. To test this possibility we immunostained 1339-org and C3-TAg tumor organoids with Vimentin and E-Cadherin antibodies, to specify different states within the spectrum of EMT re-programming. Vimentin^neg^/E-cadherin^pos^, Vimentin^pos^/E-cadherin^pos^ and Vimentin^pos^/E-cadherin^neg^ cells were all capable of leading collective invasion (**Figure 5F-G)**. Notably, the presence of more Vimentin^pos^/E-cadherin^pos^ and Vimentin^pos^/E-cadherin^neg^ trailblazer cells with increasing time after Tgfβ pathway activation correlated with an increased extent of 1339-org collective invasion (**Figure 5F)**. Our combined results suggested that there were multiple trailblazer states induced by Tgfβ and that they had different capacities to initiate collective invasion **(Figure 5H)**.

### The transcription factors Zeb1, Zeb2 and Fra1 confer a trailblazer state

To further understand how Tgfβ induced a trailblazer state, we next determined how siRNAs targeting 12 EMT regulatory transcription factors influenced the invasion of 1863T trailblazer cells. Smad3 siRNA transfection served as a positive control. The 1863T trailblazer cells test the requirements for trailblazer cell invasion after long-term activation of the Tgfβ pathway **(Figure 6—figure supplement 1A)**. The siRNAs targeting Fra1, Zeb1 and Zeb2 suppressed 1863T invasion with a p< 0.05 **(Figure 6A)**, consistent with Fra1, Zeb1 and Zeb2 being the EMT-TFs that were more highly expressed in C3-TAg tumor organoids (**Figure 2F)**. Fra1, Zeb1 and Zeb2 siRNAs also suppressed invasion to a similar extent as Smad3 siRNAs (**Figure 6A**). Thus we prioritized Fra1, Zeb1 and Zeb2 for further investigation.

**Figure 6 with 4 supplements.**
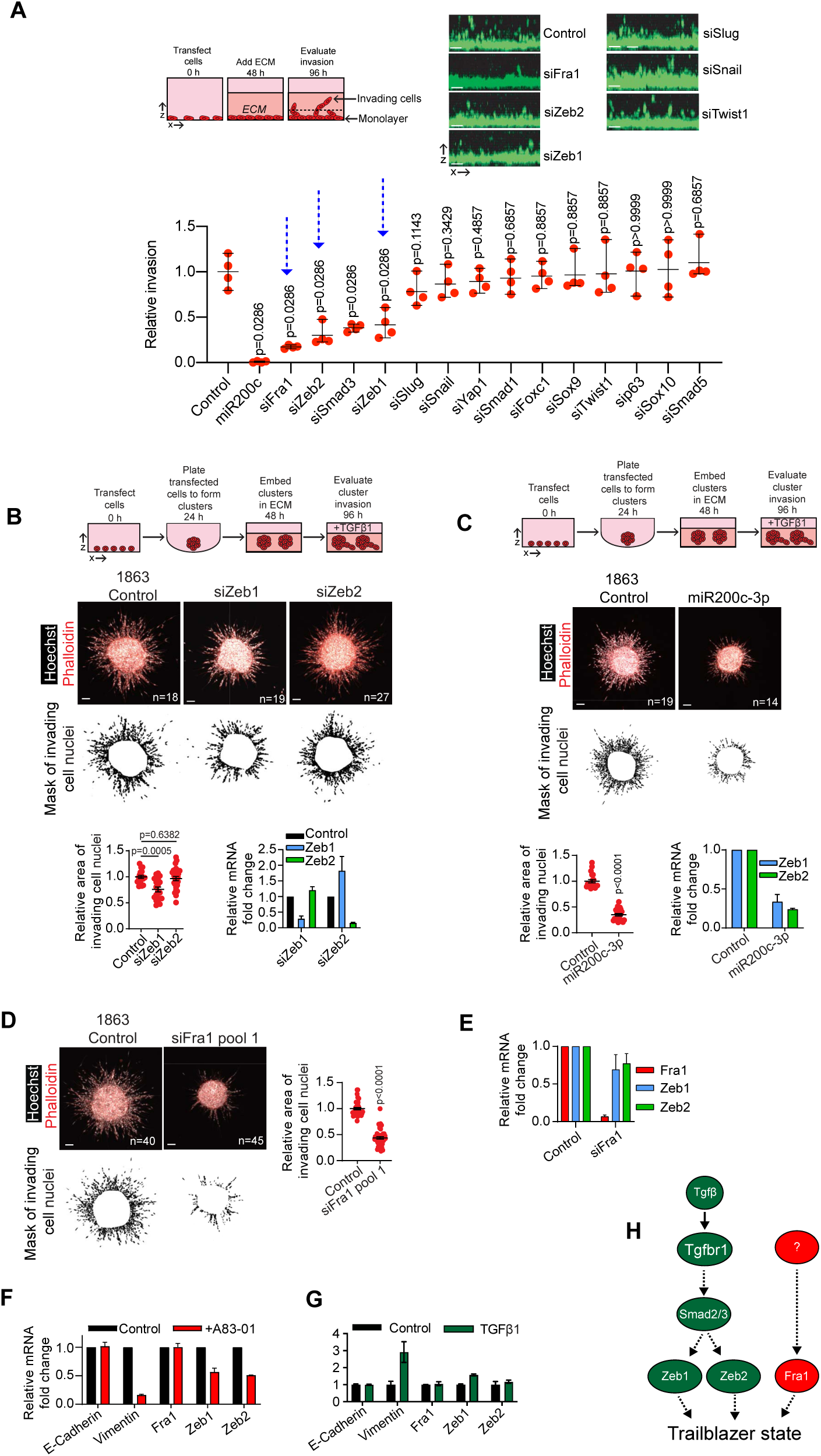
The transcription factors Fra1, Zeb1 and Zeb2 confer a trailblazer state. **A.** Invasion of 1863T cells transfected with siRNAs targeting TFs associated with EMT and invasion. Graph shows the quantification of cells vertically invading ≥40 µm into the ECM (mean±SEM, Mann Whitney test, n=4 wells from 2 independent experiments). Scale bars, 50 µm. Smad3 siRNA serves as a positive control for suppression of invasion. Blue arrows indicate genes prioritized for further analysis. **B.** Zeb1 siRNA transfection suppressed the invasion of TGFβ1 treated 1863 clusters. Dot plot shows the quantification of invasion (mean±SEM, Mann Whitney test, n=clusters, r=2). Scale bars, 50 µm. Bar graph shows Zeb1 and Zeb 2 expression (mean±SEM, n=2). **C.** Transfection with a miR200c-3p mimic decreased TGFβ1 treated 1863 cluster invasion and Zeb1 and Zeb2 expression. Dot plot shows the quantification of invasion (mean±SEM, Mann Whitney test, n=clusters, r=2). Scale bars, 50 µm. Bar graph shows Zeb1 and Zeb 2 expression, (mean±SEM, n=2). **D.** Fra1 siRNA pool 1 transfection suppressed GFβ1 treated 1863 cluster invasion (mean±SEM, Mann Whitney test n=clusters, r=5) Scale bars, 50 µm. **E.** Fra1 depletion did not suppress Zeb1 and Zeb2 expression (mean±SEM, n=2). **F-G**. E-cadherin, Vimentin, Fra1, Zeb1 and Zeb2 mRNA expression after 48 h treatment with **(F)** Tgfbr1 inhibitor A8301 and **(G)** TGFβ1 (mean±SEM, n=2). I. Model showing the parallel regulation of the trailblazer state by the Tgfβ signaling and a second pathway that induces Fra1.

The Zeb1 and Zeb2 siRNAs suppressed 1863T cluster invasion, demonstrating that Zeb1 and Zeb2 depletion influence 1863T invasion in multiple contexts **(Figure 6—figure supplement 1B)**. Our analysis in 1863T cells determined how Zeb1 and Zeb2 depletion influenced the invasion of trailblazer cells after long-term Tgfβ activation. To determine how Zeb1 and Zeb2 depletion influenced the induction of invasion upon the initial Tgfβ pathway stimulation, we analyzed the invasion of 1863 cells that were established and propagated in the presence of the Tgfbr1 inhibitor A83-01 before experimentation **(Figure 6—figure supplement 1A)**. Interestingly, only Zeb1 depletion suppressed TGFβ stimulated 1863 cluster invasion, whereas Zeb2 depletion had a negligible effect (**Figure 6B, Figure 6—figure supplement 1A)**. A miR200a-3p mimic, which suppressed Zeb1 and 2 expression, inhibited 1863 and 1863T invasion, further indicating Zeb1 and 2 contribute to the induction of a trailblazer state **(Figure 6C, Figure 6—figure supplement 1C)**. The requirement of Zeb1, but not Zeb2, for the invasion of 1863 cells upon initial TGFβ stimulation was consistent with our results showing that Zeb1 expression rapidly increased within 7 days while increased Zeb2 expression occurred after day 7 of extrinsic TGFβ exposure **(Figure 5C)**. Thus, our results suggested that the requirement of Zeb1 and Zeb2 may be dependent on the specific trailblazer state induced by Tgfβ signaling.

Additional investigation of Fra1 revealed that Fra1 depletion by 4 individual siRNAs and 2 different siRNA pools reduced 1863T and TGFβ stimulated 1863 invasion to a similar degree **(Figure 6D, Figure 6—figure supplement 2A-D)**. Fra1 depletion also reduced the invasion of D6 trailblazer cells and Tgfβ stimulated 1339 cells, which were derived from a different C3-TAg tumor than the 1863 and 1863T cells **(Figure 6—figure supplement 2E-G)**. Fra1 interacts with Jun to regulate gene expression as an AP-1 complex (42). Indeed, Jun was required for 1863T and Tgfβ stimulated 1863 cell invasion, suggesting Fra1 containing AP-1 complexes regulate the expression of genes necessary for the induction of a trailblazer state **(Figure 6—figure supplement 3A-C)**.

Exogenous Fra1 expression promotes Zeb1 and Zeb2 expression in a mouse mammary epithelial cell line, suggesting a potential connection between the functions of Fra1, Zeb1 and Zeb2 (43). However, Fra1 depletion in 1863T cells did not alter Zeb1 or Zeb2 expression, indicating that Zeb1 and Zeb2 are not regulated by endogenous Fra1 in trailblazer cells **(Figure 6E)**. In addition, Zeb1, Zeb2 and Vimentin expression were regulated by autocrine TGFβ signaling or exogenous TGFβ treatment whereas Fra1 expression was not influenced by autocrine or extrinsic TGFβ pathway stimulation (**Figure 5C, Figure 6F-G, Supplementary file 3)**. These results indicated that the regulation of Fra1 expression was uncoupled from Tgfβ, Zeb1 and Zeb2. Nevertheless, Fra1 was expressed at a higher level in C3-TAg tumor organoids compared to non-invasive spheroids, consistent with increased Fra1 expression contributing to the induction of a trailblazer state **(Figure 2F)**. In addition, FRA1 expression was higher in basal like breast cancer compared to other breast cancer subtypes **(Figure 6—figure supplement 4A)**. FRA1 expression also correlated with VIM (Vimentin) expression in breast cancer patients, suggesting that Fra1 potentially contributes to induction of a Vimentin specified trailblazer state during tumor progression in basal-like breast cancer patients **(Figure 6—figure supplement 4B)**. Thus, our data indicated that the induction of Fra1 and a trailblazer state in C3-TAg tumors required the activity of at least one additional signaling pathway acting in parallel with Tgfβ signaling **(Figure 6H)**. We therefore selected Fra1 for further analysis to more fully understand the global regulatory framework that empowered a trailblazer state.

### Egfr pathway activity promotes Fra1 expression and a trailblazer state

We next determined how Fra1 expression was regulated in C3-TAg tumor cells. Fra1 mRNA was more highly expressed in C3-TAg tumor organoids compared to opportunist spheroids in our RNA-seq analysis, indicating that the signaling pathway responsible for the control of Fra1 was more active in the C3-TAg organoids **(Figure 2F)**. GSEA identified multiple active KRAS regulated expression signatures that were more active in the C3-TAg tumor organoids **(Figure 7— figure supplement 1A-B, Supplementary file 2)**. The Kras gene can be amplified, but is not frequently mutated in C3-TAg tumors, indicating that pathway activation occurred upstream of Kras (44). Analysis of the genes in the active KRAS signature showed that the Egfr ligand Hbegf was more highly expressed in C3-TAg organoids than opportunist clones **(Figure 7A, Supplementary file 2)**. Like FRA1, HBEGF was more highly expressed in Basal type breast tumors **(Figure 7—figure supplement 1C)**. HBEGF expression correlated with FRA1 expression in breast cancer patient tumors as well, consistent with the possibility that Hbegf promotes Fra1 expression during tumor development **(Figure 7—figure supplement 1D)**. Further inspection found that the Egfr ligands Areg and Epgn were also expressed at a higher level in C3-TAg organoids **(Figure 7A, Supplementary file 2)**. These combined results prompted us to investigate how the Egfr-Ras-Mek1/2-Erk1/2 signaling pathway contributed to Fra1 expression and the induction of a trailblazer state.

**Figure 7 with 4 supplements.**
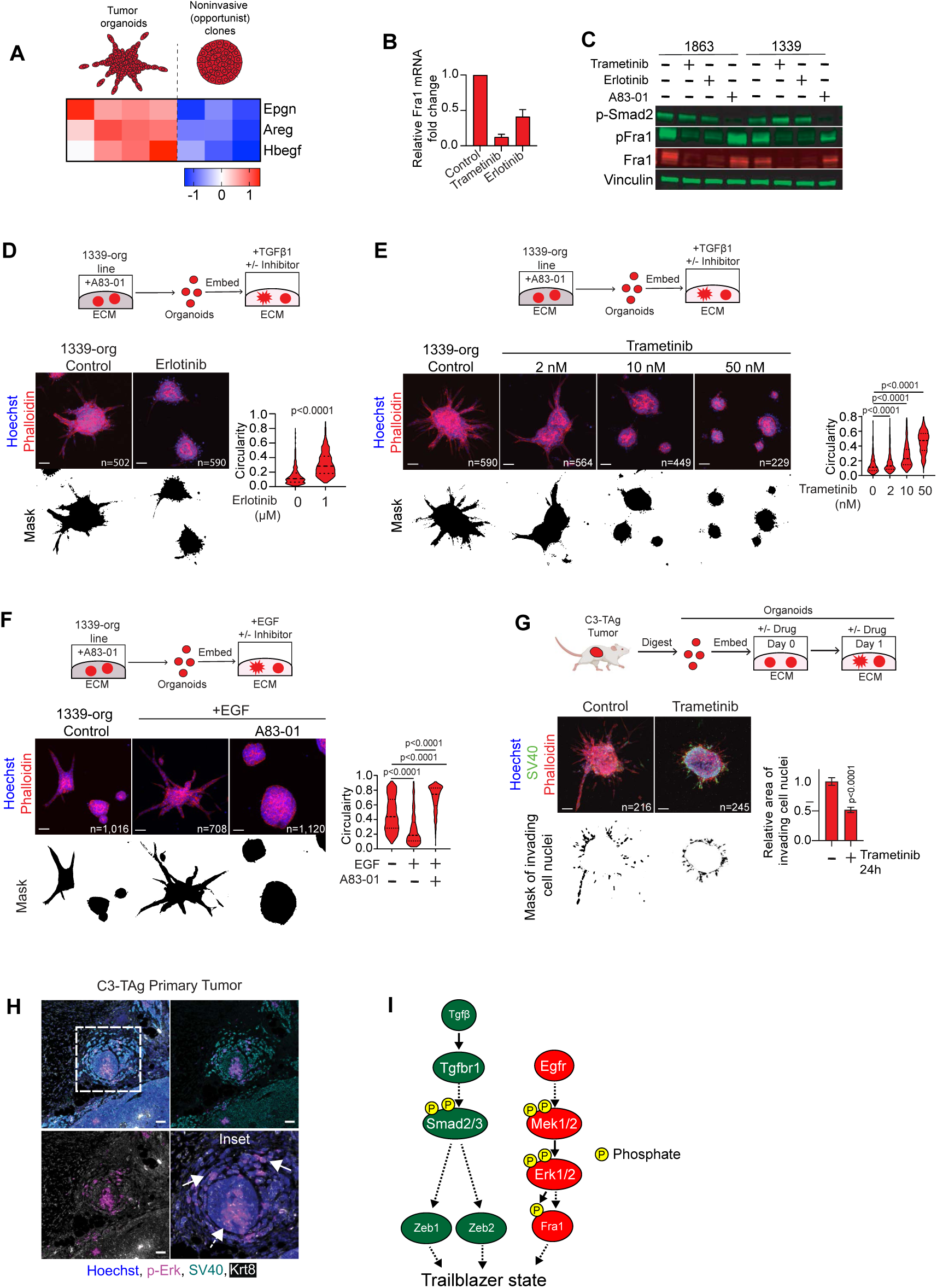
Egfr signaling regulates Fra1 expression and C3-TAg trailblazer state. **A.** Heatmap shows the expression of the Egfr ligands Epigen (Epgn), Amphiregulin (Areg) and Heparin-binding epidermal growth factor (Hbegf) in C3-TAg organoids and non-invasive clones. **B.** The Mek1/2 inhibitor trametinib (10 nM) and Egfr inhibitor erlotinib (1 µM) reduce Fra1 expression in the 1339-org line (mean±SEM, n=2). **C.** Immunoblot shows the response of 1863 and 1339 cells to trametinib, erlotinib and A83-01 treatment (n=2). **D.** Erlotinib (1 µM) suppresses the TGFβ1 induced invasion the 1339-org line. Violin plot shows the quantification of organoid circularity (Mann Whitney test, n=organoids, r=3) Scale bars, 50 µm. **E.** Trametinib suppresses the TGFβ1 induced invasion of the 1339-org line in a dose dependent manner. Violin plot shows the quantification of organoid circularity (Mann Whitney test, n=organoids, r=3). Scale bars, 50 µm. **F.** A83-01 suppresses the EGF induced invasion in the 1339-org line. Violin plot shows the quantification of organoid circularity (Mann Whitney test, n=organoids, r=4) Scale bars, 50 µm. **G.** Trametinib suppresses the invasion of freshly derived C3-TAg organoids within 24 h of original tumor processing. Graph shows the quantification of invasion from 4 different tumors (mean±SEM, Mann Whitney test, n=organoids) Scale bars, 50 µm. **H.** Images show dually phosphorylated Erk1/2 immunostaining in regions of high invasion (solid arrows) and low invasion (dotted arrow) in primary C3-TAg tumors. Scale bars, 50 µm. **I.** Model showing the parallel regulation of trailblazer cells by signaling pathways coordinated by Tgfβ and Egfr.

Erlotinib (Egfr inhibitor) and trametinib (Mek1/2 inhibitor) suppressed Fra1 mRNA and protein expression in C3-TAg cell lines and organoid lines **(Figure 7B-C, Figure 7—figure supplement 2A)**. By comparison, Jun expression was unchanged by treatment with either inhibitor **(Figure 7—figure supplement 2B)**. In addition, erlotinib and trametinib did not inhibit the activating phosphorylation of Smad2 or Zeb1 expression, indicating that the Egfr1-Mek1/2-Erk1/2-Fra1 did not influence cell state through autocrine Tgfβ signaling **(Figure 7C, Figure 7—figure supplement 2C)**. A83-01 suppressed Smad2 phosphorylation, but did not influence Fra1 expression and phosphorylation, further indicating that Tgfb signaling did not regulate Fra1 **(Figure 7C)**.

Erlotinib treatment reduced the TGFβ stimulated invasion of 1339-org cells **(Figure 7D)**. Trametinib treatment also blocked the initial stimulation of 1339-org invasion by Tgfβ, the invasion of 1339-org trailblazer cells following persistent Tgfβ treatment and the invasion D6 trailblazer cells (**Figure 7E, Figure 7—figure supplement 2D-E)**. Moreover, extrinsic EGF induced the invasion of 1339-org, 1863-org and 1788-org cells and this enhancement was blocked by A83-01, further indicating that the integration of Tgfbr1 and Egfr pathways promoted a trailblazer state **(Figure 7F, Figure 7—figure supplement 2F-G)**. Trametinib inhibited the invasion of C3-Tag primary tumor organoids and phosphorylated Erk1/2 was detected in collectively invading cells in C3-TAg primary tumors **(Figure 7G-H)**. Erk1/2 activity is pulsatile in response to EGF stimulation, and these pulses promote Fra1 expression (45). Thus, persistent Erk1/2 phosphorylation was not necessarily expected in all collectively invading cells or to promote Fra1 expression. Consistent with these results, A83-01, erlotinib and trametinib suppressed the invasion of human MCF10A-Trailblazer cells and MCFDCIS cells, which harbor features of basal-like breast cancer (25, 46). This indicated that the integration of Tgfb and Egfr pathway signaling was necessary for the induction of a trailblazer state in multiple contexts (**Figure 7—figure supplement 3A-C, Figure 7—figure supplement 4A-B)**. Together, these results indicate that the regulation of Fra1 by Egfr and Erk1/2 promotes the induction of a trailblazer EMT **(Figure 7I)**.

### A compromise between Tgfbr1 and Egfr regulated signaling programs confers the trailblazer state

To understand the contribution of the Tgfbr1 and Egfr regulated signaling pathways towards the induction of the trailblazer state, we performed RNA-seq on organoids treated with diluent, TGFβ, TGFβ+erlotinib, EGF and EGF+A83-01 **(Figure 8A, Figure 8—figure supplement 1A)**. Polar plots were used to cluster genes based on the nature and the degree of integration between the Tgfbr1 and Egfr signaling pathways (47). This analysis revealed multiple modes of interaction between the two pathways in response to either extrinsic TGFβ or EGF and that the type of interaction (positive or negative) was associated with different biological processes **(Figure 8B-C, Figure 8—figure supplement 1B-F, Supplementary files 4 and 5)**. Thus, the relationship was far more complex than the simple positive signal integration previously suggested by phenotypic analysis and the expression of specific canonical EMT program genes (48, 49). In addition, the gene expression changes induced by extrinsic TGFβ and EGF were associated with unique combinations of biological processes, indicating that extrinsic TGFβ and extrinsic EGF induced distinct types of trailblazer cells **(Figure 8C, Figure 8—figure supplement 1F)**. Nevertheless, there were conserved features of pathway integration that influenced the regulation of processes associated with invasion regardless of the external stimulus. Genes involved in controlling the organization and composition of the ECM, including previously defined trailblazer genes, were predominantly induced by Tgfbr1 and not positively impacted by Egfr signaling **(Figure 8C, Figure 8—figure supplement 1F, Supplementary files 4 and 5).** Notably, autocrine and extrinsic Egfr activation suppressed a number of Tgfbr1 regulated ECM reorganization genes and Hallmark EMT genes, including Postn, Bmp2, Itgb3, Orai2, Mmp11 and Erg **(Figure 8D, Figure 8—figure supplement 1G, Supplementary files 4 and 5)**, which can contribute to invasion and metastasis (50-55). By comparison, autocrine or extrinsic EGF activity alone, or in combination with Tgfbr1 signaling, induced the expression of genes that promote cell motility, such as Axl, Sema7a, Ncam1 and Itga2 **(Figure 8C and E, Figure 8—figure supplement 1F and H, Supplementary files 4 and 5)** and (56-59). Thus, the induction of a set of pro-motility genes corresponded with a compromise between the Tgfbr1 and Egfr signaling programs. To determine how the Egfr dependent motility genes influenced the induction of the trailblazer state, we tested the function of the receptor tyrosine kinase Axl since it can be directly regulated by Fra1 (60). The Axl inhibitor BGB324 suppressed C3-TAg organoid invasion, demonstrating that a gene positively regulated by combined Tgfbr1 and Egfr activity was necessary for trailblazer invasion **(Figure 8F-G)**. Together, these results suggested that the induction of the trailblazer state involved a compromise between the Tgfbr1 and Egfr regulated gene expression programs that limited the expression of genes involved in ECM reorganization and metastatic potential to promote cell motility that was essential for trailblazer cell invasion **(Figure 8H)**.

**Figure 8 with 1 supplement.**
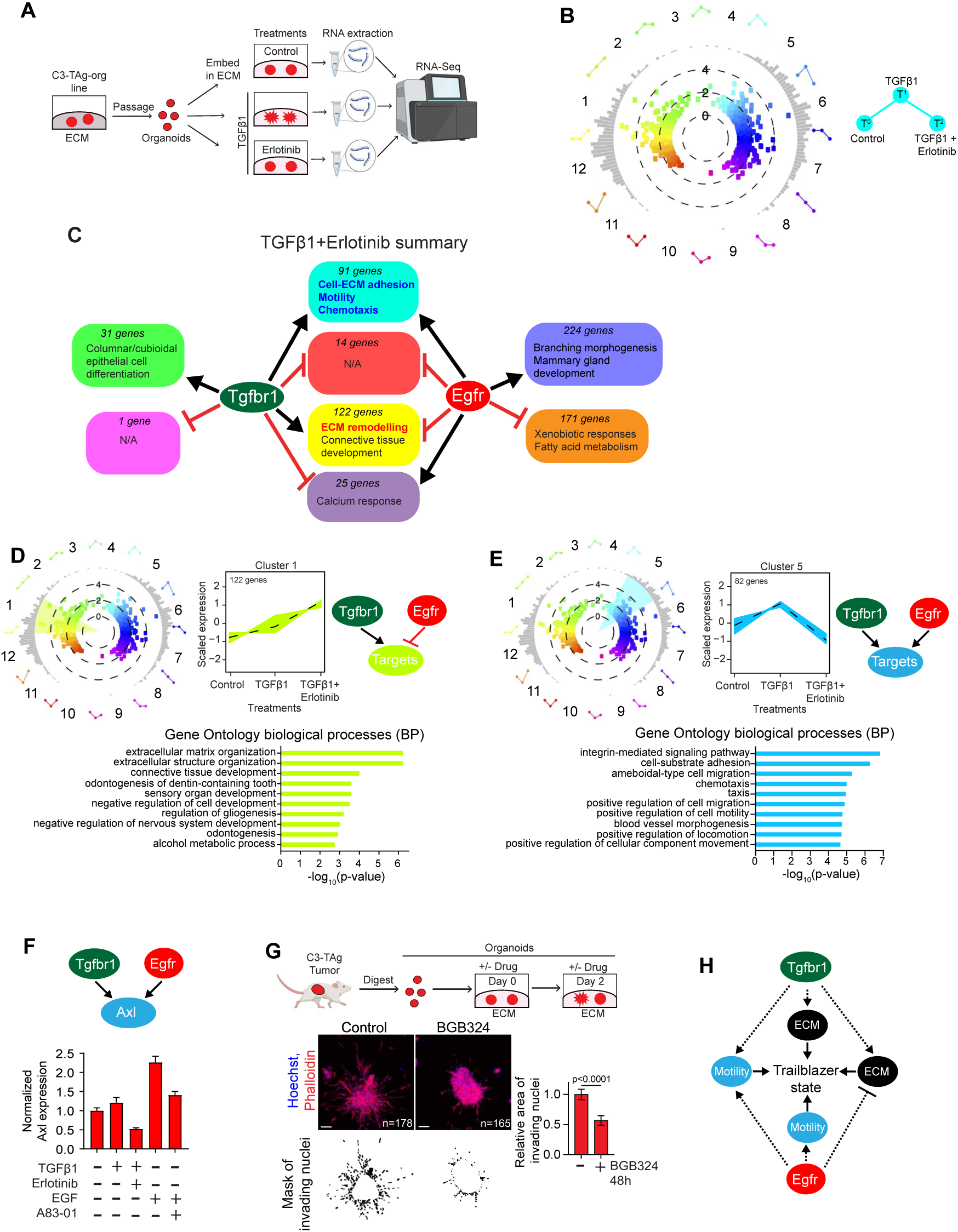
A compromise between the Tgbβ and Egfr signaling programs confers the trailblazer state. **A.** Overview of the approach to test the interactions between the Egfr and Tgfβ signaling pathways. **B.** Polar plot organizing genes represented as individual dots based on their change in expression in response to treatment with TGFβ1 or co-treatment with TGFβ1 and erlotinib. Genes that underwent ≥2 fold-change with a *p*< 0.05 after Benjamini-Hochberg correction in at least one condition relative to one of the other conditions are shown (3 total comparisons). The distance from the center of the plot represents the Δlog_2_ value of each gene (increased difference in expression, further from the center). The bar plots on the surface of the circle show the number of genes at the degree point in the plot. Genes are grouped into 12 color-coded groups along 30 degree increments over the 360-degree plot. As an example, genes in cluster 5 are increased in expression by TGFβ1 and this increase in expression is inhibited by Erlotinib. In contrast, genes in cluster 1 are increased in expression by TGFβ1, and this induction is further enhanced by treatment with Erlotinib. **C.** Summary of the number of genes and represented biological processes associated with different modes of interaction between the Tgfbr1 and Egfr pathways in cells treated with TGFβ1 or TGFβ1+erlotinib. Also see **Figure 8—figure supplement 1C-D** and **Supplementary file 5** for the assignment of polar plot clusters and detailed biological processes used to define the summary integration groups. **D.** Egfr suppressed the expression of a subset of genes induced by extrinsic TGFβ1. Line plots indicate the expression of genes in Cluster 1. Dashed line indicates the average scaled expression of all Cluster 1 genes. Lower bar graph shows the top 10 biological processes (GO-BP) associated with Cluster 1 genes (Fisher’s exact test). **E.** Egfr was necessary for the expression of a subset of genes induced by extrinsic TGFβ. Line plots show the expression of genes in Cluster 5. Dashed line indicates the average scaled expression of all Cluster 5 genes. Lower bar graph shows the top 10 biological processes (GO-BP) associated with Cluster 5 genes (Fisher’s exact test). **F.** Graph of Axl expression (mean±SEM, n=2). **G.** The Axl inhibitor BGB324 (1 µM) suppressed freshly isolated C3-TAg tumor organoid invasion. Graph shows the quantification of invasion (mean±SEM, Mann Whitney test, n=organoids, r=2). **H.** Model showing how the induction of the trailblazer state involves a compromise between the Tgfβ and Egfr regulatory programs. The Egfr dependent induction of genes, such as Axl, that are required for trailblazer cell motility has the cost of restricting the expression ECM remodeling genes known to promote invasion and metastasis.

## DISCUSSION

Our findings indicate that defining how cells metastasize requires an understanding of how trailblazer cells are induced in different breast tumors **(Figure 9)**. The Tgfβ regulated trailblazer program that we have defined in the basal-like C3-TAg mammary tumor cells induced a more aggressive form of collective invasion than was detected in luminal B PyMT tumors and functioned independently of increased Krt14 and p63 expression. This indicated that the Tgfβ program was mechanistically and phenotypically distinct from a previously described Krt14-high trailblazer program (20). In addition, PyMT tumor cells rarely expressed detectable Vimentin, which suggested that the Tgfβ regulated trailblazer program that we identified was not active in PyMT tumors. Thus, our results indicate that there are at least 2 different trailblazer programs and that their activity and requirement for invasion varies between different types of mammary tumors associated with previously defined breast cancer intrinsic subtypes **(Figure 9)**. Genes induced as part of the C3-TAg trailblazer program are more frequently detected at a high level in basal-like breast cancer patients. Vimentin expression also correlated with collective invasion in TNBC patients, which are most often classified as basal-like. Vimentin expression was detected in basal-like TNBC PDX metastases as well. In addition, genes induced as part of the Tgfβ trailblazer program have previously been shown to correlate with reduced odds of survival in TNBC, but not patients classified as ER-positive or HER2-positive (21). These combined results indicate that the Tgfβ trailblazer program is active in basal-like TNBC and influences patient outcome. Tumor subtype specific mechanisms for inducing a trailblazer state are potentially a general property of cancer progression, as molecular subtypes are a feature of many carcinomas (61-64). Further investigation is required to determine the extent to which the Tgfβ and Krt14 trailblazer programs align with existing tumor subtypes or define new categories of inter-tumor heterogeneity. Thus, our results provide new insight into approaches used to stratify patients into outcome groups.

**Figure 9.**
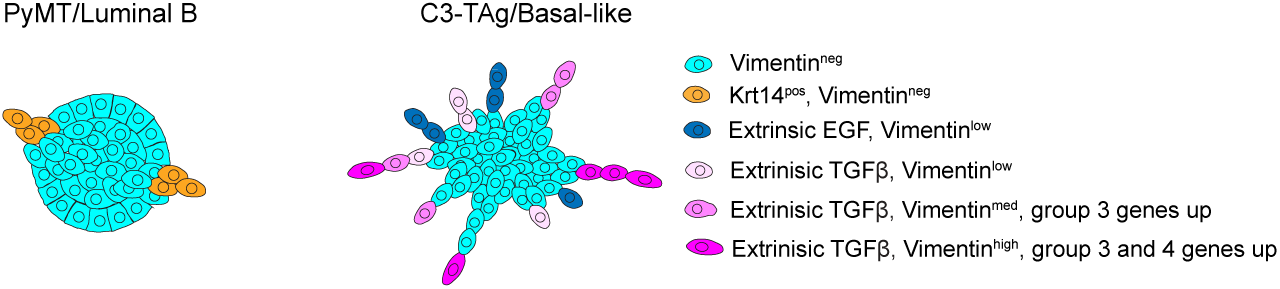
Model of inter- and intra-tumor trailblazer cell heterogeneity. The characteristic features and functional requirements of the trailblazer cells induced by Tgfβ and Egfr signaling programs in C3-TAg/basal-like tumors are fundamentally distinct from the trailblazer cells in PyMT/luminal B tumors.

The heterogeneity in C3-TAg tumor organoid invasion indicated that the trailblazer population itself can be heterogeneous within tumors with the same primary driving genetic factors and molecular subtype **(Figure 9)**. Previous analysis of other tumor cell models had suggested that trailblazer cells were in a single uniform state (20, 21, 24). Notably, our analysis of 1339-org cells revealed that invasive variability can be a consequence of differences in cellular responses to a single stimulus, in this case TGFβ signaling. Therefore trailblazer cell phenotypic heterogeneity within a population did not necessarily reflect differences between EMT programs that were initiated by different activating signals (65). Gene expression changes in response to TGFβ are heterogeneous and temporally regulated in other systems as well (66). However, the influence of this gene expression heterogeneity on cell phenotype has not been clear and has often been modeled as a progressive accumulation of new functions (67). The phased changes in gene expression we defined in 1339-org cells suggested how trailblazer phenotypes evolved over time in response to TGFβ activity, with early onset gene expression changes inducing trailblazer cells that were further modulated by later stage reprogramming events **(Figure 9)**. Thus, differences between tumor cells in the duration of TGFβ pathway activation may contribute to the heterogeneity we detect in trailblazer populations. Interestingly, long-term TGFβ induced changes in gene expression that were associated with other cell lineages. This suggests that the enhanced invasion of trailblazer cells in response to persistent TGFβ stimulation may be associated with a more general lineage plasticity and not simply a re-balancing of epithelial and mesenchymal gene expression.

Our gene expression analysis and functional studies have begun to unravel the details of the transcriptional regulation of the trailblazer state. We detected an increase in Fra1, Zeb1 and Zeb2 expression in C3-TAg trailblazer cells and human breast cancer trailblazer cell lines, with Fra1 regulated by Egfr signaling and Zeb1 and Zeb2 regulated by Tgfβ. Fra1 and Zeb1 were required for the initial Tgfβ induced induction of a trailblazer invasion as well as invasion of cells after they had converted to trailblazer state. Interestingly, Zeb2, which showed a delayed induction in response to Tgfβ, was only required for the invasion of a trailblazer cell line with the characteristics of long-term Tgfβ pathway activation. These results suggest that the transcriptional regulation of trailblazer populations may be different depending on the specific trailblazer state. We also determined how siRNAs targeting 10 additional transcription factors, including the canonical EMT TFs Snail (Snai1), Slug (Snai2) and Twist1, influenced trailblazer cell invasion. The siRNAs targeting the 10 TFs did not yield a clear suppression of invasion and we did not detect a correlation between the increased expression of the TFs and the induction of a trailblazer state. However, our results do not exclude the possibility that these TFs contribute to the Tgfβ dependent induction of a trailblazer state in some context.

Our results revealed that a new compromise mode of integration between Tgfbr1 and Egfr regulated gene expression programs. The integration of parallel signaling pathways is frequently required to confer specific cell states and is typically classified from the standpoint of being a positive or negative interaction (49, 68, 69). Indeed, prior studies have suggested that Tgfβ and Egfr-Ras regulated pathways positively integrate at the point of EMT-TF expression to initiate an EMT program (48, 49, 70, 71). While the combined activity of the Tgfbr1 and Egfr signaling pathways was necessary to induce a trailblazer state, there was an equivalent degree of positive and antagonistic interaction between the programs in C3-TAg tumor cells. This finding revealed a new type of integration between the two signaling programs in which the amplitude of a subset of the gene expression changes induced by either pathway alone are compromised. The selective activation and restriction of Tgfβ and Egf dependent gene expression suggests that compromise integration is dictated by differences in the expression or activity of co-factors that modulate chromatin accessibility or DNA binding. Tgfβ can increase Egfr expression and activate Erk1/2 signaling in a cell line dependent manner (72, 73). Similarly, Egfr-Mek1/2-Erk1/2 signaling can activate Tgfβ signaling through the phosphorylation of SMAD2/3 and increased expression of Tgfβ expression depending on cell context (43, 74, 75). However, this manner of integration is unlikely contributing to the compromise integration in the C3-TAg tumor cells given that Tgfbr1 inhibition did not alter ERK1/2 phosphorylation or Fra1 expression and Egfr and Mek1/2 inhibition failed to impact Smad2/3 phosphorylation. The direct control of the activity of one pathway by the other would also be expected to yield a more uniform positive or negative integration. Defining the mechanistic basis of selective antagonism of gene expression by the Tgfβ and Egfr pathways is an important line of future investigation.

The requirement of Tgfβ and Egfr pathway activity for trailblazer cell invasion indicated that the compromise mode of integration was a feature of the trailblazer state. Egfr pathway stimulation can also promote the motility of non-trailblazer cells, raising the possibility that Egfr signaling has distinct functions in different subpopulations of invading cells, with the precise regulatory contribution dictated by Tgfβ pathway activity (76, 77). The requirement of Tgfβ and Egfr signaling for the collective invasion of MCF10A-Trailblazer and MCFDCIS cells indicates that the integration of the pathways is essential for the induction of a trailblazer state in multiple contexts. However, the Egfr ligand Epgn restricts the invasion of luminal B PyMT organoids, potentially due to the ability of Epgn to suppress the expression of EMT-TFs and ECM reorganization genes in PyMT cells (78). This result suggests that integration of Egfr signaling is not essential for the induction of Krt14 specified trailblazer cells, like those detected in PyMT tumors (20). The differences in how Egfr signaling influences invasion may be a consequence of how tumor cells obtain the ability to resist TGFβ induced growth suppression and apoptosis (79). This is accomplished either through the clonal selection of mutants that inactivate Tgfβ signaling, which does not occur in C3-TAg tumor cells, or an intrinsic reprogramming of cellular responses to TGFβ (80). The type of reprogramming to TGFβ stimulation that is selected for during tumor progression may determine whether Tgfbr1 and Egfr signaling engage in a compromise mode of integration or if there is a predominantly positive or negative interaction between the pathways.

Our findings indicate that defining the nature of Tgfβ and Egfr pathway integration is necessary to understand tumor cell properties that influence metastasis and treatment response. In C3-TAg cells in which Tgfβ and Egfr activity combined to promote a trailblazer state, Egfr also restricted the expression of genes that could endow trailblazer cells with additional invasive and metastatic properties (53, 81-84). Similarly, Tgfβ limited the Egf induced expression of chemokines that increase invasion and the recruitment of tumor promoting immune cells (85, 86). Thus, the compromise mode of integration limited the expression of genes that could promote tumor progression whereas a strictly positive integration would have no such restrictions. The type of interaction between the Tgfβ and Egfr programs also influences the rationale for the therapeutic targeting of trailblazer cells. Inhibiting the Egfr pathway with the clinically approved drugs erlotinib and trametinib suppressed C3-TAg tumor cell invasion (87-89). However, combining either drug with newly developed anti-Tgfβ antibodies (90) may have additional benefit due to the fact that the Egfr and Tgfβ pathways function in parallel and each independently regulate large cohorts of genes. By comparison, cells in which Tgfβ activates Egfr-ERK1/2 signaling or vice versa may derive little or no benefit from combination therapy (72, 73). Thus, understanding how the mode of Tgfβ and Egfr pathway integration influences invasion, the risk of metastasis and options for treatment is an important line of future investigation.

Together, our results indicate that an improved understanding of how trailblazer cells are regulated and function in different breast cancer subtypes has the potential to improve the accuracy of patient diagnosis and reveal new treatment options.

## MATERIALS AND METHODS

### Mice

The C3-TAg [FVB-Tg(C3-1-TAg)cJeg/Jeg], PyMT [FVB/N-Tg(MMTV-PyVT)634Mul/J] and NOD.Cg-Prkdcscid Il2rgtm1Wjl/SzJ (NSG) mice were purchased from The Jackson Laboratory (C3-TAg: 013591, PyMT: 002374 and NSG: 005557) and (**Supplementary file 1**). Mice were housed, bred and euthanized in accordance with a protocol approved by the Institutional Animal Use and Care Committee at Georgetown University (IRB# 2017-0076) and in compliance with the NIH Guide for the Care and Use of Laboratory animals. Female mice were used for all experiments.

### Patient derived xenografts

Triple negative breast cancer (TNBC) PDX fragments (**Supplementary file 1**) were obtained from Dr. Alana Welm at University of Utah (39). A single fragment of HCI-001 PDX was transplanted into the cleared 4^th^ mammary fat pad of 4 weeks old NSG female mice. Tumor measurements were taken every week to monitor tumor growth. All surgical procedures and euthanasia was performed in accordance with IACUC approved protocols at Georgetown University. Tumor pieces and lungs were fixed in formalin followed by paraffin embedding and sectioning by Histology and Tissue Shared Resource (HTSR) at Georgetown University.

### Tumor organoid derivation

The largest tumors from female C3-TAg and PyMT mice were minced and dissociated for up to 120 min at 37°C in a mixture of 1 mg/ml Collagenase (Worthington, LS004188), 20 U/ml DNase (Invitrogen, 18047-019), 5% Fetal Bovine Serum (FBS) (Peak Serum, PS-FB2), ITS (Lonza, 17-838Z), non-essential amino acids (Sigma, M7145) and 10 ng/ml FGF-2 (Peprotech, 100-18C) in DMEM/F-12 (Corning, 10-092-CV). Dissociated tumors were pelleted at 80 x *g* for 2 min and the supernatant was discarded. Tumor organoids were then rinsed up to 5 times in 5% FBS in DMEM/F-12 followed by filtering through a 250 µm tissue strainer. Organoids were then tested in invasion assays or to establish organoid lines and cell lines (**supplementary file 1**).

### Cell and organoid culture

Cell and organoid lines were generated from C3-TAg tumors **(supplementary file 1)** and tested for mycoplasma (Lonza, LT07-703) prior to use and the creation of frozen stocks. Cells and organoids were routinely used within 25 passages. Culture conditions for all cell and organoid lines are summarized in **Supplementary file 1.** Organoids were passaged at least once per week by dissociating tumor organoids into single cell suspensions with Dispase (Sigma, SCM133) and TryPLE (Gibco, 12605-010). The cell suspensions were plated at a density of 200,000-500,000 cells per well in a 24-well ultra-low adhesion plate (Sigma, CLS3474) to form multicellular aggregates for at least 16 h. The aggregates were then embedded in Matrigel (Corning, 354230) or Cultrex (Biotechne, 3533-005-02) and overlaid with Organoid Media (**Supplementary file 1**) in 6-well ultra-low adhesion plates (Sigma, CLS3471). H2B:GFP retrovirus (91) and H2B:mCherry and GFP lentivirus were produced and cells were infected as described previously (77).

### Orthotopic Tumor Models

For orthotopic transplantation experiments, 1339-org cells were first cultured in Organoid media (Control) or Organoid media without A83-01 and supplemented with 2 ng/ml TGFꞵ1 (+TGFꞵ1). After 21 days of treatment, 500,000 control or +TGFꞵ1 1339-org cells were injected in the right 4^th^ fat pad of 8-12 weeks old female NSG mice. Seven mice per group from 2 independent experiments were analyzed. Mice were euthanized and tissue collection was performed 34 days post injection. Representative portions of primary tumors and complete lungs were fixed in formalin followed by paraffin embedding and sectioning by Histology and Tissue Shared Resource (HTSR) at Georgetown University. The remaining primary tumor portions were used for new organoid derivation and analysis.

### Blood collection and Circulating Tumor Cell (CTC) isolation

For CTC isolation, mice bearing Control or +TGFꞵ1 1339-org orthotopic tumors were anesthetized with isoflurane and 800 µl of blood was extracted via cardiac puncture using a 25 G needle (BD, 305122). The blood was immediately transferred to EDTA coated micro tubes (Sarstedt, 4111395105) to avoid coagulation. Blood collected from all mice from the same treatment group (Control or +TGFꞵ1) were pooled together, mixed with red blood cell (RBC) lysis buffer (Thermo Fisher, 00433357) and incubated on ice for 10 min. The samples were then spun down at 4°C for 10 min at 250 g. The supernatant was discarded and the same steps were repeated once more to completely remove the RBCs. The cell pellet was then resuspended in cold 1X PBS and plated in poly-l-lysine pre-coated 8 well chamber slides (0.4 mL per well). The slide was placed in the 37°C incubator for at least 30 min. The cells were then fixed, permeabilized, blocked and stained with anti-Krt8 antibody to detect circulating C3-TAg tumor cells.

### Organoid invasion

Organoids in 30 µl of ECM were plated onto a 20 µl base layer of ECM in 8 well chamber slides (Falcon, 354108) and allowed to invade for 48 h unless otherwise indicated. The ECM was a mixture of 2.4 mg/ml rat tail collagen I (Corning, CB-40236) and 2 mg/ml of reconstituted basement membrane (Matrigel or Cultrex) unless otherwise indicated. Tumor organoids were analyzed in a base media of DMEM/F-12 and ITS in **Figure 1A, D, E and G, Figure 1—figure supplement 1B, Figure 4B, C and G, Figure 5G, Figure 7G and Figure 8G**, and 1% FBS + 10 ng/ml FGF2 in **Figure 1B and F**. A base media of DMEM/F-12 and ITS was used for the analysis of 1339-org, 1863-org and 1788-org lines. Growth factors (2 ng/ml TGFβ1; 50 ng/ml EGF) and inhibitors were added to the base media and tested as indicated in the text and **Supplementary file 1**. Organoids from established organoid lines **(Supplementary file 1)** were tested after dissociation and aggregation for at least 16 h as described for organoid passaging. Organoids were fixed and stained with Hoechst and Phalloidin. Images were acquired with a Zeiss LSM800 laser scanning confocal microscope using 10x/0.45 (Zeiss, 1 420640-9900-000) or 20x/0.8 (Zeiss, 1 420650-9902-000) objectives. To determine the area of invasion and circularity, an image mask for each organoid was generated using ImageJ (NIH) based on the Hoechst (invasion area) or Phalloidin (circularity) signal. The total area of the invading cell nuclei was determined using the “Measure” function in ImageJ. Circularity was quantified using the “Analyze Particle” function.

### siRNA experiments

Cells were transfected with 50 nM of siRNA using RNAiMax transfection reagent (Invitrogen, 13778-150) for 48 h before testing. Cells in all conditions designated as “Control” were transfected with a pool of siRNAs that does not target mouse genes. The catalog numbers for each siRNA are located in **Supplementary file 1**.

### Vertical invasion assay

Vertical invasion assays were performed and quantified as described previously using a Collagen/Basement Membrane (BM) mix (2.4 mg/ml Collagen I and 2 mg/ml Matrigel or Cultrex) (25). Cells expressing H2B:GFP (10,000-18,000/well) were transfected in 96-well plates (Thermo, 165305) for 48 h. Media was then replaced with 50 µl Collagen/BM) ECM and overlaid with fresh media (DMEM-F12, ITS, 1% FBS, 10 ng/ml FGF2) for 48 h. Cells were imaged with a 10X/0.30 objective (Zeiss, 1 420340-9901-000) on a Zeiss LSM800 laser scanning confocal microscope. Twenty-one z-slices at 10 µm intervals over a total span of 200 μm were acquired for 9 tiled x, y positions per well. At least 3 wells (27 x,y positions total) were imaged per condition in each experiment. ImageJ software (NIH) was used to process images and quantify invasion using a custom macro as described (21). Invasion was normalized by dividing the total number of cells that had migrated ≥ 40 µm above the monolayer by the number of cells in the monolayer. The normalized invasion value for each condition was divided by the normalized invasion for the control wells transfected with an siRNA pool that does not target any mouse genes to provide a relative invasion value.

### Cluster invasion

Cells (10,000-18,000/well) were transfected in 96-well plates (Corning, 3904). After 24 h, the transfected cells were trypsinized and clustered in 96-well Nunclon Sphera U-bottom low adhesion plates (Thermo Scientific, 174925) overnight (1000 cells/well). Homogeneous and mosaic clusters were also generated in U-bottom Nunclon Sphera low adhesion plates. Cells then aggregated into clusters for at least 16 h before embedding in ECM. Clusters were resuspended in 30 μl Collagen I/BM mix (2.4 mg/ml Collagen I and 2 mg/ml Matrigel or Cultrex), plated on 20 μl of a base layer of Matrigel/Collagen I, overlaid with media containing 1% FBS and 2 ng/ml TGFβ1 and allowed to invade for 24 h-48 h unless stated otherwise. The spheroids were fixed and stained with Hoechst and Phalloidin and imaged with a 10x/0.45 objective (Zeiss, 1 420640-9900-000) using a Zeiss LSM800 laser scanning confocal microscope. To determine the area of invasion, Hoechst staining was used to identify and mask nuclei invading into the ECM using ImageJ. The total area of the invading nuclei was then determined using the “Measure” function in ImageJ.

### Live cell imaging

Imaging was performed using a Zeiss LSM800 laser scanning confocal microscope enclosed in a 37° C chamber supplemented with humidified CO_2_. Images were acquired every 30 min for 14-18 h with a 10X/0.30 objective (Zeiss, 1 420340-9901-000). At least 15 different x,y coordinates with 5 to 7 z-slices over 50-100 μm span for each condition were acquired in parallel.

### Immunofluorescence and immunoblotting

Experiments and analysis were performed as described (25) using antibodies detailed in **Supplementary file 1.** Immunoblots were imaged using an Odyssey scanner (Licor, 9120). Immunofluorescence images were acquired using 10x/0.45 (Zeiss, 1 420640-9900-000) and 20x/0.8 (Zeiss, 1 420650-9902-000) objectives. For analysis of the relationship between Krt14 expression and invasion, organoids were classified as invasive if they had at least two protrusions with each protrusion having at least two cells invading outward from the center mass. Images were exported as TIFFs and analyzed with ImageJ. The freehand selection tool was used to define ROIs containing individual organoids for analysis. The Krt14 fluorescence for each organoid was then defined as the “Mean Gray Value”, as determined using the “Measure” function. To define Krt14 expression in cells leading invasion, >10 organoids stained with Phalloidin, Hoechst, Krt14 and Krt8 were analyzed. Seven Z slices at 10 µm intervals over a total span of 100 µm were acquired, and the leading cells of each protrusion were designated as Krt14^low^ or Krt14^high^. Vimentin and E-cadherin expression were analyzed in 5 µm histology sections. Organoids in 8-well chamber slides (Falcon, 354108) were washed 3x with distilled water. After detaching the chamber walls from the slides, the organoid/ECM gels were gently picked up using a single edge blade (Personna, 94-120-71) and laid out on a strip of parafilm (Bemis, PM-999). The 8-well chamber walls were placed around the gels and 150-200 µl of hydroxyethyl agarose processing gel (Thermo Fisher, HG 4000-012) was poured into the well. After the hydroxyethyl agarose processing gel solidified, it was transferred into histosettes (Simport, M498-3) and stored in 70% EtOH until embedding in paraffin by the Georgetown Histology and Tissue Shared Resource. The 5 µm sections were deparaffinized and immunostained as described (21) to detect Vimentin and E-cadherin expression (**Table S1**). For individual organoids, Vimentin expression was defined as the “Mean Gray Value” using the “Measure” function in ImageJ. Using ImageJ, Vimentin expression was defined as the “Mean Gray Value” for individual organoids using the “Measure” function. For characterization of the trailblazer cell state, the cell leading a collectively invading group of cells was classified as a “trailblazer cell” while cells detached from the tumor organoid were classified as “single cells”. Each cell was assigned E-caderhin^pos^/ Vimentin^neg^, E-Cadherin^pos^/Vimentin^pos^ or E-Caderin^neg^/ Vimentin^pos^ status based on E-Cadherin and Vimentin expression.

### Immunohistochemistry

Sample processing and staining was performed as described (25). Images were acquired using 10x/0.45 (Zeiss, 1 420640-9900-000) and 20x/0.8 (Zeiss, 1 420650-9902-000) objectives. At least 10 regions of interest (ROIs) were randomly selected for each tumor. Cell Profiler (Broad Institute) was used to define image masks based on Krt8, Krt14 or Vimentin signal. To define their expression in tumor cells, masks based on the area of overlap between the Krt14 and Krt8 and the Vimentin and Krt8 signals were determined for each ROI. The area of individual channel and overlap masks created in CellProfiler were then loaded onto ImageJ and quantified using the “Measure” function.

### Quantitative real-time PCR

Total RNA was isolated using the RNeasy Kit (Qiagen, 74104) or the PureLink™ RNA Mini Kit (Invitrogen, 12183025). Reverse transcription was performed using iScript cDNA Synthesis Kit (Bio-Rad, 1708891) following the manufacturer’s protocol. Real time PCR was performed with iQ SYBR Green Supermix (Bio-Rad, 1708882) using a RealPlex 2 thermocycler (Eppendorf, 2894). All primers were designed and purchased from Integrated DNA Technologies and their sequences are listed in **Supplementary file 1**.

### Flow Cytometry

Cells were dissociated with TrypLe (Gibco, 12605-010) into single cell suspensions, and then washed 2x with FACs buffer (PBS + 2% FBS). Cells were then incubated with anti-CD326 (EpCAM)-PE/Dazzle 594 (Biolegend, 118236) at 4°C for 30 minutes. After staining, cells were washed with FACs buffer 2x, and stained with Helix NP Blue (Biolegend, 425305) in order to exclude dead cells. Cells were then subjected to flow cytometry analysis using BD LSR Fortessa. Results were analyzed using FCSExpress 7 (De Novo Software).

### RNA sequencing

For the analysis of C3-TAg tumor organoids and non-invasive spheroids in **Figure 2 and Figure 2–figure supplement 1**, at least 1000 C3-TAg tumor organoids or 60,000-80,000 non-invasive cells were embedded in a 500 µl ECM mixture (2.4 mg/ml Collagen I and 2 mg/ml Matrigel or Cultrex) and plated on top of 500 uL base layer of ECM in 6 cm dishes (Falcon, 353004). C3-TAg tumor organoids were allowed to invade for 48 h. Non-invasive cells were allowed to grow and form spheroids for 6 days. Experiments were performed twice using frozen stocks of tumors and cell lines. For the analysis of 1339-org responses to growth factors and inhibitors in **Figure 4, Figure 8 and Figure 8–figure supplement 1**, 250,000 1339-org cells were embedded in a 100 µl ECM mixture (2.4 mg/ml Collagen I and 2 mg/ml Matrigel or Cultrex) and plated on top of a 200 µl ECM base layer in 6 cm dishes (Falcon, 353004). The gels were overlaid with control or treatment media. Organoid/ECM gels were collected in 15 mL conical tubes (Fisher, 339651), snap frozen and stored in −80° C freezer. Experiments were performed twice. For RNA isolation in all experiments, the gels were thawed on ice and disintegrated in 1 mL Trizol (Invitrogen, 15596026) by repeatedly passing the mixture through a 23G needle (BD, 305143). After the gels were homogenized, the Trizol-gel mixture was allowed to sit at RT for 5 min. Then 200 µL of chloroform (Sigma-Aldrich, C2432-25ML) was added. Following vigorous shaking for 15 sec, the mix was left for 2 min at RT to allow for phase separation. The mix was then centrifuged at 12,000 x g for 15 m at 4° C. The upper aqueous phase was then transferred to a new tube and RNA was then further purified using the RNeasy Kit (Qiagen, 74104) and or PureLink™ RNA Mini Kit (Invitrogen, 12183025). For **Figure 2 and Figure 2–figure supplement 1**, RNA sequencing, was performed by the Genomics and Epigenomics Shared Resource at Georgetown University. The cDNA libraries were generated using the Illumina TruSeq Stranded Total RNA Library Preparation Kit (Illumina, 20020597). Sequencing was performed using an Illumina NextSeq 500 at a depth of 12 million paired-end 75 base pair reads per sample. Data is available from GEO at GSE188312. Reads were quantified using Salmon with mouse reference genome (GRCm38/mm10). Counts were normalized and differential expression was calculated using DESeq2. For **Figure 4, Figure 8 and Figure 8–figure supplement 1**, RNA sequencing was performed by Novogene. RNA quality analysis was assessed by Bioanalyzer using the RNA Nano 6000 Assay Kit (Agilent, 5067-1511). The cDNA libraries were prepared and indexed using the NEBNext Ultra RNA Library Prep Kit for Illumina (NEB, E7530L). Libraries were sequenced with the NovaSeq 6000 at a depth of 20 million paired-end 150 base pair reads per sample. Sequencing data available from the GEO at GSE188300 and GSE188323. Reads were mapped to the mouse reference genome (GRCm38/mm10) using HISAT2 and reads per gene determined using HTSeq. Counts were normalized and differential expression was calculated using DESeq2.

### Polar plot analysis

Expression data was normalized and differentially expressed genes with a ≥2-fold change in expression with *p*< 0.05 after Benjamini-Hochberg (BH) correction were identified using the *DESeq2* package in R (92, 93). Normalized expression data was collapsed for all genes as the average between condition replicates and were subset to only include genes that were differentially expressed between at least one of the three conditions (control, growth factor, growth factor plus inhibitor) assessed. To define the integration between the pathways, we modified a previously described approach for generating polar plots that has been used to identify dynamic trends in gene expression (47, 94). Relative mean signal changes over successive conditions, of control, growth factor, and growth factor plus inhibitor treatment (T°, T^1^, and T^2^, respectively), were transformed to an angle between −180 and +180 degrees (θ). This angle was further transformed to a scale from 0 to 360 degrees and then interpolated into a color based on the angle. Differentially expressed genes were plotted using polar coordinates (*r*,θ) with the radius (*r*), or distance from the focus, being equal to the value of largest absolute log_2_ fold change between conditions and theta (θ) being equal to the angle corresponding to the trend of change and calculated using the following equations:

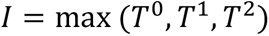

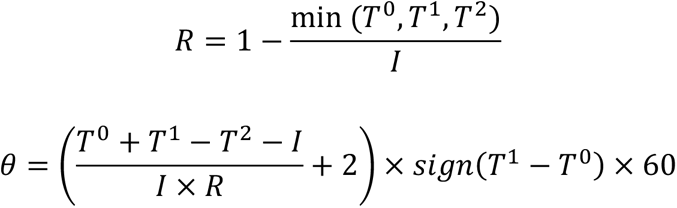

The intensity (*I*) is equivalent to the maximum expression for a gene across conditions and the ratio (*R*) between the minimum expression of a gene across conditions over the intensity, are used to calculate the angle of expression trend (θ). As such, genes are plotted as a point with an angle (θ) corresponding to relative changes in a gene’s expression with subsequent treatments at a distance from the focus corresponding to the largest magnitude of change between any pair of conditions in the treatment series. Three-point profile annotations of expression change (corresponding from left to right as T°, T^1^, and T^2^, respectively) are indicated on the outer edge of the polar scatter plots every 30 degrees in the color assigned to that particular θ. These profiles are accompanied by a radial histogram denoting the density of differentially changed genes across organoid conditions at particular θ of the polar scatterplot.

To summarize the meaning of genes with particular profiles in the polar scatterplots, genes were grouped by profiles at 30-degree intervals starting at 0 degrees, which denotes genes that were unchanged in expression from T° to T^1^ but demonstrated a significant increase in expression following growth factor plus inhibitor treatment of the organoids (T^2^). Genes from each group were assessed for enrichment of Gene Ontology biological processes (GP) using the EnrichGo function of the *clusterProfiler* package in R (95). Processes with *p*< 0.01 after BH correction are included in **Supplementary files 4 and 5**. Values were transformed (-log_10_) and displayed alongside polar scatterplots. Genes within each profile group of the polar scatterplot and utilized for GO enrichment were scaled and displayed as a line plot. A dotted line was added to these linear plots to denote the mean scaled expression pattern across organoid treatment conditions for all genes within a particular interval.

### Breast Cancer Tissue Microarray

The construction of the TNBC tissue microarray (TMA) containing duplicate cores from 50 patients was performed by the Histopathology and Tissue Shared Resource (HTSR) at Georgetown University. Details of the methodology, patient consent, de-identification and demographics have been previously described (96). Multiplex staining of deparaffinized and rehydrated cores was performed through sequential rounds of staining, stripping and re-probing with antibodies recognizing KRT14, Vimentin and E-cadherin and OPAL fluorescent secondary antibodies **(Supplementary file 1)**. The TMA cores were imaged with a Vectra3 Multi-Spectral Imaging microscope (Perkin Elmer) and analyzed with inForm software (Perkin Elmer). Tumor and stroma were segmented based on E-cadherin expression (tumor). Cores with folded sections or no clear tumor-stomal interface by manual inspection were excluded from analysis. Cores were classified “high invasion” if invasive strands were detected in >50% of the tumor-stomal interface by manual inspection. The KRT14 and Vimentin signals were not visible when classifying invasion status.

### Human Breast Cancer Gene Expression Analysis

The mRNA expression data for 996 patients in the Breast Invasive Carcinoma (The Cancer Genome Atlas (TCGA), PanCancer Atlas) was analyzed and downloaded from the cBioPortal (97, 98). Tumor subtype specific gene expression values are Z-scores relative to all samples. Correlative expression values are Log2 transformed. The mRNA expression for human breast cancer cells was derived from a prior analyses (21, 31) and available at the GEO (GSE58643 and GSE62569).

### Statistical Methods

Data was analyzed with Prism 9.0.2 (Graphpad) and *DESeq2, GSEA* and *ClusterProfiler* packages in R. Data with a normal distribution, as determined by Shapiro-Wilk test, was analyzed by two tailed Student’s t-test. Data that did not pass a normality test were analyzed by Mann-Whitney U test or Kruskal Wallis test with Dunn’s Multiple comparison test. Differentially expressed genes with *p*< 0.05 after Benjamini-Hochberg correction were identified using the *DESeq2* package in R. Fisher’s exact tests to determine the enrichment of Gene Ontology biological processes (BP) were performed using the EnrichGo function of the *ClusterProfiler* package in R. The specific statistical tests are indicated in the figure legends. Sample numbers are defined and indicated in the figure legends or on the figure panels. All experiments include a minimum of 2 biological replicates from separate biological preparations.

## Data Availability

The RNA sequencing data generated in this study are publicly available at the GEO at GSE188300, GSE188312, GSE188323, GSE58643 and GSE62569. All other data are available within the article and its supplementary data files.

## Materials Availability

Cell and organoid lines derived from C3-TAg tumors will be made available by the corresponding author upon reasonable request.

## AUTHORS’ CONTRIBUTIONS

**A. Nasir:** Conceptualization, investigation, writing-review and editing. **S. Camacho:** Conceptualization, investigation, writing-review and editing. **A.T. McIntosh:** Investigation, writing-review and editing. **G. Graham:** Investigation, writing-review and editing. **R. Rahhal:** Investigation, writing-review and editing. **M.E. Huysman:** Investigation. **F. Alsharief:** Investigation. **A.T. Riegel:** Resources, funding acquisition. **G.W. Pearson:** Conceptualization, investigation, writing-review and editing, funding acquisition.

## ACKNOWLEDGEMENTS

Work was supported by NIH R01CA218670, Georgetown Women and Wine (to G.W. Pearson), NIH T32CA009686 (to S. Camacho, G. Graham) and NIH P30CA051008.

---

**Figure 1—figure supplement 1.**
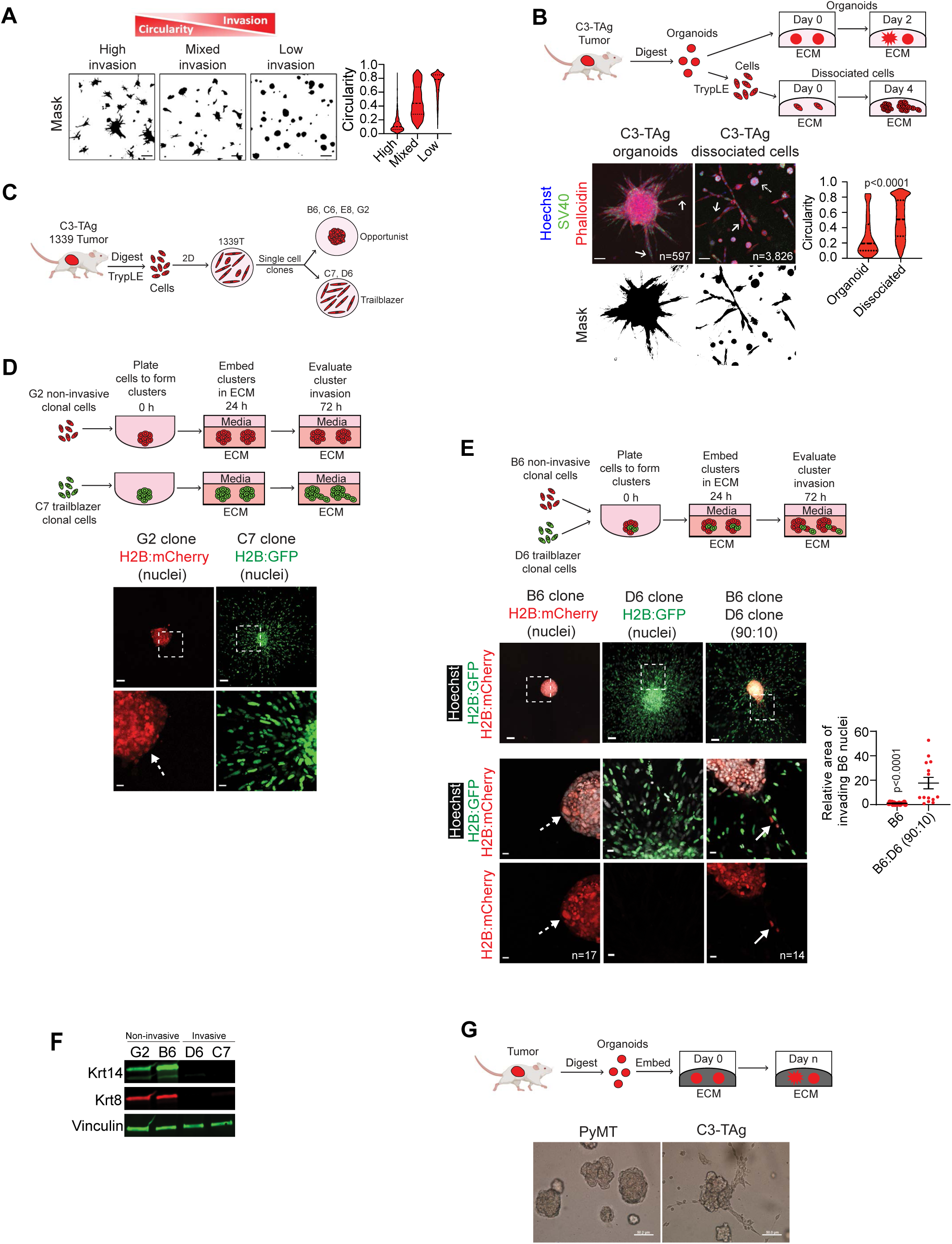
**A.** Representative masked images of C3-TAg organoids showing reduced organoid invasion which is represented by increased organoid circularity. Scale bars, 50 µm. **B.** C3-TAg organoids and dissociated cells from the same tumors were grown in ECM for 2 and 4 days respectively to evaluate invasion. Solid arrows indicate representative invasive cells. Dashed arrow indicates a representative non-invasive spheroid. Violin plot shows the quantification of circularity (Mann Whitney test, n=organoids, dissociated cell spheroids or individual dissociated cells, r=2). Scale bars, 50 µm. **C.** Graphic shows the methodology for deriving opportunist and trailblazer clonal cell lines. **D.** Representative images of non-invasive (G2) and invasive (C7) clonal cell line clusters. Scale bars, 100 µm (top), 20 µm (bottom, inset). **E.** Trailblazer D6-H2B:GFP cells promoted the collective invasion of non-invasive opportunist B6-H2B:mCherry cells. Top row, 2×2 tiled LED fluorescence images. Inset images are confocal sections of the ROIs indicated by the dashed line boxes. Graph shows the quantification of B6-H2B:mCherry cell invasion (mean±SEM, Mann Whitney test, n=clusters, r=2). Scale bars, top row 100 µm; inset images 20 µm. **F.** Immunoblots showing the expression of Krt14 and Krt8 in non-invasive (G2, B6) and invasive (D6, C7) C3-TAg clonal cell lines. Vinculin serves as a loading control. (n=2) **G.** Representative images from 2 independent experiments show that C3-TAg organoids are intrinsically more invasive than PyMT organoids in ECM composed of low Collagen I.

**Figure 1—Video 1. Live imaging of C3-TAg derived tumor organoids in organotypic culture (GFP, green, tumor cell).** Fluorescent and bright field images were acquired for 14 h at 30 min intervals. The video corresponds with invasive heterogeneity in C3-TAg organoids shown in **Figure 1A and Figure 1—figure supplement 1B**.

**Figure 1—Video 2. Live imaging of C3-TAg derived tumor dissociated organoids in organotypic culture.** Bright field images were acquired for 5 h at 30 min intervals. The video corresponds with the intratumor invasive heterogeneity in C3-TAg tumor dissociated organoids shown in **Figure 1—figure supplement 1B**.

**Figure 1—Video 3. Live imaging of homogenous clusters of C3-TAg tumor derived noninvasive clonal cells B6 (H2B:mCherry, red, nuclei).** Fluorescent and bright field images were acquired for 14 h at 30 min intervals. The video corresponds with non-invasive B6 clusters shown in **Figure 1B and Figure 1—figure supplement 1E**.

**Figure 1—Video 4. Live imaging of homogenous clusters of C3-TAg derived tumor cells.** Bright field images were acquired for 14 h at 30 min intervals. The video corresponds with invasive C3-TAg tumor cells clusters shown in Figure 1B.

**Figure 1—Video 5. Live imaging of heterogeneous clusters of C3-TAg derived tumor cells and non-invasive clonal cells B6 (H2B:mCherry, red, nuclei)**. Fluorescent and bright field images were acquired for 14 h at 30 min intervals. The video corresponds with opportunistic invasion of B6 cells induced by C3-TAg tumor cells as shown in Figure 1B.

**Figure 1—Video 6. Live imaging of heterogeneous clusters of C3-TAg derived invasive (D6) and non-invasive (B6) clonal cells (H2B: GFP, green, nuclei; H2B:mCherry, red, nuclei)**. Fluorescent and bright field images were acquired for 14 h at 30 min intervals. The video corresponds with opportunistic invasion of B6 cells induced by D6 cells as shown in Figure 1— figure supplement 1E.

**Figure 1—Video 7. Live imaging of PyMT derived tumor organoids in organotypic culture.** Bright field images were acquired for 18 h at 30 min intervals. The video corresponds with less invasive PyMT organoids as shown in Figure 1F.

**Figure 1—Video 8. Live imaging of C3-TAg derived tumor organoids in organotypic culture.** Bright field images were acquired for 18 h at 30 min intervals. The video corresponds with more highly invasive C3-TAg organoids as shown in Figure 1F.

**Figure 2—figure supplement 1.**
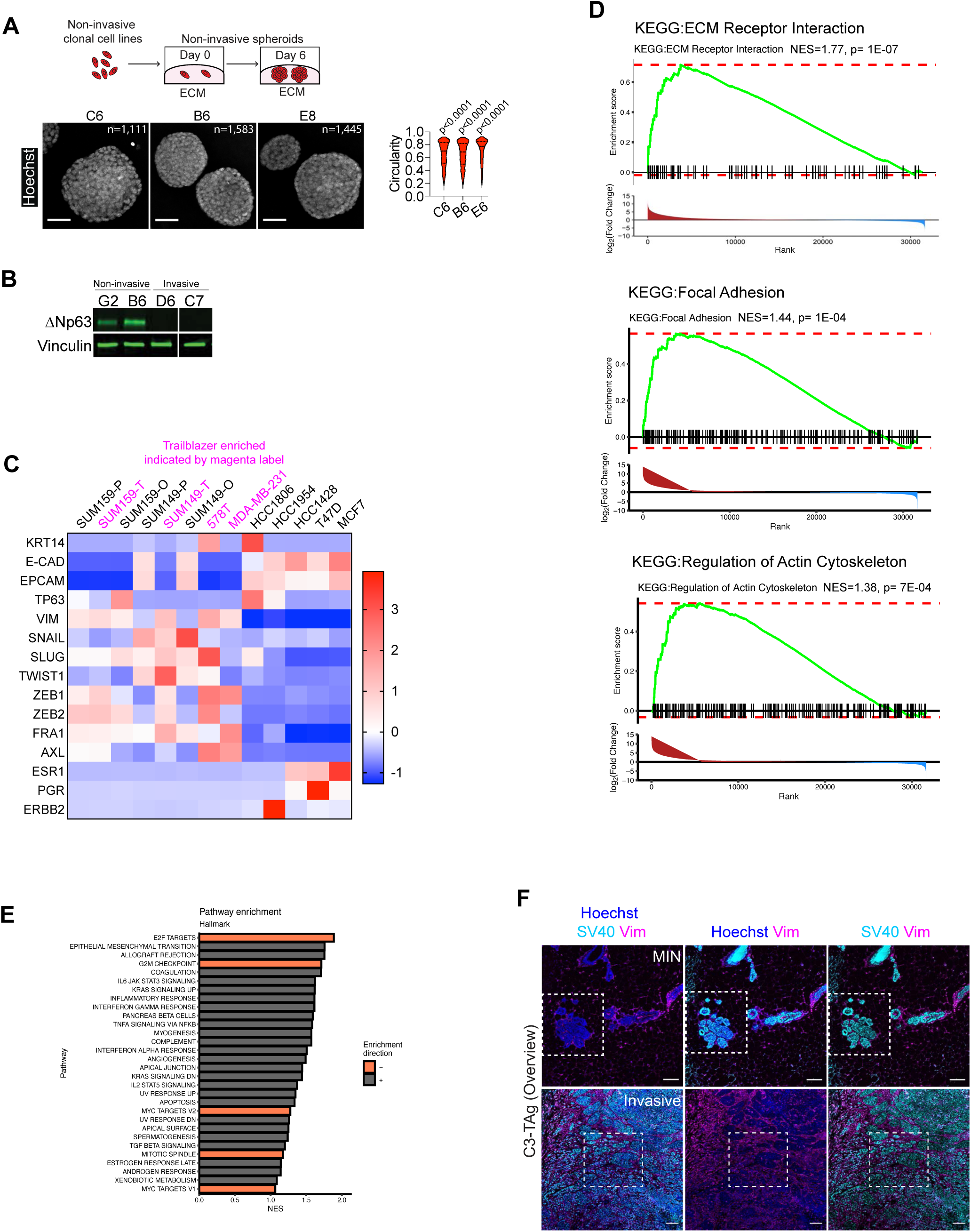
**A.** Non-invasive opportunist C3-TAg clonal cell lines were grown in ECM for at least 6 days. Violin plot shows the quantification of spheroid circularity (Mann Whitney test, n=spheroids, r=2). Scale bars, 50 µm. **B.** Immunoblots showing the expression of ΔNp63 in non-invasive (opportunist) (G2, B6) and invasive (trailblazer) (D6, C7) C3-TAg clonal cell lines (n=2). Lysates are from the same gel. **C.** Heatmap showing the expression of the indicated genes in human breast cancer cell lines. Trailblazer enriched populations are indicated in magenta. P=parental, T=trailblazer and O=opportunist when parental cell lines and their daughter trailblazer and opportunist subpopulations are shown (SUM159 and SUM149). **D.** GSEA was performed on RNA-seq data from C3-TAg organoids and non-invasive C3-TAg clones spheroids to identify pathway enrichment in C3-TAg organoids. Plots show the GSEA for the indicated KEGG pathways. **E.** Graph shows Hallmark pathways with NES ≥1. **F**. Overview images of noninvasive carcinoma in situ (CIS) and invasive carcinoma cells in C3-TAg tumors. Dashed boxes showing the ROIs from Figure 2G. Scale bars, 100 µm.

**Figure 3—figure supplement 1.**
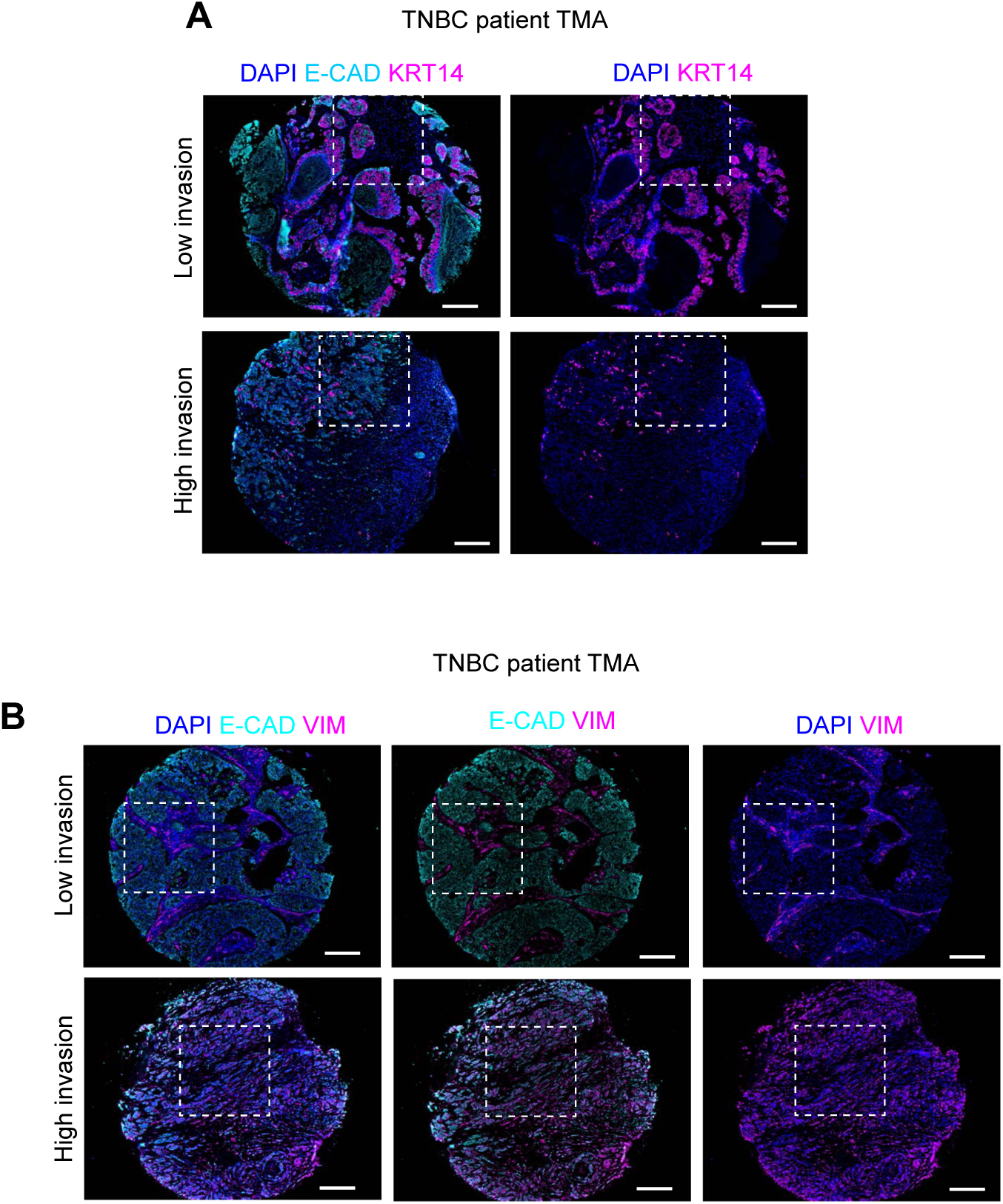
**A.** Full TNBC patient tumors images with dashed boxes showing the ROIs from Figure 3B. Scale bars, 200 µm. **B.** Full TNBC patient tumor images with dashed boxes showing the ROIs from Figure 3D. Scale bars, 200 µm.

**Figure 4—figure supplement 1.**
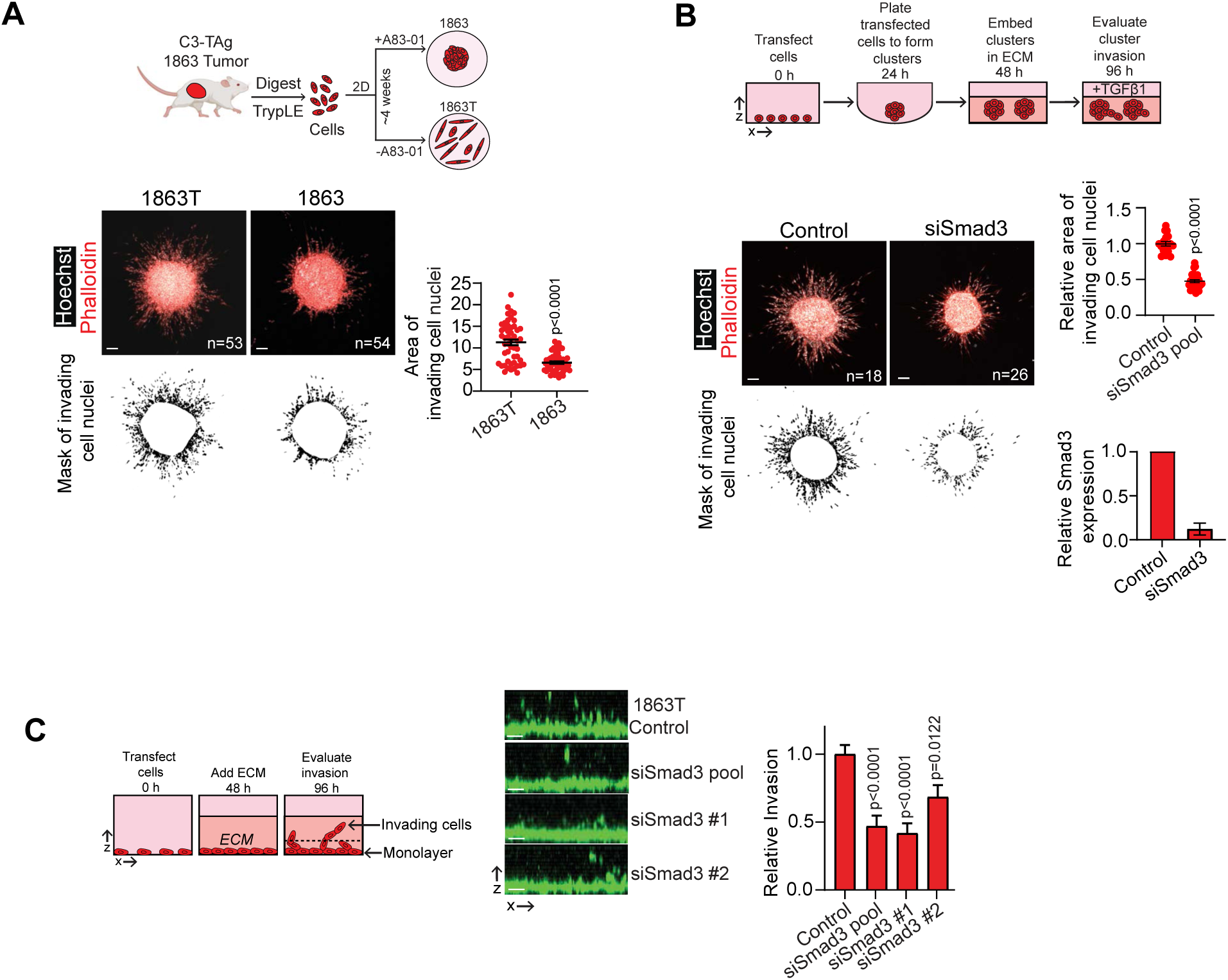
**A.** 1863 cells derived in the presence of the Tgfbr1 inhibitor A83-01 are less invasive than 1863T cells, which were derived from the same tumor but without A83-01. Graph shows 1863 and 1863T cluster invasion (mean±SEM, Mann Whitney test, n=clusters, r=2). Scale bars, 50µm. **B.** Smad3 siRNA transfection suppresses TGFβ1 treated 1863 cluster invasion. Dot plot shows the quantification of invasion (mean±SEM, Mann Whitney test, n=clusters, r=2). Scale bars, 50 µm. Bar graph shows Smad3 expression (mean±SEM, n=2). **C.** Pooled and individual Smad3 siRNAs suppress 1863T vertical invasion (mean±SEM, unpaired Student’s t test, n=3). Scale bars, 50 µm.

**Figure 4—figure supplement 2.**
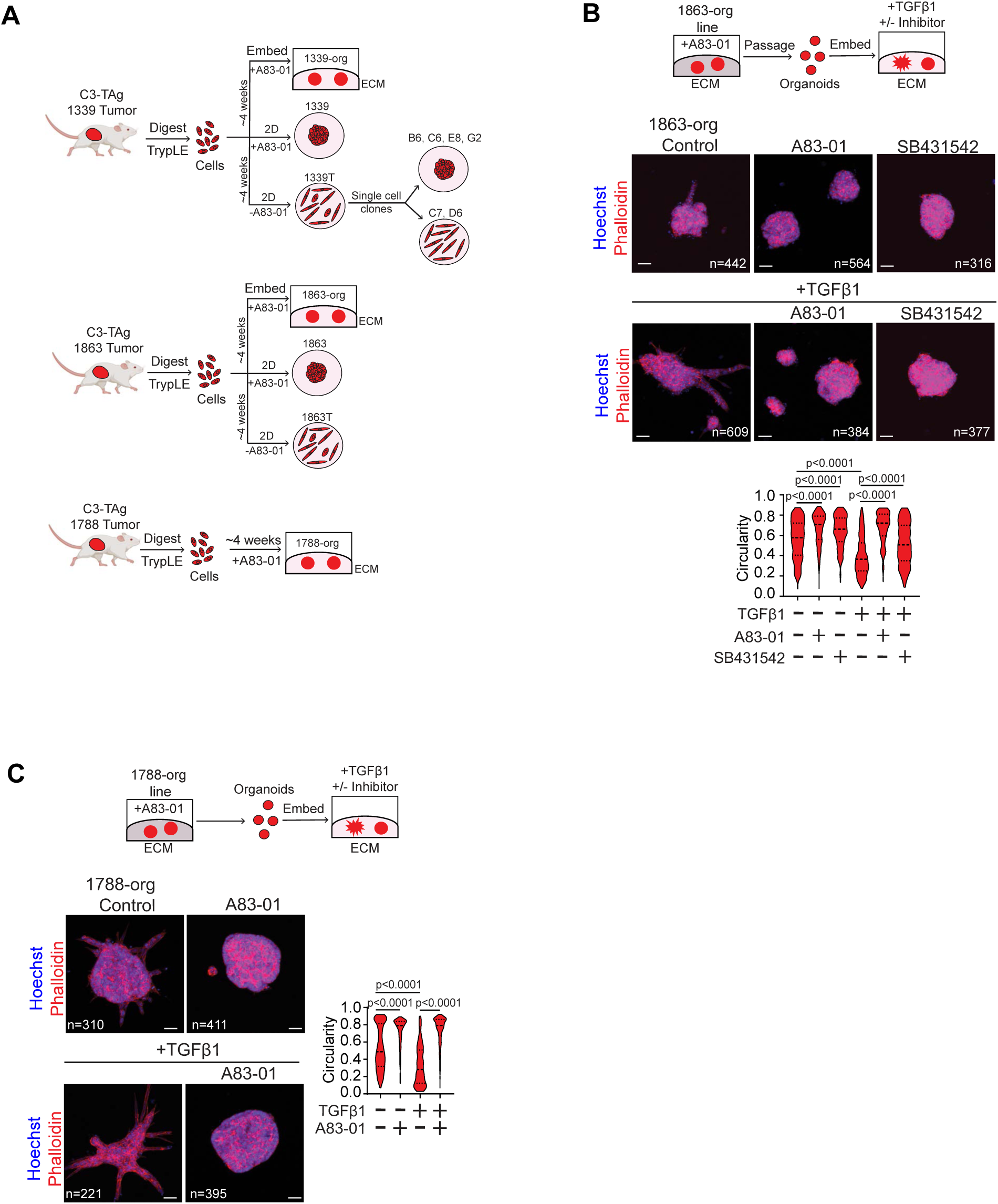
**A.** Graphics show the process of deriving 1339-org, 1863-org and 1788-org lines and their relationship to 2D cell lines derived from the same tumors. **B-C**. Tgfbr1 inhibition suppresses autocrine and exogenous TGFβ1 induced invasion of C3-TAg **(B)** 1863-org and **(C)** 1788-org lines. Organoids were treated with the Tgfbr1 inhibitors A83-01 (500 nM), SB431542 (1 µM) or exogenous TGFβ1 (2 ng/ml) as indicated. Violin plot shows the quantification of organoid circularity (Mann Whitney test, n=organoids, r=2). Scale bars, 50 µm.

**Figure 4—figure supplement 3.**
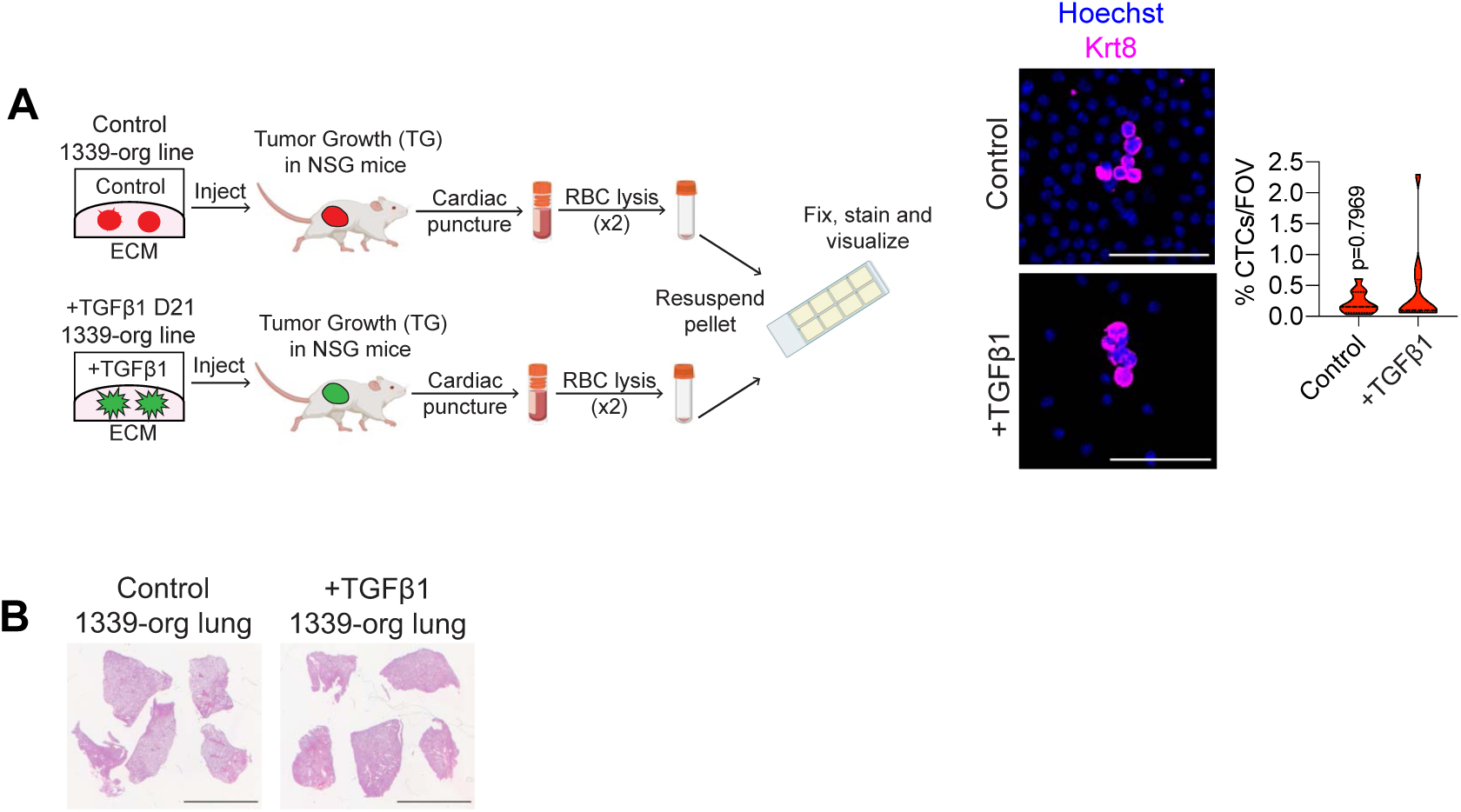
**A.** Graphic showing the process of CTC isolation from mice. Representative images show Krt8 expressing CTCs detected in mice with control 1339-org tumors and +TGFβ1 1339-org tumors. Violin plot shows percentage of total cells that were CTCs per field of view (Mann Whitney test, representative of blood collected from 7 mice per group, r=2). Scale bars, 100 µm; 50 µm (inset).

**Figure 4—Video 1. Live imaging of C3-TAg derived 1339-org line organoids in serum starved media (H2B:GFP, green, nuclei).** Fluorescent and bright field images were acquired for 18 h at 30 min intervals. The video corresponds with the baseline invasion of C3-TAg derived 1339-org line organoids as shown in **Fig. 3C, S3C**.

**Figure 4—Video 2. Live imaging of C3-TAg derived 1339-org line organoids in serum starved media + 500 nM A830-01 (Tgfbr1 inhibitor) (H2B:GFP, green, nuclei)**. Fluorescent and bright field images were acquired for 18 h at 30 min intervals. The video corresponds with suppression of 1339-org line organoid invasion by inhibiting autocrine TGFβR1 activity as shown in Fig. S3C.

**Figure 4—Video 3. Live imaging of C3-TAg derived 1339-org line organoids in presence of exogenous TGFβ1 (48 h) (H2B:GFP, green, nuclei).** Fluorescent and bright field images were acquired for 18 h at 30 min intervals. The video corresponds with TGFβ1 induced invasion in 1339-org line organoids as shown in **Fig. 3C, S3C**.

**Figure 4—Video 4. Live imaging of C3-TAg derived 1339-org line organoids in presence of persistent exogenous TGFβ1 (Day 16) (H2B:GFP, green, nuclei).** Fluorescent and bright field images were acquired for 18 h at 30 min intervals. The video corresponds with progressively increased TGFβ1 induced invasion in 1339-org line organoids as shown in Fig. 3C.

**Figure 5—figure supplement 1.**
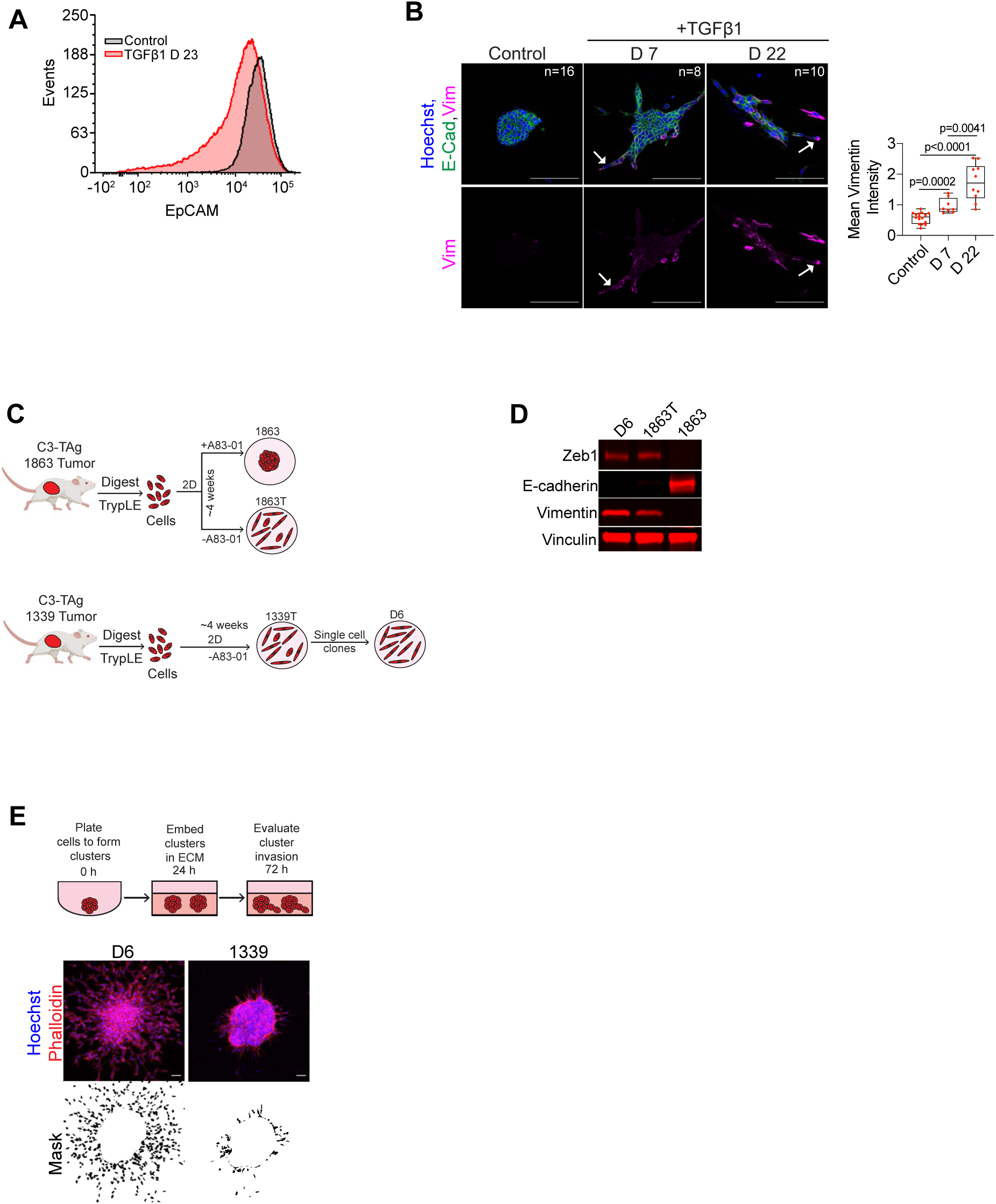
**A.** FACS showing that TGFβ1 treated 1339-org cells undergo a shift in the distribution of EpCAM expression (representative of 2 independent experiments). **B.** Images showing a progressive increase of Vimentin expression in TGFβ1 treated 1339-org cells. Arrows indicate representative Vimentin expressing cells leading collective invasion. Graph shows Vimentin expression in 1339-org organoids (Student’s t-test, n=organoids). Scale bars, 100 µm. **C.** Immunoblots showing the expression of Zeb1, E-cadherin and Vimentin in D6, 1863T and 1863 cell lines. Vinculin serves as a loading control (n=2). **D.** Immunoblots showing the expression of E-cadherin and Vimentin in non-invasive B6 and trailblazer D6 cell lines. Vinculin serves as a loading control. (n=2). **E. Representative images of D6 and 1339 clusters. Scale bars 50 µm.**

**Figure 6—figure supplement 1.**
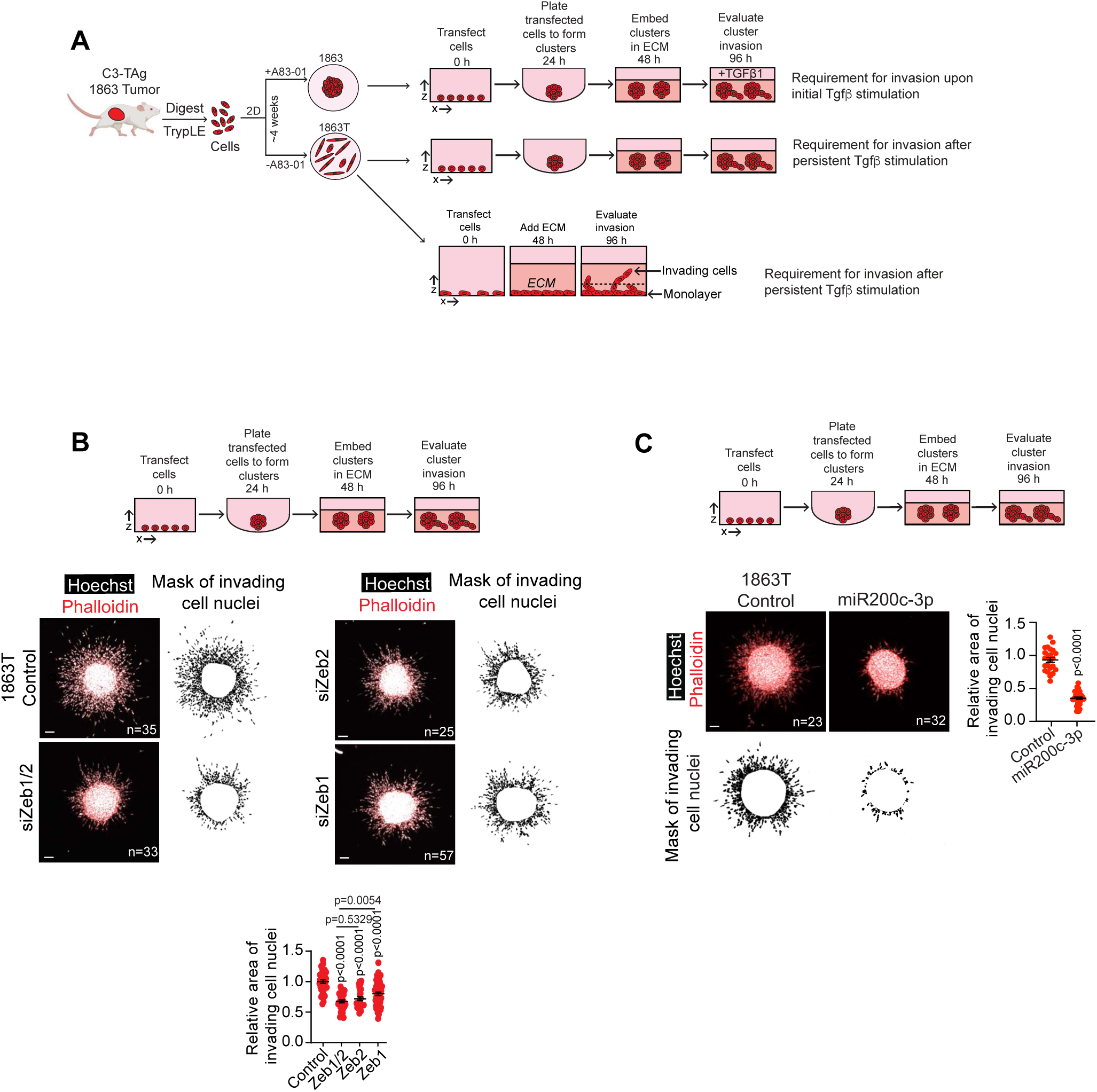
**A.** Zeb1 and Zeb2 siRNA transfection suppresses 1863T cluster invasion (mean±SEM, Mann Whitney test, n=clusters, r=3). Scale bars, 50 µm. **B.** Transfection with a miR200c-3p mimic decreases 1863T cluster invasion (mean±SEM, Mann Whitney test, n=clusters, r=2). Scale bars, 50 µm.

**Figure 6—figure supplement 2.**
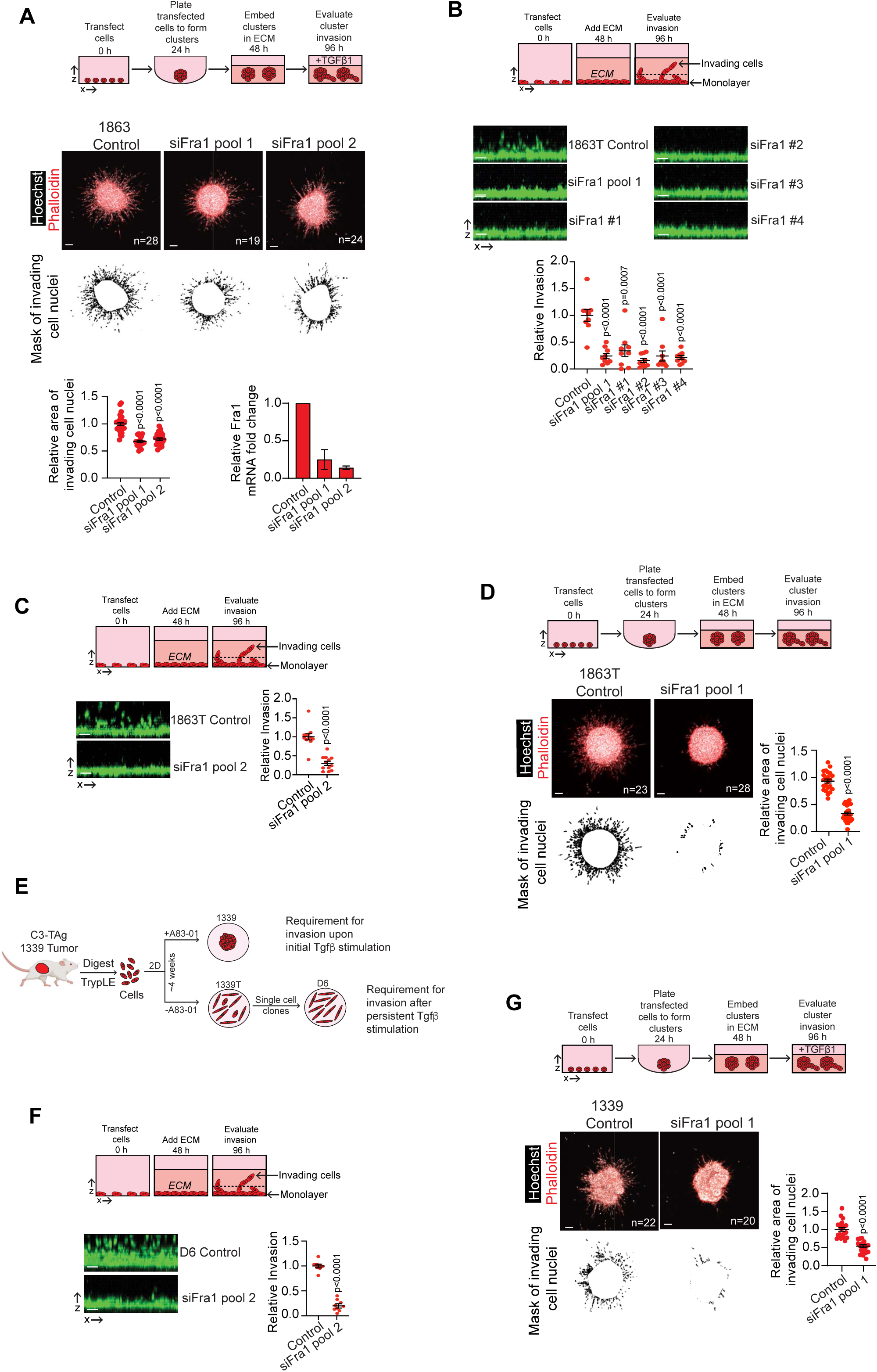
**A.** Fra1 depletion by a second distinct Fra1 siRNA pool (pool 2) decreases TGFβ1 treated 1863 cluster invasion. Dot plot shows the quantification of invasion (mean±SEM, Mann Whitney test, n=clusters, r=2). Bar graph shows the depletion of Fra1 expression by two different pools of Fra1 siRNA (mean±SEM, n=3 for pool 1 and n=2 for pool 2). Scale bars, 50 µm. **B.** 1863T vertical invasion is suppressed by each of the four individual sequences from Fra1 siRNA pool 1 (mean±SEM, Mann Whitney test, n= 9 wells, r=3) Scale bars, 50 µm. **C.** Fra1 siRNA pool 2 reduces the vertical invasion of 1863T cells (mean±SEM, Mann Whitney test, n=12 wells, r=4). Scale bars, 50 µm. **D.** Fra1 depletion suppresses the invasion of 1863T clusters (mean±SEM, Mann Whitney test, n=clusters, r=2). Scale bars, 50 µm. **E.** Fra1 depletion reduces the vertical invasion of D6 cells (mean±SEM, Mann Whitney test, n= 9 wells, r=3). Scale bars, 50 µm. **F.** Fra1 depletion reduces the invasion of TGFβ1 treated 1339 clusters (mean±SEM, Mann Whitney test, n=clusters, r=2). Scale bars, 50 µm.

**Figure 6—figure supplement 3.**
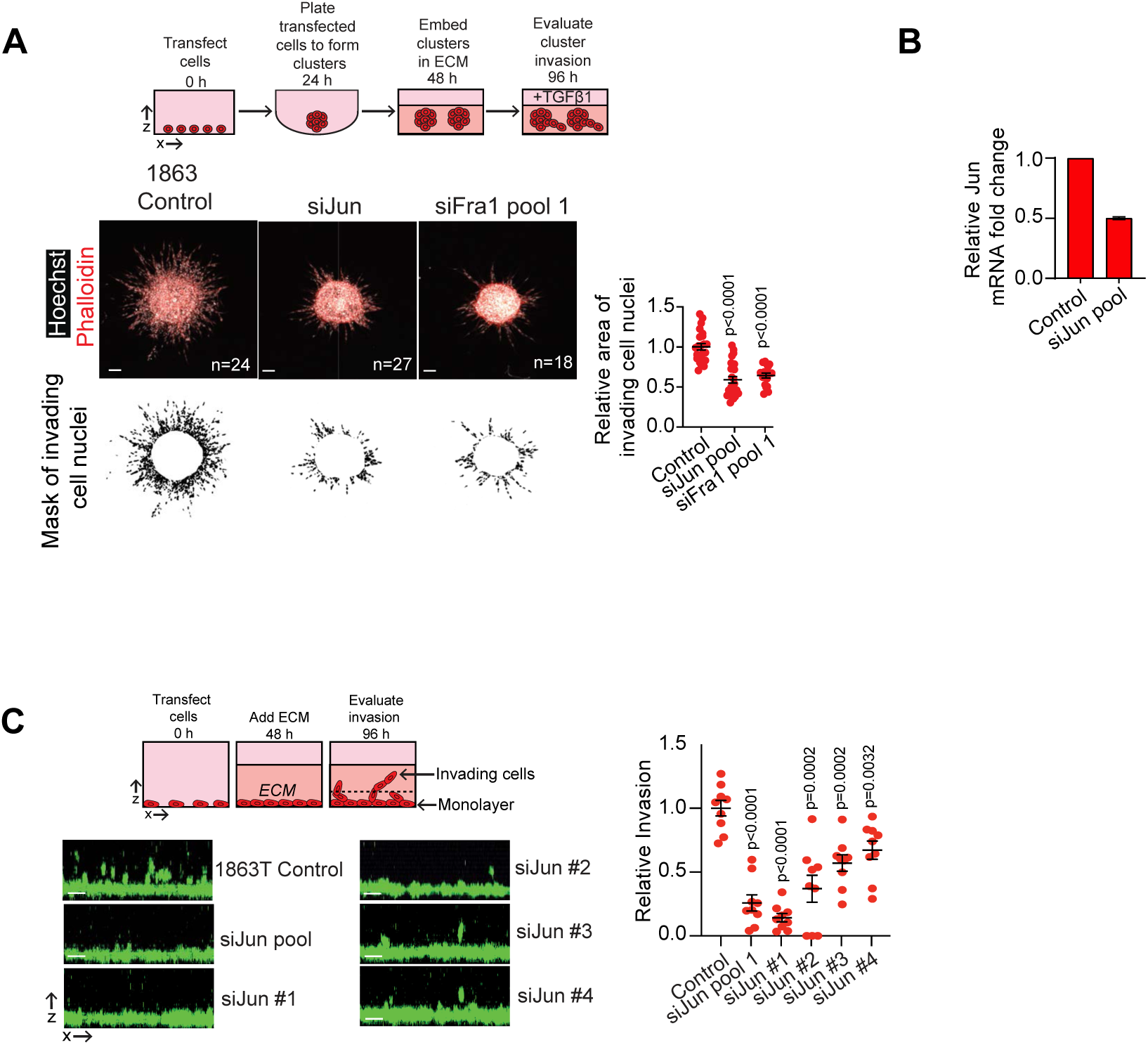
**A.** Jun depletion suppresses TGFβ1 induced 1863 cluster invasion (mean±SEM, Mann Whitney test, n=clusters, r=2) Scale bars, 50 µm. **B.** Depletion of Jun expression by Jun siRNA transfection (mean±SEM, n=2). **C.** Pooled and individual Jun siRNAs suppress 1863T vertical invasion (mean±SEM, unpaired Student’s t test, n=3). Scale bars, 50 µm.

**Figure 6—figure supplement 4.**
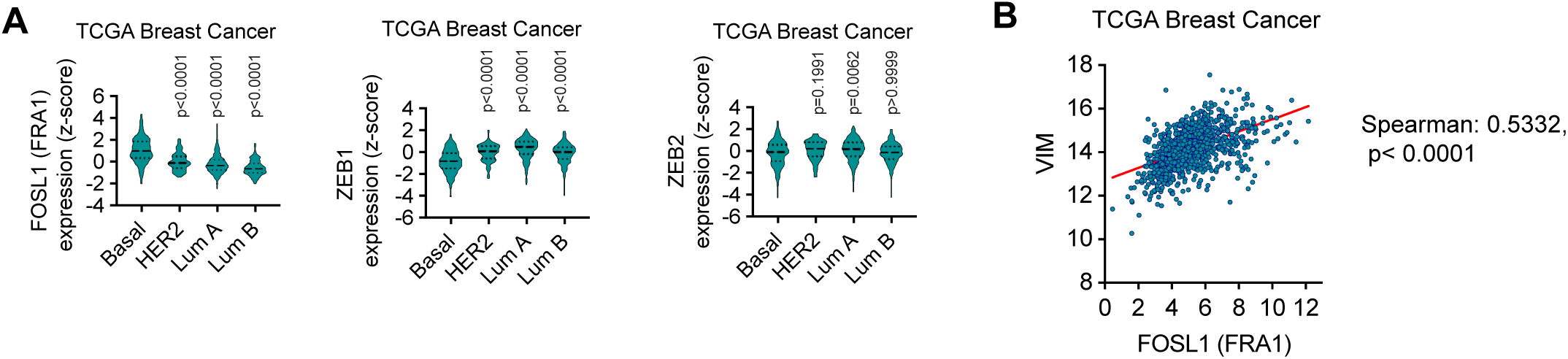
**A.** FOSL1 (FRA1) mRNA is more highly expressed in the basal-like breast cancer patient tumors compared to other breast cancer subtypes. By comparison, ZEB1 and ZEB2 mRNA is more highly expressed in other breast cancer cancer subtypes compared to basal-like breast cancer. Z-scores relative to all samples are shown (Kruskal Wallis test with Dunn’s multiple comparisons test). P-values are in comparison to basal-like tumors **B.** Correlation of VIM and FOSL1 (FRA1) expression in breast cancer patient tumors.

**Figure 7—figure supplement 1.**
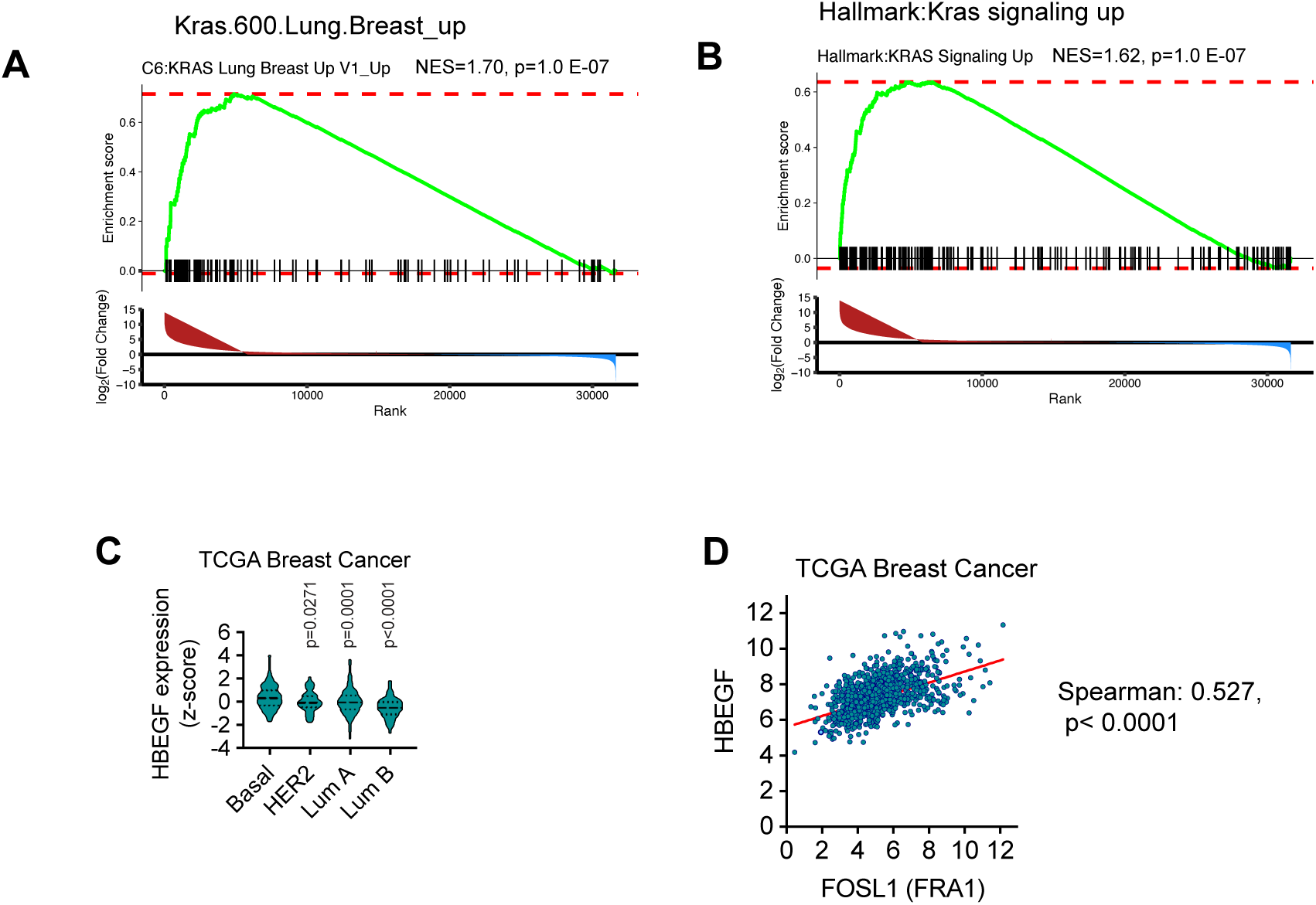
**A-B.** GSEA was performed on RNA-seq data from C3-TAg organoids and non-invasive C3-TAg clone spheroids to identify pathway enrichment in C3-TAg organoids. Plots show the GSEA associated with Kras signaling. **C**. HBEGF mRNA is more highly expressed in the basal-like breast cancer patient tumors compared to other breast cancer subtypes. Z-scores relative to all samples are shown (Kruskal Wallis test with Dunn’s multiple comparisons test). P-values are in comparison to the basal-like tumors. **D**. Correlation of FOSL1 (FRA1) and HBEGF expression in breast cancer patient tumors.

**Figure 7—figure supplement 2.**
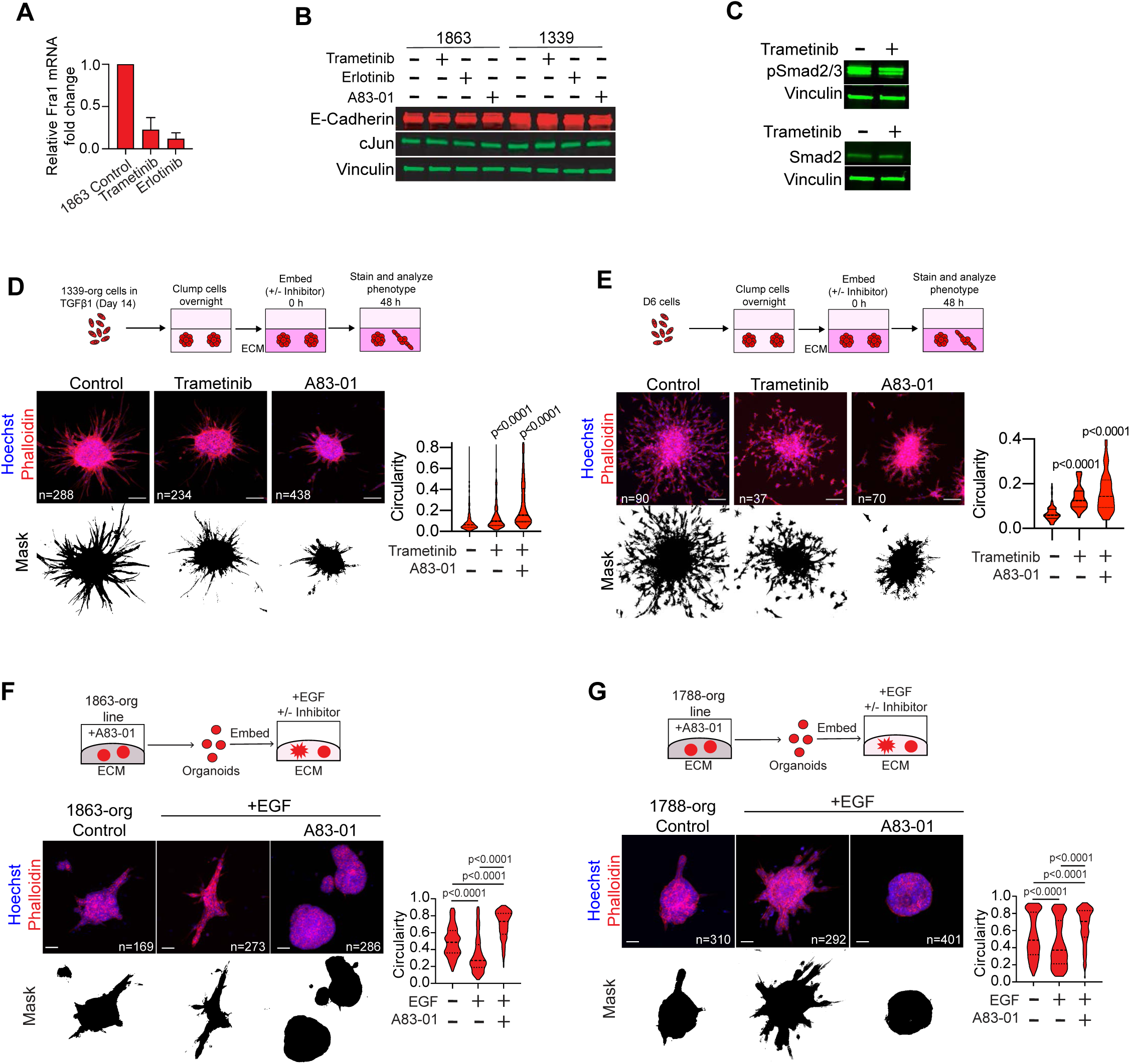
**A**. Trametinib (10 nM) and erlotinib (1 µM) reduce Fra1 expression in 1863 cells (mean±SEM, n=2). **B**. Immunoblot showing that Jun expression is not perturbed by treatment with trametinib (10 nM), erlotinib (1 µM) or A83-01 (500 nM) in 1863 and 1339 cells (n=2 for erlotinib, A8301; n=3 for trametinib). **C**. Immunoblot shows that trametinib (10 nM) does not reduce Zeb1 expression or Smad2 phosphorylation in C3-TAg cells (n=2). **D**. 1339-org organoids were persistently stimulated with exogenous TGFβ1 for 14 days to induce a trailblazer state (D14 TGFβ1 1339-org). The ability of trametinib and A83-01 to suppress then invasion of the D14 TGFβ1 1339-org cells was then determined. Violin plots show the quantification of invasion (Mann Whitney test, n=organoids, r=2). Scale bars, 100 µm. **E.** Trametinib and A83-01 suppress the invasion of D6 trailblazer cells (Mann Whitney test, n=spheroids, r=2). **F-G**. The Tgfbr1 inhibitor A83-01 suppresses the EGF induced invasion of **(F)** 1863-org and **(G)** 1788-org organoids. Violin plot shows the quantification of organoid circularity (Mann Whitney test, n=organoids, r=2). Scale bars, 50 µm.

**Figure 7—figure supplement 3.**
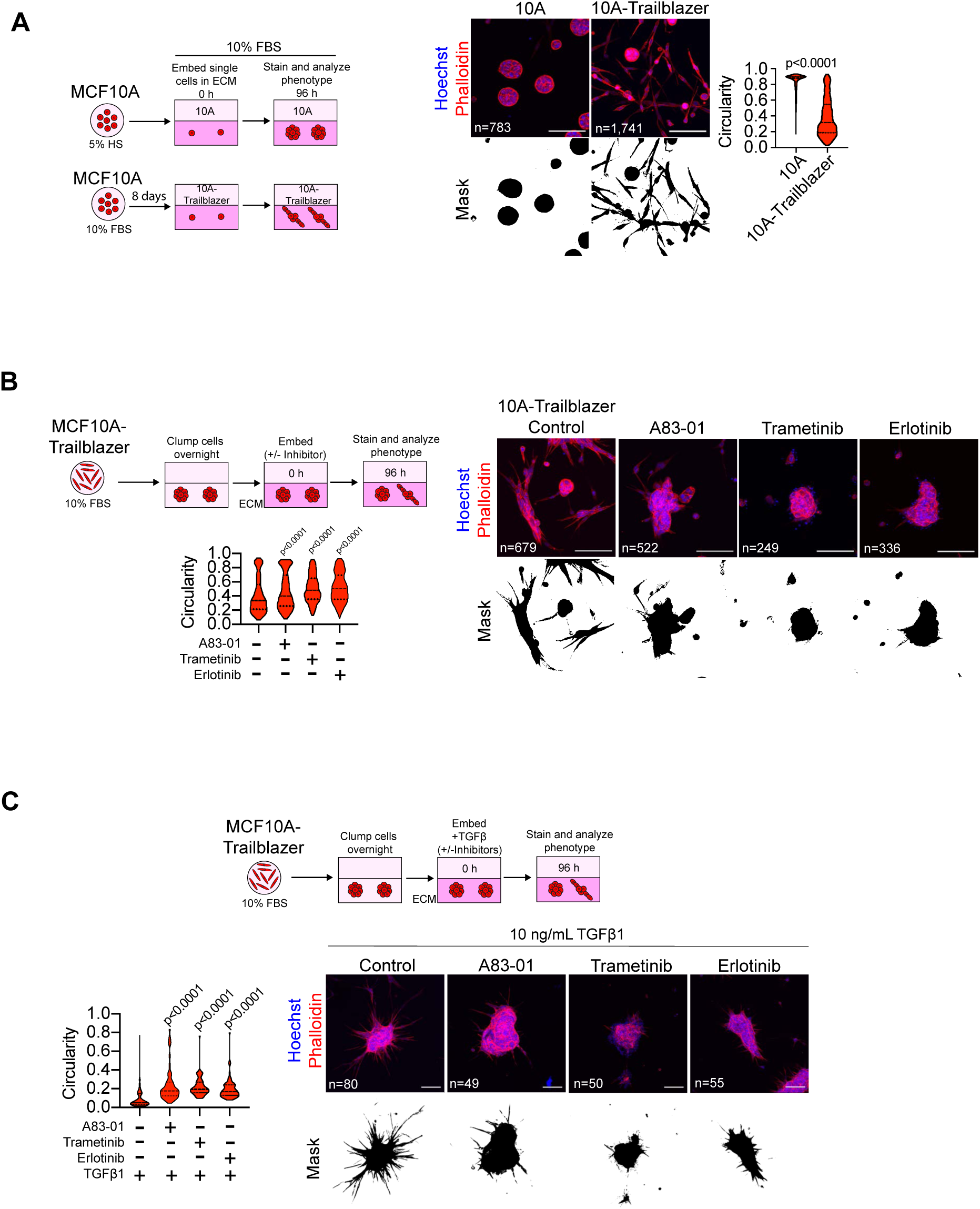
**A**. Graphic and data shows how a MCF10A-trailblazer cell line was established be treating MCF10A cells with 10% FBS. MCF10A cells were supplemented with 10% FBS for 8 days in monolayer culture. The invasive ability of control and 10% FBS treated MCF10A cells was then determined. The 10% FBS induced a trailblazer state in the MCF10A cells, which we term “MCF10A-Trailblazer”. Violin plot shows quantification of circularity. (Mann Whitney test, n=spheroids, r=2). Scale bars, 100 µm. **B.** A83-01, trametinib and erlotinib suppresses MCF10A-trailblazer invasion (Mann Whitney test, n=cspheroids, r=2) Scale bars, 100 µm. **C.** A83-01, trametinib and erlotinib suppress the invasion of MCF10A-trailblazer cells treated with exogenous TGFβ1 (Mann Whitney test, n=spheroids, r=2) Scale bars, 100 µm.

**Figure 7—figure supplement 4.**
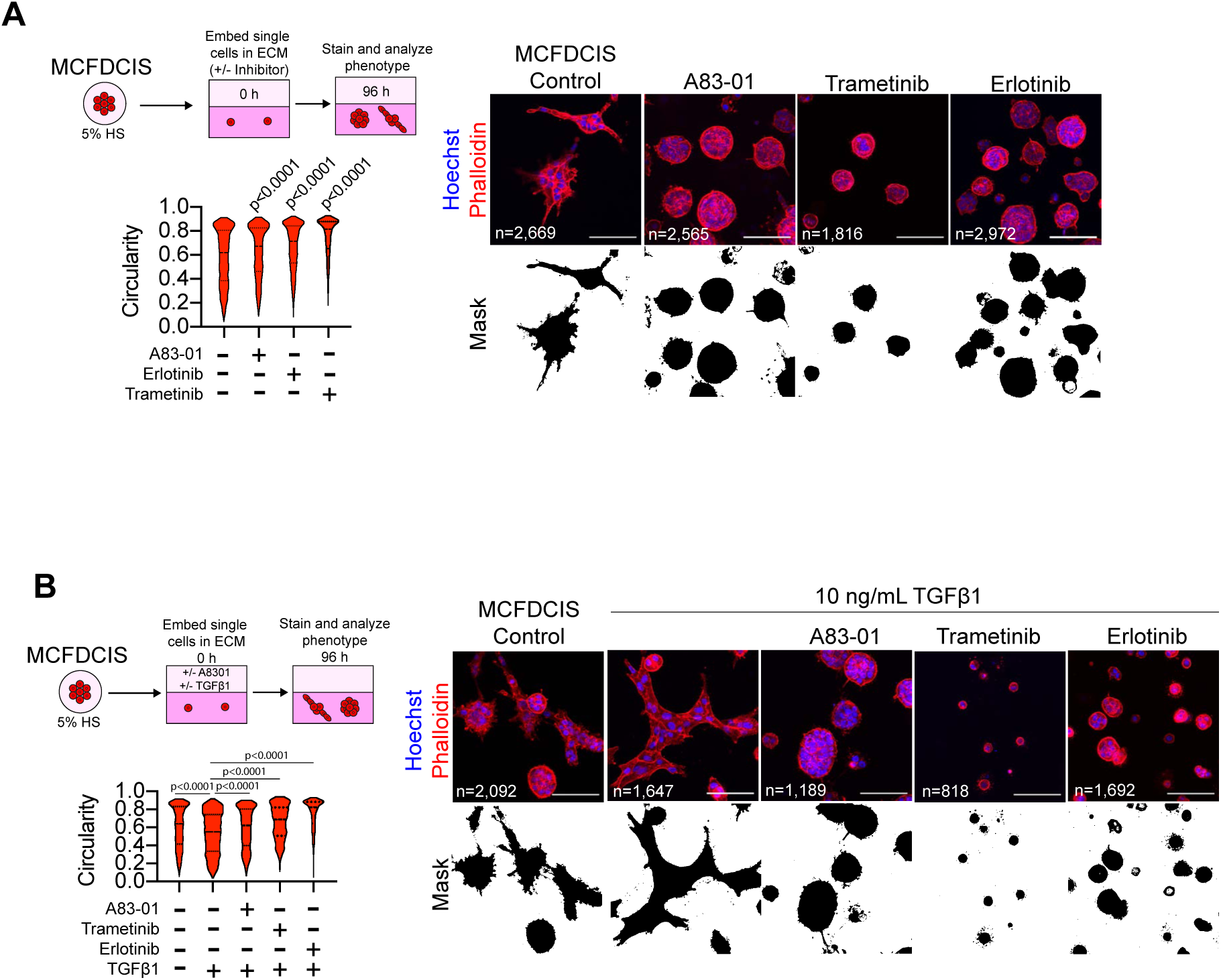
**A**. A83-01, trametinib and erlotinib suppress MCFDCIS spheroid invasion (Mann Whitney test, n=spheroids, r=2) Scale bars, 100 µm. **B**. A83-01, trametinib and erlotinib suppress the TGFβ1 induced invasion of MCFDCIS spheroids (Mann Whitney test, n=spheroids, r=2) Scale bars, 100 µm.

**Figure 8—figure supplement 1.**
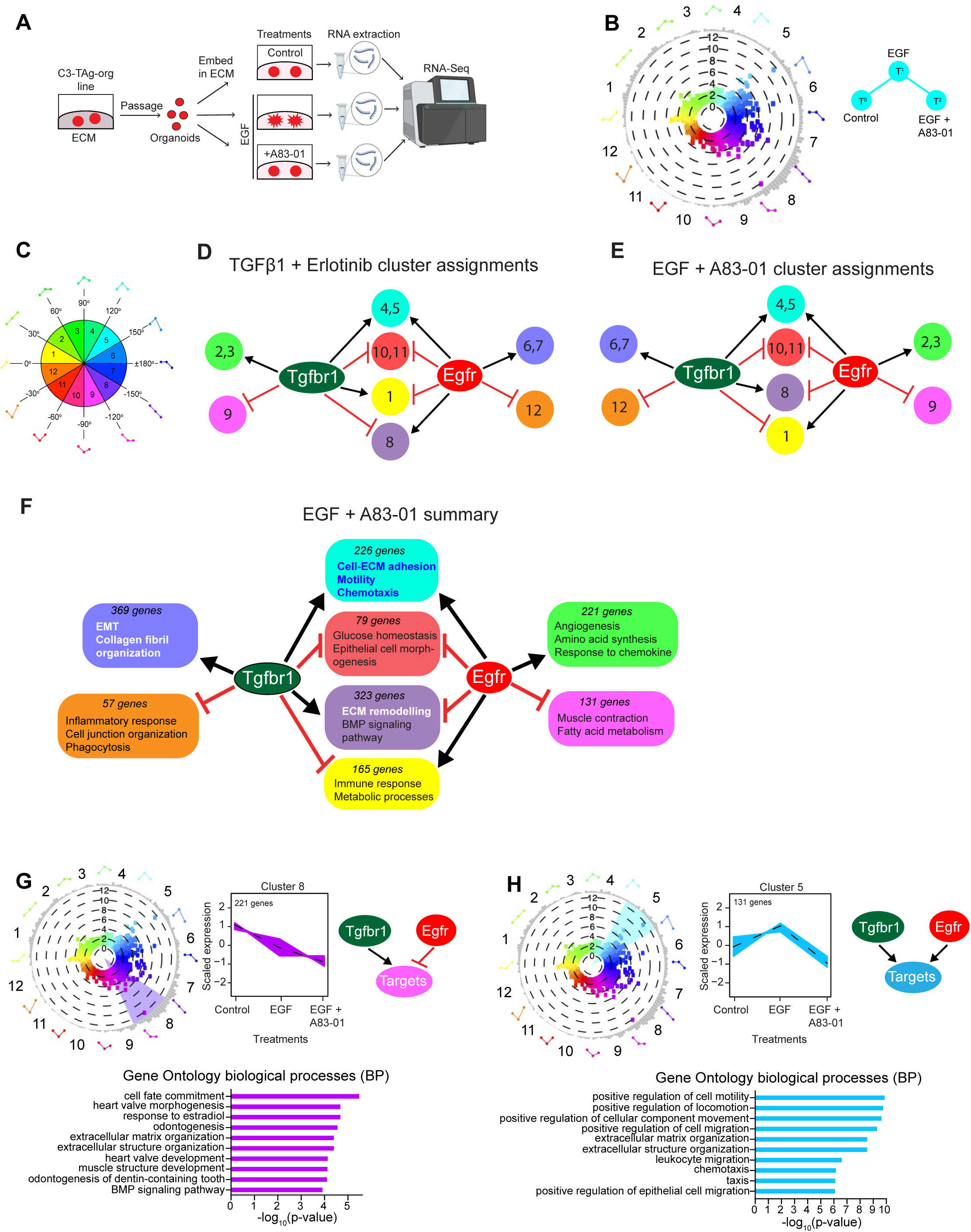
**A**. Overview of the approach to test the interactions between the Egfr and Tgfβ signaling pathways when cells were treated with EGF or EGF+A83-01. **B.** Polar plot organizing genes represented as individual dots based on their change in expression in response to treatment with EGF or co-treatment with EGF and A83-01. Genes that underwent ≥2 fold-change with a p< 0.05 after Benjamini-Hochberg correction in at least one condition relative to one of the other conditions are shown (3 total comparisons). The distance from the center of the plot represents the Δlog2 value of each gene (increased difference in expression, further from the center). The bar plots on the surface of the circle show the number of genes at the degree point in the plot. Genes are grouped into 12 color-coded groups along 30 degree increments over the 360 degree plot. As an example, genes in group 5 were increased in expression by EGF and this increase in expression was inhibited by A83-01. In contrast, genes in cluster 1 were increased in expression by EGF, and this induction is further enhanced by treatment with A83-01. **C.** Plot shows the cluster locations along the 30 degree increments in the polar plots shown in **Figure 8B and Figure 8—figure supplement 1B**. **D.** Clusters assigned to different summary groups for TGFβ1 and TGFβ1+erlotinib treated organoids shown in Figure 8B. **E.** Clusters assigned to different summary groups for EGF and EGF+A83-01 treated organoids shown in **Figure 8—figure supplement 1B**. **F.** Summary of the number of genes and represented biological processes associated with different modes of interaction between the Tgfbr1 and Egfr pathways in cells treated with EGF and EGF+A83-01. Also see **Supplementary file 5** for the assignment of polar plot clusters and detailed biological processes used to define the summary integration groups. **G.** Egfr restricta the expression of a subset of genes induced by autocrine Tgfβ. Line plots show the expression of genes in Cluster 1. Dashed line indicates the average scaled expression of all Cluster 1 genes. Graph shows the top 10 biological processes (GO-BP) associated with Cluster 1 genes (Fisher’s exact test). **H.** Tgfbr1 activity was necessary for the expression of a subset of genes induced by extrinsic EGF. Line plots show the expression of genes in Cluster 5. Dashed line indicates the average scaled expression of all Cluster 5 genes. Graph shows the top 10 biological processes (GO-BP) associated with Cluster 5 genes (Fisher’s exact test).

**Supplementary file 1.** Description of GEM models, organoid/cell line models and reagents used in the study.

**Supplementary file 2.** RNA-Seq data (log2(CPM) genes x samples) from comparison between non-invasive clones and C3-TAg tumor organoids. Pathway enrichment analysis from comparison between non-invasive clones and C3-TAg tumor organoids

**Supplementary file 3.** Gene clusters and enriched biological processes from comparison between control and TGFβ1 treated C3-TAg 1339-org line organoids

**Supplementary file 4.** Gene clusters and enriched biological processes from comparison between control (T°), TGFβ1 (T^1^) and TGFβ1 + Erlotinib (T^2^) treated C3-TAg 1339-org line organoids.

**Supplementary file 5.** Gene clusters and enriched biological processes from comparison between control (T°), EGF (T^1^) and EGF + A8301 (T^2^) treated C3-TAg 1339-org line organoids.

## Source Data Legends

Full annotated and raw immunoblots corresponding to **Figure 1—figure supplement 1F, Figure 2—figure supplement 1B, Figure 5—figure supplement 1D, Figure 7C and Figure 7— figure supplement 2B-C.**

## REFERENCES

1. Marusyk A, Polyak K. Tumor heterogeneity: causes and consequences. Biochim Biophys Acta. 2010;1805(1):105–17. Epub 2009/11/26. doi: 10.1016/j.bbcan.2009.11.002. PubMed PMID: 19931353; PMCID: PMC2814927.

2. Beca F, Polyak K. Intratumor Heterogeneity in Breast Cancer. Adv Exp Med Biol. 2016;882:169–89. Epub 2016/03/19. doi: 10.1007/978-3-319-22909-6_7. PubMed PMID: 26987535.

3. McGranahan N, Swanton C. Biological and therapeutic impact of intratumor heterogeneity in cancer evolution. Cancer Cell. 2015;27(1):15–26. doi: 10.1016/j.ccell.2014.12.001. PubMed PMID: 25584892.

4. Marusyk A, Janiszewska M, Polyak K. Intratumor Heterogeneity: The Rosetta Stone of Therapy Resistance. Cancer Cell. 2020;37(4):471–84. Epub 2020/04/15. doi: 10.1016/j.ccell.2020.03.007. PubMed PMID: 32289271; PMCID: PMC7181408.

5. Ganesh K, Massagué J. Targeting metastatic cancer. Nat Med. 2021;27(1):34–44. Epub 2021/01/15. doi: 10.1038/s41591-020-01195-4. PubMed PMID: 33442008; PMCID: PMC7895475.

6. Calbo J, van Montfort E, Proost N, van Drunen E, Beverloo HB, Meuwissen R, Berns A. A functional role for tumor cell heterogeneity in a mouse model of small cell lung cancer. Cancer Cell. 2011;19(2):244–56. Epub 2011/02/15. doi: 10.1016/j.ccr.2010.12.021. PubMed PMID: 21316603.

7. Dongre A, Rashidian M, Reinhardt F, Bagnato A, Keckesova Z, Ploegh HL, Weinberg RA. Epithelial-to-Mesenchymal Transition Contributes to Immunosuppression in Breast Carcinomas. Cancer Res. 2017;77(15):3982–9. Epub 2017/04/22. doi: 10.1158/0008-5472.Can-16-3292. PubMed PMID: 28428275; PMCID: PMC5541771.

8. Celià-Terrassa T, Meca-Cortés O, Mateo F, Martínez de Paz A, Rubio N, Arnal-Estapé A, Ell BJ, Bermudo R, Díaz A, Guerra-Rebollo M, Lozano JJ, Estarás C, Ulloa C, Álvarez-Simón D, Milà J, Vilella R, Paciucci R, Martínez-Balbás M, de Herreros AG, Gomis RR, Kang Y, Blanco J, Fernández PL, Thomson TM. Epithelial-mesenchymal transition can suppress major attributes of human epithelial tumor-initiating cells. J Clin Invest. 2012;122(5):1849–68. Epub 2012/04/17. doi: 10.1172/jci59218. PubMed PMID: 22505459; PMCID: PMC3366719.

9. Marusyk A, Tabassum DP, Altrock PM, Almendro V, Michor F, Polyak K. Non-cell-autonomous driving of tumour growth supports sub-clonal heterogeneity. Nature. 2014;514(7520):54–8. doi: 10.1038/nature13556. PubMed PMID: 25079331; PMCID: 4184961.

10. Zhang M, Tsimelzon A, Chang CH, Fan C, Wolff A, Perou CM, Hilsenbeck SG, Rosen JM. Intratumoral heterogeneity in a Trp53-null mouse model of human breast cancer. Cancer Discov. 2015;5(5):520–33. doi: 10.1158/2159-8290.CD-14-1101. PubMed PMID: 25735774; PMCID: 4420701.

11. Neelakantan D, Zhou H, Oliphant MUJ, Zhang X, Simon LM, Henke DM, Shaw CA, Wu MF, Hilsenbeck SG, White LD, Lewis MT, Ford HL. EMT cells increase breast cancer metastasis via paracrine GLI activation in neighbouring tumour cells. Nat Commun. 2017;8:15773. doi: 10.1038/ncomms15773. PubMed PMID: 28604738; PMCID: 5472791.

12. Dongre A, Rashidian M, Eaton EN, Reinhardt F, Thiru P, Zagorulya M, Nepal S, Banaz T, Martner A, Spranger S, Weinberg RA. Direct and Indirect Regulators of Epithelial-Mesenchymal Transition-Mediated Immunosuppression in Breast Carcinomas. Cancer Discov. 2021;11(5):1286–305. Epub 2020/12/18. doi: 10.1158/2159-8290.Cd-20-0603. PubMed PMID: 33328216; PMCID: PMC8432413.

13. Sharif GM, Campbell MJ, Nasir A, Sengupta S, Graham GT, Kushner MH, Kietzman WB, Schmidt MO, Pearson GW, Loudig O, Fineberg S, Wellstein A, Riegel AT. An AIB1 Isoform Alters Enhancer Access and Enables Progression of Early-Stage Triple-Negative Breast Cancer. Cancer Res. 2021;81(16):4230–41. Epub 2021/06/18. doi: 10.1158/0008-5472.CAN-20-3625. PubMed PMID: 34135000; PMCID: PMC8373795.

14. Zhou H, Neelakantan D, Ford HL. Clonal cooperativity in heterogenous cancers. Semin Cell Dev Biol. 2017;64:79–89. doi: 10.1016/j.semcdb.2016.08.028. PubMed PMID: 27582427; PMCID: 5330947.

15. Friedl P, Gilmour D. Collective cell migration in morphogenesis, regeneration and cancer. Nat Rev Mol Cell Biol. 2009;10(7):445–57. Epub 2009/06/24. doi: 10.1038/nrm2720. PubMed PMID: 19546857.

16. Nguyen-Ngoc KV, Cheung KJ, Brenot A, Shamir ER, Gray RS, Hines WC, Yaswen P, Werb Z, Ewald AJ. ECM microenvironment regulates collective migration and local dissemination in normal and malignant mammary epithelium. Proc Natl Acad Sci U S A. 2012;109(39):E2595–604. doi: 10.1073/pnas.1212834109. PubMed PMID: 22923691; PMCID: 3465416.

17. Bronsert P, Enderle-Ammour K, Bader M, Timme S, Kuehs M, Csanadi A, Kayser G, Kohler I, Bausch D, Hoeppner J, Hopt UT, Keck T, Stickeler E, Passlick B, Schilling O, Reiss CP, Vashist Y, Brabletz T, Berger J, Lotz J, Olesch J, Werner M, Wellner UF. Cancer cell invasion and EMT marker expression: a three-dimensional study of the human cancer-host interface. J Pathol. 2014;234(3):410–22. doi: 10.1002/path.4416. PubMed PMID: 25081610.

18. Cheung KJ, Padmanaban V, Silvestri V, Schipper K, Cohen JD, Fairchild AN, Gorin MA, Verdone JE, Pienta KJ, Bader JS, Ewald AJ. Polyclonal breast cancer metastases arise from collective dissemination of keratin 14-expressing tumor cell clusters. Proc Natl Acad Sci U S A. 2016;113(7):E854–63. doi: 10.1073/pnas.1508541113. PubMed PMID: 26831077; PMCID: 4763783.

19. Pearson GW. Control of Invasion by Epithelial-to-Mesenchymal Transition Programs during Metastasis. J Clin Med. 2019;8(5). doi: 10.3390/jcm8050646. PubMed PMID: 31083398; PMCID: 6572027.

20. Cheung KJ, Gabrielson E, Werb Z, Ewald AJ. Collective invasion in breast cancer requires a conserved basal epithelial program. Cell. 2013;155(7):1639–51. doi: 10.1016/j.cell.2013.11.029. PubMed PMID: 24332913; PMCID: 3941206.

21. Westcott JM, Prechtl AM, Maine EA, Dang TT, Esparza MA, Sun H, Zhou Y, Xie Y, Pearson GW. An epigenetically distinct breast cancer cell subpopulation promotes collective invasion. J Clin Invest. 2015;125(5):1927–43. doi: 10.1172/JCI77767. PubMed PMID: 25844900.

22. Kim YH, Choi YW, Lee J, Soh EY, Kim JH, Park TJ. Senescent tumor cells lead the collective invasion in thyroid cancer. Nat Commun. 2017;8:15208. doi: 10.1038/ncomms15208. PubMed PMID: 28489070; PMCID: 5436223.

23. Summerbell ER, Mouw JK, Bell JSK, Knippler CM, Pedro B, Arnst JL, Khatib TO, Commander R, Barwick BG, Konen J, Dwivedi B, Seby S, Kowalski J, Vertino PM, Marcus AI. Epigenetically heterogeneous tumor cells direct collective invasion through filopodia-driven fibronectin micropatterning. Sci Adv. 2020;6(30):eaaz6197. Epub 2020/08/25. doi: 10.1126/sciadv.aaz6197. PubMed PMID: 32832657; PMCID: PMC7439406.

24. Konen J, Summerbell E, Dwivedi B, Galior K, Hou Y, Rusnak L, Chen A, Saltz J, Zhou W, Boise LH, Vertino P, Cooper L, Salaita K, Kowalski J, Marcus AI. Image-guided genomics of phenotypically heterogeneous populations reveals vascular signalling during symbiotic collective cancer invasion. Nat Commun. 2017;8:15078. doi: 10.1038/ncomms15078. PubMed PMID: 28497793; PMCID: 5437311.

25. Westcott JM, Camacho S, Nasir A, Huysman ME, Rahhal R, Dang TT, Riegel AT, Brekken RA, Pearson GW. ΔNp63-Regulated Epithelial-to-Mesenchymal Transition State Heterogeneity Confers a Leader–Follower Relationship That Drives Collective Invasion. Cancer Research. 2020;80(18):3933–44. doi: 10.1158/0008-5472.Can-20-0014.

26. Tabassum DP, Polyak K. Tumorigenesis: it takes a village. Nat Rev Cancer. 2015;15(8):473–83. doi: 10.1038/nrc3971. PubMed PMID: 26156638.

27. Celia-Terrassa T, Kang Y. Distinctive properties of metastasis-initiating cells. Genes Dev. 2016;30(8):892–908. doi: 10.1101/gad.277681.116. PubMed PMID: 27083997; PMCID: 4840296.

28. Maine EA, Westcott JM, Prechtl AM, Dang TT, Whitehurst AW, Pearson GW. The cancer-testis antigens SPANX-A/C/D and CTAG2 promote breast cancer invasion. Oncotarget. 2016;7(12):14708–26. doi: 10.18632/oncotarget.7408. PubMed PMID: 26895102.

29. Padmanaban V, Tsehay Y, Cheung KJ, Ewald AJ, Bader JS. Between-tumor and within-tumor heterogeneity in invasive potential. PLoS Comput Biol. 2020;16(1):e1007464. Epub 2020/01/22. doi: 10.1371/journal.pcbi.1007464. PubMed PMID: 31961880; PMCID: PMC6994152.

30. Winkler J, Abisoye-Ogunniyan A, Metcalf KJ, Werb Z. Concepts of extracellular matrix remodelling in tumour progression and metastasis. Nat Commun. 2020;11(1):5120. Epub 2020/10/11. doi: 10.1038/s41467-020-18794-x. PubMed PMID: 33037194; PMCID: PMC7547708.

31. Dang TT, Esparza MA, Maine EA, Westcott JM, Pearson GW. DeltaNp63alpha Promotes Breast Cancer Cell Motility through the Selective Activation of Components of the Epithelial-to-Mesenchymal Transition Program. Cancer Res. 2015;75(18):3925–35. doi: 10.1158/0008-5472.CAN-14-3363. PubMed PMID: 26292362; PMCID: 4573836.

32. Li CI, Uribe DJ, Daling JR. Clinical characteristics of different histologic types of breast cancer. British Journal of Cancer. 2005;93(9):1046–52. doi: 10.1038/sj.bjc.6602787.

33. Yersal O, Barutca S. Biological subtypes of breast cancer: Prognostic and therapeutic implications. World J Clin Oncol. 2014;5(3):412–24. Epub 2014/08/13. doi: 10.5306/wjco.v5.i3.412. PubMed PMID: 25114856; PMCID: PMC4127612.

34. Green JE, Shibata MA, Yoshidome K, Liu ML, Jorcyk C, Anver MR, Wigginton J, Wiltrout R, Shibata E, Kaczmarczyk S, Wang W, Liu ZY, Calvo A, Couldrey C. The C3(1)/SV40 T-antigen transgenic mouse model of mammary cancer: ductal epithelial cell targeting with multi-stage progression to carcinoma. Oncogene. 2000;19(8):1020–7. doi: 10.1038/sj.onc.1203280. PubMed PMID: 10713685.

35. Herschkowitz JI, Simin K, Weigman VJ, Mikaelian I, Usary J, Hu Z, Rasmussen KE, Jones LP, Assefnia S, Chandrasekharan S, Backlund MG, Yin Y, Khramtsov AI, Bastein R, Quackenbush J, Glazer RI, Brown PH, Green JE, Kopelovich L, Furth PA, Palazzo JP, Olopade OI, Bernard PS, Churchill GA, Van Dyke T, Perou CM. Identification of conserved gene expression features between murine mammary carcinoma models and human breast tumors. Genome Biol. 2007;8(5):R76. Epub 2007/05/12. doi: 10.1186/gb-2007-8-5-r76. PubMed PMID: 17493263; PMCID: PMC1929138.

36. Hwang PY, Brenot A, King AC, Longmore GD, George SC. Randomly Distributed K14(+) Breast Tumor Cells Polarize to the Leading Edge and Guide Collective Migration in Response to Chemical and Mechanical Environmental Cues. Cancer Res. 2019;79(8):1899–912. Epub 2019/03/14. doi: 10.1158/0008-5472.Can-18-2828. PubMed PMID: 30862718; PMCID: PMC6467777.

37. Bergholtz H, Lien TG, Swanson DM, Frigessi A, Bathen TF, Borgen E, Børresen-Dale AL, Engebråten O, Garred Ø, Geisler J, Geitvik GA, Hartmann-Johnsen OJ, Hofvind S, Kristensen VN, Langerød A, Lingjærde OC, Mælandsmo GM, Naume B, Russnes H, Sauer T, Schlichting E, Skjerven HK, Daidone MG, Tost J, Wärnberg F, Sørlie T, Oslo Breast Cancer Research C. Contrasting DCIS and invasive breast cancer by subtype suggests basal-like DCIS as distinct lesions. npj Breast Cancer. 2020;6(1):26. doi: 10.1038/s41523-020-0167-x.

38. Christin JR, Wang C, Chung CY, Liu Y, Dravis C, Tang W, Oktay MH, Wahl GM, Guo W. Stem Cell Determinant SOX9 Promotes Lineage Plasticity and Progression in Basal-like Breast Cancer. Cell Rep. 2020;31(10):107742. Epub 2020/06/11. doi: 10.1016/j.celrep.2020.107742. PubMed PMID: 32521267; PMCID: PMC7658810.

39. DeRose YS, Wang G, Lin YC, Bernard PS, Buys SS, Ebbert MT, Factor R, Matsen C, Milash BA, Nelson E, Neumayer L, Randall RL, Stijleman IJ, Welm BE, Welm AL. Tumor grafts derived from women with breast cancer authentically reflect tumor pathology, growth, metastasis and disease outcomes. Nat Med. 2011;17(11):1514–20. Epub 2011/10/25. doi: 10.1038/nm.2454. PubMed PMID: 22019887; PMCID: PMC3553601.

40. Pastushenko I, Brisebarre A, Sifrim A, Fioramonti M, Revenco T, Boumahdi S, Van Keymeulen A, Brown D, Moers V, Lemaire S, De Clercq S, Minguijon E, Balsat C, Sokolow Y, Dubois C, De Cock F, Scozzaro S, Sopena F, Lanas A, D’Haene N, Salmon I, Marine JC, Voet T, Sotiropoulou PA, Blanpain C. Identification of the tumour transition states occurring during EMT. Nature. 2018;556(7702):463–8. doi: 10.1038/s41586-018-0040-3. PubMed PMID: 29670281.

41. Altschuler SJ, Wu LF. Cellular heterogeneity: do differences make a difference? Cell. 2010;141(4):559–63. Epub 2010/05/19. doi: 10.1016/j.cell.2010.04.033. PubMed PMID: 20478246; PMCID: PMC2918286.

42. Eferl R, Wagner EF. AP-1: a double-edged sword in tumorigenesis. Nature Reviews Cancer. 2003;3(11):859–68. doi: 10.1038/nrc1209.

43. Bakiri L, Macho-Maschler S, Custic I, Niemiec J, Guío-Carrión A, Hasenfuss SC, Eger A, Müller M, Beug H, Wagner EF. Fra-1/AP-1 induces EMT in mammary epithelial cells by modulating Zeb1/2 and TGFβ expression. Cell Death Differ. 2015;22(2):336–50. Epub 2014/10/10. doi: 10.1038/cdd.2014.157. PubMed PMID: 25301070.

44. Liu ML, Von Lintig FC, Liyanage M, Shibata MA, Jorcyk CL, Ried T, Boss GR, Green JE. Amplification of Ki-ras and elevation of MAP kinase activity during mammary tumor progression in C3(1)/SV40 Tag transgenic mice. Oncogene. 1998;17(18):2403–11. Epub 1998/11/12. doi: 10.1038/sj.onc.1202456. PubMed PMID: 9811472.

45. Albeck John G, Mills Gordon B, Brugge Joan S. Frequency-Modulated Pulses of ERK Activity Transmit Quantitative Proliferation Signals. Molecular Cell. 2013;49(2):249–61. doi: https://doi.org/10.1016/j.molcel.2012.11.002.

46. Neve RM, Chin K, Fridlyand J, Yeh J, Baehner FL, Fevr T, Clark L, Bayani N, Coppe JP, Tong F, Speed T, Spellman PT, DeVries S, Lapuk A, Wang NJ, Kuo WL, Stilwell JL, Pinkel D, Albertson DG, Waldman FM, McCormick F, Dickson RB, Johnson MD, Lippman M, Ethier S, Gazdar A, Gray JW. A collection of breast cancer cell lines for the study of functionally distinct cancer subtypes. Cancer Cell. 2006;10(6):515–27. Epub 2006/12/13. doi: 10.1016/j.ccr.2006.10.008. PubMed PMID: 17157791; PMCID: PMC2730521.

47. Siersbæk R, Madsen JGS, Javierre BM, Nielsen R, Bagge EK, Cairns J, Wingett SW, Traynor S, Spivakov M, Fraser P, Mandrup S. Dynamic Rewiring of Promoter-Anchored Chromatin Loops during Adipocyte Differentiation. Molecular Cell. 2017;66(3):420–35.e5. doi: https://doi.org/10.1016/j.molcel.2017.04.010.

48. Oft M, Peli J, Rudaz C, Schwarz H, Beug H, Reichmann E. TGF-beta1 and Ha-Ras collaborate in modulating the phenotypic plasticity and invasiveness of epithelial tumor cells. Genes Dev. 1996;10(19):2462–77. Epub 1996/10/01. doi: 10.1101/gad.10.19.2462. PubMed PMID: 8843198.

49. Derynck R, Muthusamy BP, Saeteurn KY. Signaling pathway cooperation in TGF-β-induced epithelial–mesenchymal transition. Current Opinion in Cell Biology. 2014;31:56–66. doi: https://doi.org/10.1016/j.ceb.2014.09.001.

50. Parvani JG, Galliher-Beckley AJ, Schiemann BJ, Schiemann WP. Targeted inactivation of β1 integrin induces β3 integrin switching, which drives breast cancer metastasis by TGF-β. Mol Biol Cell. 2013;24(21):3449–59. Epub 2013/09/06. doi: 10.1091/mbc.E12-10-0776. PubMed PMID: 24006485; PMCID: PMC3814150.

51. Blee AM, Huang H. ERG-Mediated Cell Invasion: A Link between Development and Tumorigenesis. Medical Epigenetics. 2015;3(2-3):19–29. doi: 10.1159/000440978.

52. Huang P, Chen A, He W, Li Z, Zhang G, Liu Z, Liu G, Liu X, He S, Xiao G, Huang F, Stenvang J, Brünner N, Hong A, Wang J. BMP-2 induces EMT and breast cancer stemness through Rb and CD44. Cell Death Discovery. 2017;3(1):17039. doi: 10.1038/cddiscovery.2017.39.

53. González-González L, Alonso J. Periostin: A Matricellular Protein With Multiple Functions in Cancer Development and Progression. Front Oncol. 2018;8:225. Epub 2018/06/28. doi: 10.3389/fonc.2018.00225. PubMed PMID: 29946533; PMCID: PMC6005831.

54. Ma B, Ran R, Liao H-Y, Zhang H-H. The paradoxical role of matrix metalloproteinase-11 in cancer. Biomedicine & Pharmacotherapy. 2021;141:111899. doi: https://doi.org/10.1016/j.biopha.2021.111899.

55. Wu S, Chen M, Huang J, Zhang F, Lv Z, Jia Y, Cui Y-Z, Sun L-Z, Wang Y, Tang Y, Verhoeft KR, Li Y, Qin Y, Lin X, Guan X-Y, Lam K-O. ORAI2 Promotes Gastric Cancer Tumorigenicity and Metastasis through PI3K/Akt Signaling and MAPK-Dependent Focal Adhesion Disassembly. Cancer Research. 2021;81(4):986. doi: 10.1158/0008-5472.CAN-20-0049.

56. Holland SJ, Powell MJ, Franci C, Chan EW, Friera AM, Atchison RE, McLaughlin J, Swift SE, Pali ES, Yam G, Wong S, Lasaga J, Shen MR, Yu S, Xu W, Hitoshi Y, Bogenberger J, Nör JE, Payan DG, Lorens JB. Multiple roles for the receptor tyrosine kinase axl in tumor formation. Cancer Res. 2005;65(20):9294–303. Epub 2005/10/19. doi: 10.1158/0008-5472.Can-05-0993. PubMed PMID: 16230391.

57. Zecchini S, Bombardelli L, Decio A, Bianchi M, Mazzarol G, Sanguineti F, Aletti G, Maddaluno L, Berezin V, Bock E, Casadio C, Viale G, Colombo N, Giavazzi R, Cavallaro U. The adhesion molecule NCAM promotes ovarian cancer progression via FGFR signalling. EMBO Molecular Medicine. 2011;3(8):480–94. doi: https://doi.org/10.1002/emmm.201100152.

58. Black SA, Nelson AC, Gurule NJ, Futscher BW, Lyons TR. Semaphorin 7a exerts pleiotropic effects to promote breast tumor progression. Oncogene. 2016;35(39):5170–8. Epub 2016/04/12. doi: 10.1038/onc.2016.49. PubMed PMID: 27065336; PMCID: PMC5720143.

59. Adorno-Cruz V, Hoffmann AD, Liu X, Dashzeveg NK, Taftaf R, Wray B, Keri RA, Liu H. ITGA2 promotes expression of ACLY and CCND1 in enhancing breast cancer stemness and metastasis. Genes & Diseases. 2021;8(4):493–508. doi: https://doi.org/10.1016/j.gendis.2020.01.015.

60. Sayan AE, Stanford R, Vickery R, Grigorenko E, Diesch J, Kulbicki K, Edwards R, Pal R, Greaves P, Jariel-Encontre I, Piechaczyk M, Kriajevska M, Mellon JK, Dhillon AS, Tulchinsky E. Fra-1 controls motility of bladder cancer cells via transcriptional upregulation of the receptor tyrosine kinase AXL. Oncogene. 2012;31(12):1493–503. Epub 2011/08/09. doi: 10.1038/onc.2011.336. PubMed PMID: 21822309.

61. Cancer Genome Atlas Research N. The Molecular Taxonomy of Primary Prostate Cancer. Cell. 2015;163(4):1011–25. doi: 10.1016/j.cell.2015.10.025. PubMed PMID: 26544944; PMCID: 4695400.

62. Chen F, Zhang Y, Parra E, Rodriguez J, Behrens C, Akbani R, Lu Y, Kurie JM, Gibbons DL, Mills GB, Wistuba II, Creighton CJ. Multiplatform-based molecular subtypes of non-small-cell lung cancer. Oncogene. 2017;36(10):1384–93. doi: 10.1038/onc.2016.303.

63. McConkey DJ, Choi W. Molecular Subtypes of Bladder Cancer. Current Oncology Reports. 2018;20(10):77. doi: 10.1007/s11912-018-0727-5.

64. Collisson EA, Bailey P, Chang DK, Biankin AV. Molecular subtypes of pancreatic cancer. Nature Reviews Gastroenterology & Hepatology. 2019;16(4):207–20. doi: 10.1038/s41575-019-0109-y.

65. Katsuno Y, Derynck R. Epithelial plasticity, epithelial-mesenchymal transition, and the TGF-β family. Developmental Cell. 2021;56(6):726–46. doi: https://doi.org/10.1016/j.devcel.2021.02.028.

66. van Dijk D, Sharma R, Nainys J, Yim K, Kathail P, Carr AJ, Burdziak C, Moon KR, Chaffer CL, Pattabiraman D, Bierie B, Mazutis L, Wolf G, Krishnaswamy S, Pe’er D. Recovering Gene Interactions from Single-Cell Data Using Data Diffusion. Cell. 2018;174(3):716–29.e27. doi: https://doi.org/10.1016/j.cell.2018.05.061.

67. Lambert AW, Weinberg RA. Linking EMT programmes to normal and neoplastic epithelial stem cells. Nat Rev Cancer. 2021;21(5):325–38. Epub 2021/02/07. doi: 10.1038/s41568-021-00332-6. PubMed PMID: 33547455.

68. Chakrabarti R, Hwang J, Andres Blanco M, Wei Y, Lukačišin M, Romano RA, Smalley K, Liu S, Yang Q, Ibrahim T, Mercatali L, Amadori D, Haffty BG, Sinha S, Kang Y. Elf5 inhibits the epithelial-mesenchymal transition in mammary gland development and breast cancer metastasis by transcriptionally repressing Snail2. Nat Cell Biol. 2012;14(11):1212–22. Epub 2012/10/23. doi: 10.1038/ncb2607. PubMed PMID: 23086238; PMCID: PMC3500637.

69. Lehmann W, Mossmann D, Kleemann J, Mock K, Meisinger C, Brummer T, Herr R, Brabletz S, Stemmler MP, Brabletz T. ZEB1 turns into a transcriptional activator by interacting with YAP1 in aggressive cancer types. Nature Communications. 2016;7(1):10498. doi: 10.1038/ncomms10498.

70. Doehn U, Hauge C, Frank SR, Jensen CJ, Duda K, Nielsen JV, Cohen MS, Johansen JV, Winther BR, Lund LR, Winther O, Taunton J, Hansen SH, Frödin M. RSK Is a Principal Effector of the RAS-ERK Pathway for Eliciting a Coordinate Promotile/Invasive Gene Program and Phenotype in Epithelial Cells. Molecular Cell. 2009;35(4):511–22. doi: https://doi.org/10.1016/j.molcel.2009.08.002.

71. Shin S, Dimitri CA, Yoon S-O, Dowdle W, Blenis J. ERK2 but Not ERK1 Induces Epithelial-to-Mesenchymal Transformation via DEF Motif-Dependent Signaling Events. Molecular Cell. 2010;38(1):114–27. doi: https://doi.org/10.1016/j.molcel.2010.02.020.

72. Lee MK, Pardoux C, Hall MC, Lee PS, Warburton D, Qing J, Smith SM, Derynck R. TGF-β activates Erk MAP kinase signalling through direct phosphorylation of ShcA. The EMBO Journal. 2007;26(17):3957–67. doi: https://doi.org/10.1038/sj.emboj.7601818.

73. Zhao Y, Ma J, Fan Y, Wang Z, Tian R, Ji W, Zhang F, Niu R. TGF-β transactivates EGFR and facilitates breast cancer migration and invasion through canonical Smad3 and ERK/Sp1 signaling pathways. Mol Oncol. 2018;12(3):305–21. Epub 2017/12/08. doi: 10.1002/1878-0261.12162. PubMed PMID: 29215776; PMCID: PMC5830653.

74. Blanchette F, Rivard N, Rudd P, Grondin F, Attisano L, Dubois CM. Cross-talk between the p42/p44 MAP Kinase and Smad Pathways in Transforming Growth Factor &#x3b2;1-induced Furin Gene Transactivation *. Journal of Biological Chemistry. 2001;276(36):33986–94. doi: 10.1074/jbc.M100093200.

75. Hough C, Radu M, Doré JJE. TGF-Beta Induced Erk Phosphorylation of Smad Linker Region Regulates Smad Signaling. PLOS ONE. 2012;7(8):e42513. doi: 10.1371/journal.pone.0042513.

76. Pearson GW, Hunter T. Real-time imaging reveals that noninvasive mammary epithelial acini can contain motile cells. J Cell Biol. 2007;179(7):1555–67. Epub 2008/01/02. doi: 10.1083/jcb.200706099. PubMed PMID: 18166657; PMCID: PMC2373504.

77. Dang TT, Prechtl AM, Pearson GW. Breast cancer subtype-specific interactions with the microenvironment dictate mechanisms of invasion. Cancer Res. 2011;71(21):6857–66. Epub 2011/09/13. doi: 10.1158/0008-5472.Can-11-1818. PubMed PMID: 21908556; PMCID: PMC3206184.

78. Wrenn ED, Yamamoto A, Moore BM, Huang Y, McBirney M, Thomas AJ, Greenwood E, Rabena YF, Rahbar H, Partridge SC, Cheung KJ. Regulation of Collective Metastasis by Nanolumenal Signaling. Cell. 2020;183(2):395–410.e19. doi: https://doi.org/10.1016/j.cell.2020.08.045.

79. Massagué J. TGFβ signalling in context. Nat Rev Mol Cell Biol. 2012;13(10):616–30. Epub 2012/09/21. doi: 10.1038/nrm3434. PubMed PMID: 22992590; PMCID: PMC4027049.

80. David CJ, Massagué J. Contextual determinants of TGFβ action in development, immunity and cancer. Nat Rev Mol Cell Biol. 2018;19(7):419–35. Epub 2018/04/13. doi: 10.1038/s41580-018-0007-0. PubMed PMID: 29643418; PMCID: PMC7457231.

81. Clement JH, Raida M, Sänger J, Bicknell R, Liu J, Naumann A, Geyer A, Waldau A, Hortschansky P, Schmidt A, Höffken K, Wölft S, Harris AL. Bone morphogenetic protein 2 (BMP-2) induces in vitro invasion and in vivo hormone independent growth of breast carcinoma cells. Int J Oncol. 2005;27(2):401–7. Epub 2005/07/13. PubMed PMID: 16010421.

82. Liberzon A, Birger C, Thorvaldsdóttir H, Ghandi M, Mesirov JP, Tamayo P. The Molecular Signatures Database (MSigDB) hallmark gene set collection. Cell Syst. 2015;1(6):417–25. doi: 10.1016/j.cels.2015.12.004. PubMed PMID: 26771021.

83. Huang P, Chen A, He W, Li Z, Zhang G, Liu Z, Liu G, Liu X, He S, Xiao G, Huang F, Stenvang J, Brünner N, Hong A, Wang J. BMP-2 induces EMT and breast cancer stemness through Rb and CD44. Cell Death Discov. 2017;3:17039. Epub 2017/07/21. doi: 10.1038/cddiscovery.2017.39. PubMed PMID: 28725489; PMCID: PMC5511860.

84. Cheng YX, Xiao L, Yang YL, Liu XD, Zhou XR, Bu ZF, Cao PC, Wang DK. Collagen type VIII alpha 2 chain (COL8A2), an important component of the basement membrane of the corneal endothelium, facilitates the malignant development of glioblastoma cells via inducing EMT. J Bioenerg Biomembr. 2021;53(1):49–59. Epub 2021/01/07. doi: 10.1007/s10863-020-09865-1. PubMed PMID: 33405048.

85. Zhao J, Ou B, Han D, Wang P, Zong Y, Zhu C, Liu D, Zheng M, Sun J, Feng H, Lu A. Tumor-derived CXCL5 promotes human colorectal cancer metastasis through activation of the ERK/Elk-1/Snail and AKT/GSK3β/β-catenin pathways. Mol Cancer. 2017;16(1):70. Epub 2017/03/31. doi: 10.1186/s12943-017-0629-4. PubMed PMID: 28356111; PMCID: PMC5372323.

86. Susek KH, Karvouni M, Alici E, Lundqvist A. The Role of CXC Chemokine Receptors 1-4 on Immune Cells in the Tumor Microenvironment. Front Immunol. 2018;9:2159. Epub 2018/10/16. doi: 10.3389/fimmu.2018.02159. PubMed PMID: 30319622; PMCID: PMC6167945.

87. Roberts PJ, Usary JE, Darr DB, Dillon PM, Pfefferle AD, Whittle MC, Duncan JS, Johnson SM, Combest AJ, Jin J, Zamboni WC, Johnson GL, Perou CM, Sharpless NE. Combined PI3K/mTOR and MEK inhibition provides broad antitumor activity in faithful murine cancer models. Clin Cancer Res. 2012;18(19):5290–303. doi: 10.1158/1078-0432.CCR-12-0563. PubMed PMID: 22872574; PMCID: 3715399.

88. Usary J, Zhao W, Darr D, Roberts PJ, Liu M, Balletta L, Karginova O, Jordan J, Combest A, Bridges A, Prat A, Cheang MC, Herschkowitz JI, Rosen JM, Zamboni W, Sharpless NE, Perou CM. Predicting drug responsiveness in human cancers using genetically engineered mice. Clin Cancer Res. 2013;19(17):4889–99. doi: 10.1158/1078-0432.CCR-13-0522. PubMed PMID: 23780888; PMCID: 3778918.

89. Reguart N, Remon J. Common EGFR-mutated subgroups (Del19/L858R) in advanced non-small-cell lung cancer: chasing better outcomes with tyrosine kinase inhibitors. Future Oncology. 2015;11(8):1245–57. doi: 10.2217/fon.15.15. PubMed PMID: 25629371.

90. Li S, Liu M, Do MH, Chou C, Stamatiades EG, Nixon BG, Shi W, Zhang X, Li P, Gao S, Capistrano KJ, Xu H, Cheung N-KV, Li MO. Cancer immunotherapy via targeted TGF-β signalling blockade in TH cells. Nature. 2020;587(7832):121–5. doi: 10.1038/s41586-020-2850-3.

91. Kanda T, Sullivan KF, Wahl GM. Histone-GFP fusion protein enables sensitive analysis of chromosome dynamics in living mammalian cells. Curr Biol. 1998;8(7):377–85. Epub 1998/05/16. doi: 10.1016/s0960-9822(98)70156-3. PubMed PMID: 9545195.

92. Benjamini Y, Hochberg Y. Controlling the False Discovery Rate: A Practical and Powerful Approach to Multiple Testing. Journal of the Royal Statistical Society: Series B (Methodological). 1995;57(1):289–300. doi: https://doi.org/10.1111/j.2517-6161.1995.tb02031.x.

93. Love MI, Huber W, Anders S. Moderated estimation of fold change and dispersion for RNA-seq data with DESeq2. Genome Biology. 2014;15(12):550. doi: 10.1186/s13059-014-0550-8.

94. Rauch A, Haakonsson AK, Madsen JGS, Larsen M, Forss I, Madsen MR, Van Hauwaert EL, Wiwie C, Jespersen NZ, Tencerova M, Nielsen R, Larsen BD, Röttger R, Baumbach J, Scheele C, Kassem M, Mandrup S. Osteogenesis depends on commissioning of a network of stem cell transcription factors that act as repressors of adipogenesis. Nature Genetics. 2019;51(4):716–27. doi: 10.1038/s41588-019-0359-1.

95. Yu G, Wang L-G, Han Y, He Q-Y. clusterProfiler: an R Package for Comparing Biological Themes Among Gene Clusters. OMICS: A Journal of Integrative Biology. 2012;16(5):284–7. doi: 10.1089/omi.2011.0118.

96. Fernandez AI, Geng X, Chaldekas K, Harris B, Duttargi A, Berry VL, Berry DL, Mahajan A, Cavalli LR, Győrffy B, Tan M, Riggins RB. The orphan nuclear receptor estrogen-related receptor beta (ERRβ) in triple-negative breast cancer. Breast Cancer Research and Treatment. 2020;179(3):585–604. doi: 10.1007/s10549-019-05485-5.

97. Cerami E, Gao J, Dogrusoz U, Gross BE, Sumer SO, Aksoy BA, Jacobsen A, Byrne CJ, Heuer ML, Larsson E, Antipin Y, Reva B, Goldberg AP, Sander C, Schultz N. The cBio cancer genomics portal: an open platform for exploring multidimensional cancer genomics data. Cancer Discov. 2012;2(5):401–4. doi: 10.1158/2159-8290.CD-12-0095. PubMed PMID: 22588877; PMCID: 3956037.

98. Liu J, Lichtenberg T, Hoadley KA, Poisson LM, Lazar AJ, Cherniack AD, Kovatich AJ, Benz CC, Levine DA, Lee AV, Omberg L, Wolf DM, Shriver CD, Thorsson V, Hu H. An Integrated TCGA Pan-Cancer Clinical Data Resource to Drive High-Quality Survival Outcome Analytics. Cell. 2018;173(2):400–16.e11. Epub 2018/04/07. doi: 10.1016/j.cell.2018.02.052. PubMed PMID: 29625055; PMCID: PMC6066282.

